# Complexity Begets Simplicity: Self-Supervised Learning for Palaeontological Images with Few or No Labels

**DOI:** 10.1101/2025.11.13.688022

**Authors:** Niall Rodgers

## Abstract

Palaeontology has seen widespread and growing use of machine learning to classify and analyse large datasets of fossils. However, palaeontology is a challenging field in which to apply machine learning. Datasets may be small or unlabelled, images may be complex and different from standard datasets and palaeontologists may lack specialist training and access to necessary computational resources. We show how these challenges can be addressed by utilising recent developments in self-supervised learning (SSL). Using a frozen DINOv3 feature extractor and a simple linear classifier, with reduced data, we can achieve comparable results to literature benchmarks using Convolutional Neural Networks (CNNs), the previous standard, when classifying fossil tracks, pollen, radiolaria, foraminifera and a dataset of diverse fossil images. Additionally, the rich feature vectors generated by the model can be used for few-shot learning, unsupervised clustering and quantification of disparity. Using state-of-the-art self-supervised methods increases accessibility by reducing code, compute and data required. It also maintains accuracy, while increasing reproducibility by reducing parameters and allowing simple future-proof model agnostic pipelines which may become the new standard approach in palaeontology.

## 1 Introduction

Machine learning (ML) is currently having profound impacts on society and science; however, the most recent developments in ML methodology have yet to break into palaeontology [1, 2, 3]. Palaeontology is a challenging domain for ML for a variety of reasons. Datasets can be small, and labelling is time-consuming and require specialist knowledge. Images are complex with potentially degraded objects, distinct from standard ML training data, which are difficult to segment from the background and each other. Palaeontologists may not have the mathematical and computational training found in some disciplines of the natural sciences, making applying complex ML pipelines of untested effectiveness unappealing. Moreover, due to the number of parameters and complexity of these pipelines, there are issues with reproducibility in many ML studies in palaeontology [3]. Ideally, palaeontologists could learn a single pipeline and apply it to diverse datasets with minimal augmentations and modification of parameters.

Recent developments, however, present the possibility to partially ameliorate all of the above issues. State-of-the-art models [4] pre-trained on vast amounts of unlabelled data using the approach known as self-supervised learning [4, 5, 6, 7, 8] allow robust rich feature vectors to be extracted from diverse complex images, in just a few lines of high-level Python code [9]. Using these frozen, off-the-shelf, models to extract features, it is then possible to obtain excellent performance in downstream tasks [4, 6]. For palaeontology, this raises the possibility that complex convolutional neural network pipelines could be replaced, in the first instance, with self-supervised feature extractors. The embedding quality also raises the possibility of using unsupervised learning to reveal structure in the data or use feature vectors properties as porxy for morphological disparity.

In this work, we apply the state-of-the-art self-supervised feature extractor DINOv3 [4] to various image sets across scales related to palaeontology, including foraminifera [10, 11], radiolaria [12], fossil tracks [13], pollen [14] and a dataset of diverse fossil images [15]. We show that we can accurately classify objects, with reduced data, simply using the extracted features and simple k nearest neighbour (kNN) or logistic regression classification, section 5. Furthermore, we show that applying non-linear dimensionality reduction to the feature vectors can reveal the structure of the datasets and that unsupervised learning can find meaningful groupings in the data without labels, section 6. Moreover, we propose that, under carefully controlled circumstances, the statistical properties of the set of extracted features may be used to understand the disparity of a dataset, section 8. This pipeline provides a generalisable framework that maintains accuracy, with less data, while reducing computational burden, length of code, and number of parameters. This framework could improve adoption of ML in palaeontology while increasing reproducibility and the range of useable datasets.

## 2 Literature Review

### 2.1 Machine Learning in Palaeontology

Reflective of its widespread adoption in society, ML has also been widely used in palaeontology, reviewed in detail in [1, 2, 3]. ML has been applied using images of many objects relevant to palaeontology [1, 2, 3], with examples including foraminifera [10, 11, 16], pollen [17, 14, 18], fossil tracks [13], radiolaria [19, 12], bivalves and brachiopods [20], graptolite [21] and cut marks on bone [22]. Computer vision in palaeontology has generally been dominated by Convolutional Neural Networks (CNNs) using transfer learning [1, 2, 3]. However, some studies have begun to use more recent frameworks such as: the application of metric learning to foraminifera [16], vision transformers to radiolaria [19] and the application of generative adversarial networks to pollen [18].

Despite the success of CNN-based approaches there are still limitations. A significant amount of labelled training data is still required, even for transfer learning approaches, and ideally GPU access to fine-tune the model. There are many choices that must be made and hyperparameters selected when building the CNN pipeline, such as the base model to use [3] and the data augmentation methods to employ, which can affect the performance of the pipeline [23].

Therefore, there still exists a barrier-to-entry in applying ML to a novel palaeontological dataset as enough data needs to be gathered to verify it works and the researcher has to invest time in fine-tuning the model. Recent developments such as foundation models based on self-supervised learning, which offer to partially resolve some of the above limitations, have yet to be applied to palaeontological data.

### 2.2 Self-Supervised Learning and the DINO Models

Self-supervised learning (SSL) is a methodology used to construct representations of unlabelled data such that they can be useful for downstream tasks [7, 8], which has reshaped the modern ML landscape. SSL has been a key component of the rapid advances in natural language processing beginning with models like BERT [24] and leading to the GPT family [25], allowing models to be trained on the vast amount of text available on the internet [7, 8]. Many implementations involve predicting masked parts of the data, for example prediction of words or sentences from the context or filling in the masked parts of an image [7, 8]. SSL approaches using contrastive learning frameworks [26] compete with traditional supervised frameworks in benchmark image classification tasks.

#### 2.2.1 DINO Models

The DINO (self-**di**stillation with **no** labels) approach to training vision transformers for feature extraction without labels was first proposed in 2021 [5]. This was then enhanced and iterated upon in DINOv2 and variants [6, 27] with the most up-to-date implementation being DINOv3 [4]. The DINO models are freely available (subject to the META AI custom licensing agreement) and were built by Meta AI, leveraging their access to large amounts of data and computational resources.

All DINO models build on the same core concept, the coupled training of a student and teacher network [5]. Later variants added the ability to scale to larger datasets and other modifications to improve the stability of training and embedding quality [6, 27, 4]. In order to achieve the goal of creating rich feature vectors, the training procedure is as follows: a global view of an image is passed to the teacher network, which produces an output vector; different augmented views of the same image are passed to the student network, which produces output vectors; a loss function is computed which measures the discrepancy in the output vectors between the teacher and student; parameters are updated to minimise this loss function and the teacher network is updated using an exponentially moving average of the student network. This training process coupled with large amounts of data is what allows the models to produce useful feature vectors even on unseen complex objects.

In this work, we focus on DINOv3, the state-of-the-art variant at the time of writing, which produces representations that may share similarities with those in the human brain [28]. DINOv3 was trained in two configurations by MetaAI, one using satellite image data and another using web data [4]. We focus on the web data variant as this is more appropriate for our datasets. To begin training DINOv3 roughly 17 billion images were collected from Instagram, this dataset was then subsampled to create a balanced dataset of approximately 1.7 billion images used to train the model [4]. The complete DINOv3 model has around 7 billion parameters, so to reduce computational burden the model was distilled into a range of smaller variants, which approximate its behaviour [4].

The DINO training framework can compete with (or exceed) the results of supervised convolutional neural networks in image classification tasks [4, 6, 5] using only a simple linear or kNN classifier applied to the feature vectors. This represents a paradigm shift in ML as CNN-based supervised approaches have traditionally dominated ML for image classification [29, 30]. For domains such as palaeontology, the default way to deal with image classification has been to use transfer learning of standard CNNs pre-trained on ImageNet [1]. However, this standard approach can be called into question as the power of a frozen feature extractor alone can achieve the same results with better generalisation to unseen contexts as shown by the rapid uptake and success of the DINO model family.

At the time of writing DINOv3 [4] has been released too recently to have been widely applied in the academic literature, although initial results on medical images [31], anomaly [32] and AI-generated image detection shows promise [33]. However, previous iterations of the DINO family have been applied in diverse range of contexts, with the paper proposing DINOv2 [6] cited more than 4000 times on Google Scholar since 2023. Examples of applications where DINO features are used as part of the pipeline include but are not limited to: geological rock images [34], identification of plant species [35], individual animals [36], geophysical data [37], industrial anomaly detection [38], robotics [39, 40, 41] and medical imaging [42, 43, 44, 45]. The large number of citations and diverse range of applications imply a powerful generalisability, ideal for palaeontology.

## 3 Feature Extraction

The promise of this methodology is to make the conversion of images into rich feature as simple as possible. The cornerstone of this is the Transformers Python library produced by the company Hugging Face [9]. This library allows for the easy sharing of models, providing easy access to state-of-the-art ML. Additionally, it is very well documented, see example code and detailed description of the model used in this work at https://huggingface.co/facebook/dinov3-vitl16-pretrain-lvd1689m.

The library [9] allows for the automatic selection of the appropriate preprocessor for a given model. This allows us to directly pass images to the model irrespective of the image format or dimension, simplifying the preprocessing for non-specialists. We apply the selected preprocessor, which converts the image to an RGB tensor in a format appropriate for the PyTorch library [46] of size 224 × 224 rescaled so that pixels take values between 0 and 1 and normalised using the ImageNet mean [0.485, 0.456, 0.406] and standard deviation [0.229, 0.224, 0.225]. The model can then be automatically initiated using the Transformers library [9] and the model ID. Once loaded it can be applied to the preprocessed images and the output features collated.

From the DINOv3 model family [4], we select the large vision transformer trained on web data with approximately 0.3 billion parameters, distilled from the full DINOv3 model. The selected model converts images to 1024-dimensional feature vectors. This size of model balances inference speed and embedding quality for the purposes required in this work; smaller models could be used in settings where computational resources are more restricted, and larger models could be used where gains in accuracy are critical and resources are less constrained. Feature vectors can then be passed directly to the classification pipeline based on logistic regression and kNN (section 5.1), for unsupervised clustering (section 6.2) or used quantify the structure of the dataset (section 8).

## 4 Datasets

To test the proposed pipeline, we selected five datasets of varying size and class number relevant to various parts of palaeontology. Our criteria for dataset selection are that the data (or a representative subset) is freely accessible from a public repository. The datasets should also have been used for classification such that a headline accuracy score, the primary currency of papers in the area, is generated which can be reasonably compared to the robust cross-validated pipeline we present. There are many difficulties surrounding simultaneous exact reproductions of multiple ML pipelines in palaeontology, something also discussed in [3]. These include the differences in splits and cross-validations; discrepancies in reported number of images and number downloaded; variations in approaches to imbalanced data and accessibility of test sets.

As a result of this, we choose to compare our pipeline to the headline numbers presented in the literature rather than an exact reproduction of every step of every paper. This is justified by the fact that although the presented results show that we can match the headline accuracy, even if this was not the case, the methodology would still be valuable based on the data efficiency, accessibility and unsupervised learning potential. Thus we encourage researchers to implement this approach and test on their own data. Additionally, this is required for brevity and accessibility to reach the widest possible audience in palaeontology. This is one of the few works to simultaneously tackle ML across different areas of palaeontology using a single pipeline. The datasets, where all classification used CNNs, are described in detail as follows.

### 4.1 Pollen

Pollen grains generally preserve well and are important for our understanding of Earth’s history. Hence, to test the ability of this pipeline to classify pollen grains we select the dataset used in [14]. This dataset contains pollen grains from 46 classes (mainly species with some family classes spread over 37 families) from the New Zealand and Pacific region. The roughly 19,500 images are collapsed “z-stacks” of microscope images taken at different focal planes, artificially coloured to show depth.

The classification of this type of pollen can be done exceptionally well by automated means with [14] achieving 97.86% validation accuracy in their cross-validated pipeline. Ideally, the CNN pipeline would also contain a withheld test set to prevent data leakage, during early-stopping or any hyper-parameter tuning, however data and computational requirements can restrict this. In contrast, as DINO features and simple classifier do not require a validation set this reduces the opportunities for data leakage.

### 4.2 Radiolaria

To test the methodology on a smaller dataset with a number of classes amenable to interpretation for unsupervised classification we use the dataset of radiolaria assembled in [12]. The work of [12] splits the dataset into three sets, with one set labelled “S” being the full data and the others the same data expect for the removal of broken or blurred objects. This full dataset we use contains 1085 microscope image of radiolaria across 8 species from the genus *Podocyrtis*.

On the full dataset the CNN-based method of [12] achieved a validation accuracy of 88.46%. A dataset of this size can be processed extremely quickly by our pipeline.

### 4.3 Foraminifera

Foraminifera are a key group of organisms to which ML has been applied in palaeontology, such as in the creation of the Endless Forams dataset in [10], since many samples can be collected. ML is also appealing due to the possibility of pairing ML with automated systems to physically manipulate objects [47]. The large number of samples has allowed the implementations of methodologies that require large amounts of data, such as [48] which uses vision transformers and generative adversarial networks to generate synthetic data. The Endless Forams dataset and subsets of it have been used to test automated classification of foraminifera several times [10, 11, 16]. The total Endless Forams dataset contains approximately 34,000 images which are available online [10]. However, they are not downloadable as single folder so we take the subset of this data approximately 27,000 images split into 35 species classes which was uploaded to Zenodo [49] as part of [11].

Using the complete dataset and standard CNN [10] achieved a maximum accuracy of 87.4% on a crossvalidated pipeline. More specialist architectures which are trained from scratch on the approximately 27,000 image dataset (while removing small classes) managed to achieve a maximum accuracy of 90.3% [11], at the cost of increasing training time. Using higher resolution images and metric learning on the approximately 34,000 image dataset [16] managed to reach 92% accuracy, again this is at the cost of increasing training time, requiring 48 hours on a GPU [16]. The discrepancies in image numbers, how small classes are dealt with and evaluations between papers highlights the difficulties in reproduction of results in ML in palaeontology.

### 4.4 Fossil Tracks

In order to test the ability of the model to work with ambiguous objects that are simple outlines without colour or textural information we select the dataset of [13]. This iconology dataset consists of 693 ornithischian tracks and 980 theropod tracks which are simple binary images consisting of the track itself and no other contextual information. In the work of [13], the dataset is down-sampled to create a balanced dataset and then tested on a small fixed set of 36 images so that the test set can be passed to human experts. As we cannot directly access the split of the downsample or the test set we train on the full dataset, our pipeline offers the benefit that we train with the class balanced option when using the scikit-learn [50] implementation of logistic regression, which mitigates the imbalance without an additional resampling step.

In [13], human experts and the neural network achieve similar accuracy scores as some of the examples are fundamentally ambiguous. The work of [13] presents results with a binary classifier or with a third unsure option; however, for simplicity and consistency we focus on the binary case where their methodology achieves 86% accuracy on the fixed test set after a validation accuracy of 83% during the training process.

### 4.5 Diverse Fossil Images (RFID)

Finally, to test our methodology on diverse data we test our method on the images that were web-scraped in the work of [15]. This large dataset consists of 50 major fossil groupings covering invertebrates, vertebrates, plants and microfossils. As to be anticipated from data which is scraped from web data there are a large variations in image quality (and relevance), background and image type (isolated skeletal subsections, whole specimens or sketches).

In this work, we select the reduced but still large version of the dataset of approximately 60,000 images as this is more than enough data for our methodology to be effective. This data set hence referred to reduced fossil image dataset (RFID) has been downsampled to remove the imbalance present in the full dataset with each class containing approximately 1200 images. The authors of [15] manage to achieve a test top-1 accuracy of 82.97% on this dataset and also reported a top-3 accuracy of 92.57%.

## 5 Supervised Classification

### 5.1 Classification Pipeline

The goal of our classification pipeline is accurate and robust results while maximising accessibility and simplicity. Hence, we directly pass the feature vectors to two simple classifiers, logistic regression and k nearest-neighbours (kNN) from [50].

A kNN classifier simply takes an unclassified data point and finds the *k* nearest-neighbours, according to the given distance metric, then classifies the unknown point by taking the majority class of the *k* nearestneighbours. Logistic regression (or multinomial logistic regression in the case of more than two classes) uses linear boundaries in feature space. This is achieved by applying a sigmoid (binary) or softmax (multi-class) function to linear combinations of the variables to convert them to probabilities of belonging to a particular class.

When using the kNN classifier we use a cosine distance metric, *k* = 5 neighbours and *l*^2^ (euclidean) normalisation of the feature vector. When performing logistic regression we directly use the default settings of sklearn [50] aside from fixing a random seed for reproducibility, increasing the maximum iterations to 1000 and using the balanced class weight setting. This directly handles data imbalance by weighting the importance of examples inversely to the number in that class.

We do not claim that this pipeline is optimal. Improvements could be made by doing dimension reduction, using a different classifier or by hyper-parameter tuning. However, we explicitly choose the minimal simplest pipeline to demonstrate that DINOv3 feature vectors are rich and that literature benchmarks can be matched (or exceeded) with this approach. In certain situations, small gains in accuracy could be made by using a non-linear classifier. However, DINO feature vectors generally tend towards linear separability of classes so gains are likely to be marginal whilst risking over-fitting.

Another benefit of this approach is that because of the ease of training, we can adopt a rigorous repeatable cross-validation pipeline, ensuring each sample is used for both testing and training. When performing supervised learning, we take the complete dataset (defined as all the data to which we have access) and sample the following fractions [0.1, 0.3, 0.5, 0.7, 1], repeating this process 10 times for each fraction. For each sample, our aim is to perform 5-fold cross-validation. For small dataset fractions, this may prove impractical. If the minimum number of examples per class is *<* 5, then we simply perform an 80 : 20 train-test split, attempting stratification if possible, and repeat 5 times to approximate multiple folds.

Classification metrics are reported as the mean and standard deviation combined across folds and repeats. In the main text, we focus on accuracy to compare to the literature with tables listing precision, recall, f1 score, specificity, as well as balanced, top-2 and top-3 accuracy given in the supplementary information.

### 5.2 Matching Literature Benchmarks

**Figure 1:**
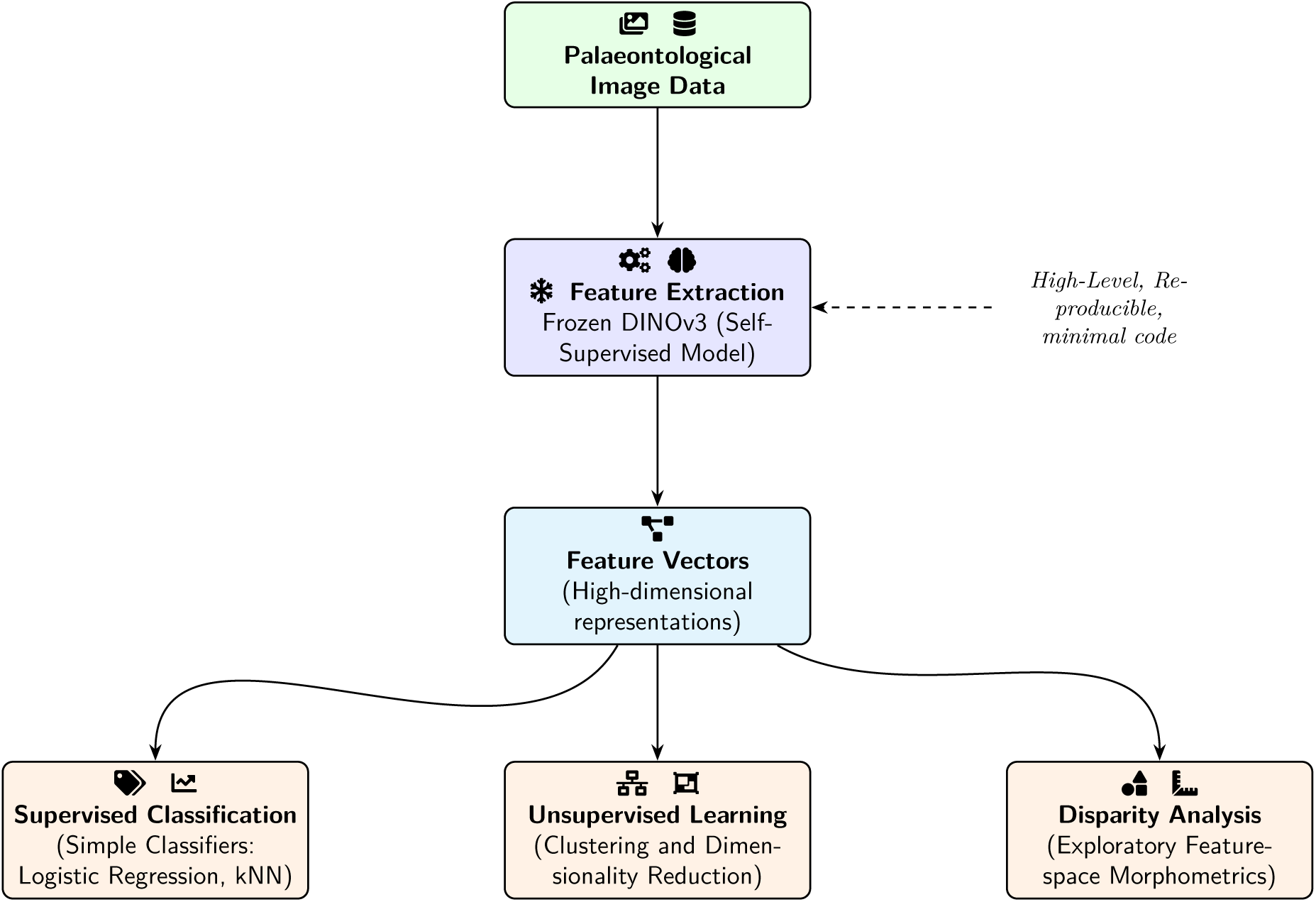
High-Level Overview of the pipeline proposed in this study.

In table 1, we compare the maximum accuracy score of our method to the headline accuracy score shown in the literature (the maximum accuracy given in the cited work). For all cases, our maximum accuracy occurs when applying logistic regression to the maximum amount of data. On the RFID, the kNN classifier is numerically very close to the logistic regression, as it is likely that the dataset clusters separately in feature space.

**Table 1:** Comparison of maximum classification pipeline accuracy to reported headline literature accuracy.

| Dataset | Maximum Accuracy [our protocol, self-supervised] (%) | Representative Headline Literature Accuracy (%) [CNN] |
| --- | --- | --- |
| Pollen | **98.34** | 97.86 [14] |
| Radiolaria | **94.30** | 88.46 [12] |
| Fossil Tracks | 81.28 | **86** (small test set) [13] |
| Diverse Fossils (RFID) | **83.31** | 82.97 [15] |
| Foraminifera | 86.95 | 87.4 [10], 90.3 [11], **92.0** [16] |

Our cross-validation pipeline exceeds the reported accuracy in pollen, radiolaria and the RFID. In pollen classification, our method is exceptionally accurate, as is the CNN approach. In radiolaria, we exceed the accuracy of the literature benchmark, although this may be due to choices made during the CNN pipeline design. We also perform well on the RFID with top-3 accuracy of 94%. We do not exceed but are around the same accuracy on the fossil tracks small test set and are close to the reported validation accuracy of 83%. This is not unsurprising, as the fossil track dataset contains many fundamentally ambiguous cases. DINOv3 is designed to extract features from complex RGB images, that it performs almost as well as fine-tuned CNN on simple binary shape outlines is notable. In foraminifera classification, our reported accuracy is very close to the accuracy of [10] who used a transfer learning approach. It is below the bespoke network trained in [11] and the metric learning approach [16]. This is again unsurprising, DINOv3 plus simple classifier matches the performance of transfer learning but can be beaten at the cost of coding a complex pipeline and the use of days of GPU time [16].

However, these results taken together demonstrate that given the trade-offs (or lack thereof) between accuracy and simplicity, it makes sense for DINOv3 plus a simpler classifier to replace CNN-based transfer learning, at least in the first instance. Then only in cases where accuracy is paramount and data volume allows should bespoke methods be considered. Cross-validated confusion matrices built using logistic regression applied to all available data can be found in the SI.

### 5.3 Reducing Dataset Size

Another advantage of the our pipeline is data efficiency. Across all datasets tested performance is reasonably good at only 10% of the data and the rate of increase begins to plateau at only 30% at the data used, figure 2. Results are competitive with the literature headline accuracies with only 30% of the data when classifying pollen, radiolaria and the diverse images. While, the rate of increase is slower in the case of foraminifera and the fossil tracks. Data efficiency is a key asset for palaeontology and may allow a wider array of data to be studied, lowering the barrier to entry.

**Figure 2:**
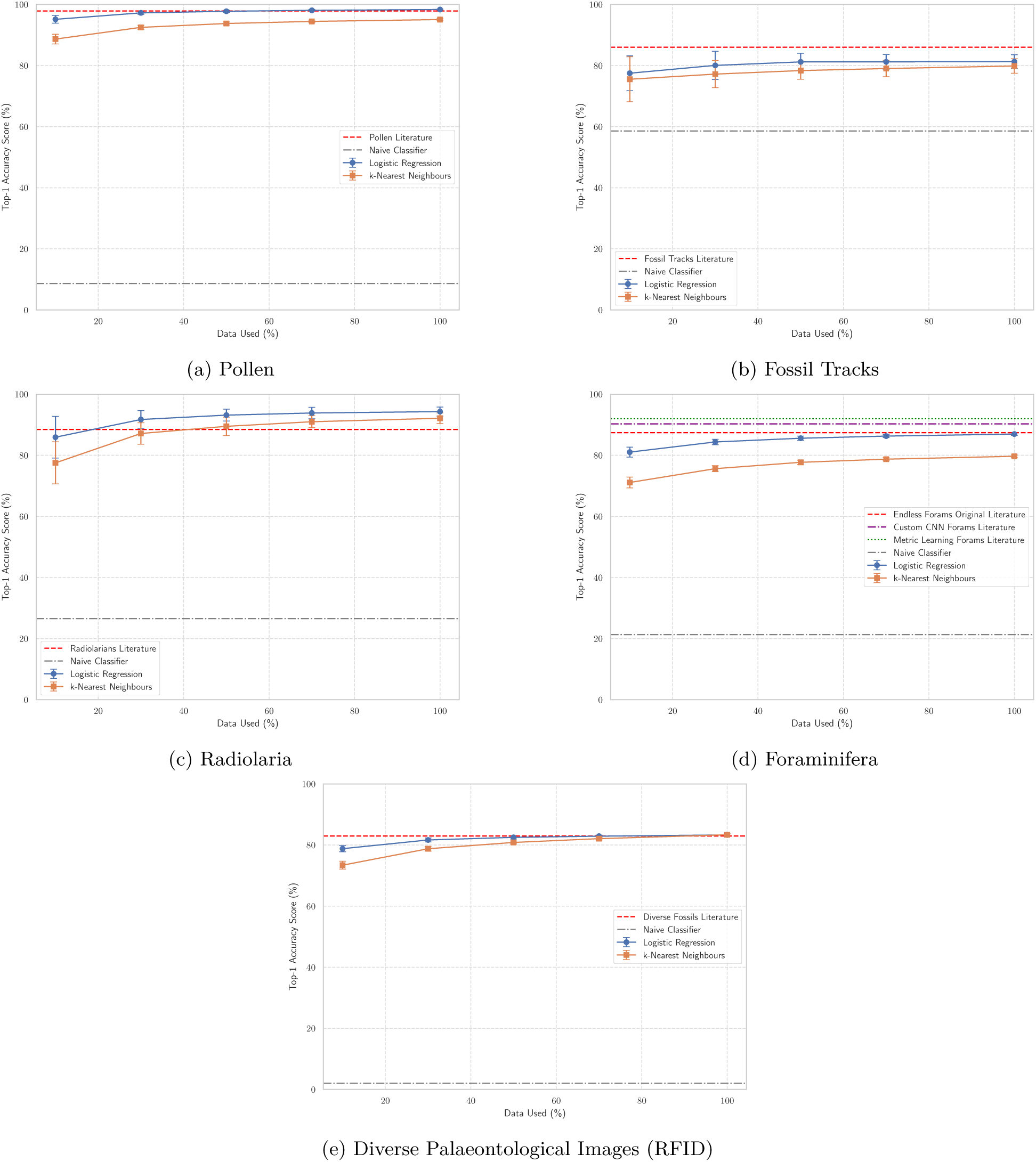
Changes in mean accuracy of our pipeline with fraction of data sampled, headline literature accuracy and a naive classifier accuracy (always predicting the most common class) are shown as horizontal lines. Error bars are standard deviations over folds and repeats.

### 5.4 Few-Shot Learning

Few-shot learning is the process whereby a ML algorithm learns from a small number of examples. Gaining insight from limited data is useful in both proof-of-concept tests and situations of data scarcity.

For few-shot learning, we use a similar classifier setup as in the previous section with the logistic regression unchanged while in the kNN approach we only change that number of neighbours is now set as either 5 or the maximum number of allowed training examples per few-shot run, whichever is smaller. We randomly select *n* training examples per class from the set [1, 2, 3, 5, 10, 20]. If *n* is higher than the number of examples in a class minus one, we select up to this point to leave at least one example in the test set. We then test on the remaining data and repeat this process 50 times.

All metrics, for a given number of training examples, are given in terms of mean and standard deviation over these repeats. In the main text, we again focus on the accuracy. Full metrics such as precision, recall, f1 score, specificity, as well as balanced, top-2 and top-3 accuracy are given in the SI.

Few-shot learning results are promising, figure 3. Something that would not be possible using a CNN-based pipeline, as it could not be fine-tuned using only a handful of examples. Pollen, radiolaria and the RFID are classified much more accurately than the naive classifier, selecting the most common class, after only a single training image. With performance steadily increasing towards the literature headline accuracy with the maximum number of examples. The fossil track images, which are both simple and ambiguous, take longer to see improvements above the naive classifier, however, this is unsurprising given this is an imbalanced binary classification problem. The foraminifera classifier shows slower growth with number of examples but again a reasonably accurate classifier can be achieved with only a small number of training examples. For both foraminifera and fossil tracks, it is likely that more examples are needed to learn the nuanced differences between classes. The tasks that performed best when the data volume was reduced, figure 2, also perform better for a smaller number of few-shot examples, showing that these are more easily distinguished in feature space.

**Figure 3:**
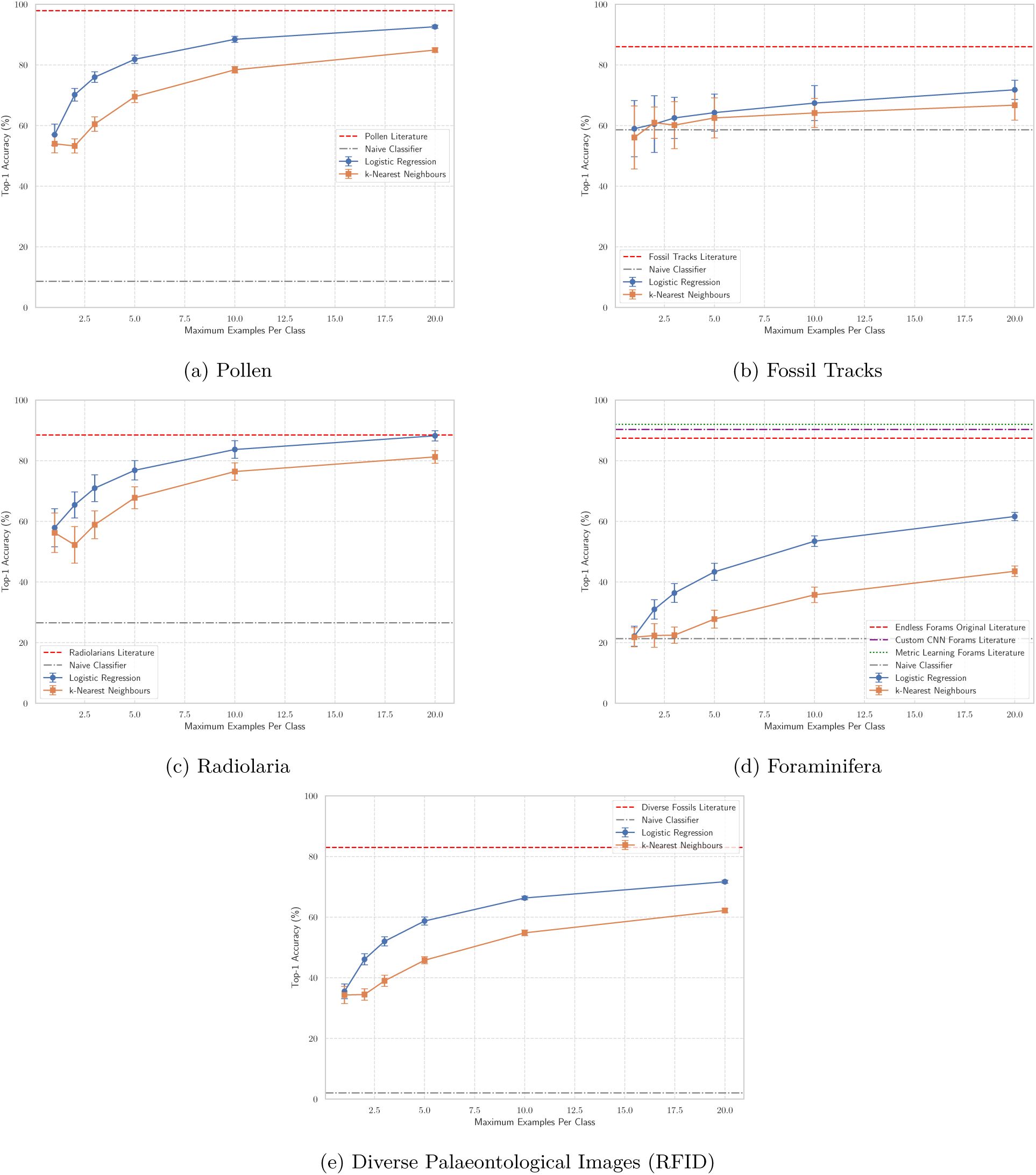
Changes in mean accuracy when doing few-shot learning for a given maximum number of examples per class, headline literature accuracy and a naive classifier accuracy (always predicting the most common class) are shown as horizontal lines. Error bars are standard deviation over repeats.

## 6 Unsupervised Learning

Unsupervised learning is the process of extracting information and groupings from unlabelled data. This is comparable to how a palaeontologist may sort fossils using morphology. In this section, we demonstrate how the rich feature vectors generated by DINOv3 can be used to find groupings and structure in data without labels. Unsupervised feature vector clustering has previously been used to group by morphology, such as in the work of [51] on mosquito classification. Firstly, we show how dimension reduction can be used to visualise the feature vectors and how the visualised structures align with the true labels. Then we apply unsupervised clustering methods to automate the detection of groupings in the data.

### 6.1 Visualisation of Feature Vectors

As the feature vectors generated by self-supervised models are high-dimensional objects, for visualisation purposes we reduce them to 2 dimensions. This can be done with linear methods such as Principal Component Analysis (PCA) or by using non-linear methods like UMAP [52, 53] or t-SNE [54] (accessed through [50]). Before applying dimension reduction methods, we *l*^2^ (euclidean distance) normalise all our vectors.

Unfortunately, see SI, PCA struggles to adequately represent these feature vectors in a small number of dimensions due to the large number of classes and how the models represent the data. As a result, we focus on non-linear dimensionality reduction methods; however, this reduces interpretability and accessibility. Interpretability decreases as distances are not preserved when doing non-linear dimension reduction. Accessibility is reduced as these methods are less common in palaeontology compared to PCA. t-distributed stochastic neighbour embedding (t-SNE) [54] is a non-linear method for visualising high dimensional data which converts data into a lower-dimensional space while trying to preserve the pairwise similarities between points. When using t-SNE for visualisation we use the scikit-learn implementation [50] setting a random seed, number of dimensions to 2, perplexity to 20, learning rate to 200, using a cosine distance metric and leaving the rest to the default settings. Uniform Manifold Approximation and Projection (UMAP) is another method for non-linear dimension reduction [52, 53]. UMAP attempts to find a low-dimensional representation of the data that preserves as much as possible the topological structure of the high-dimensional data [53]. UMAP is more computationally efficient than t-SNE and can preserve more global structure. When using UMAP for dimension reduction [52], we use a cosine distance metric, set the number of dimensions to 2, the minimum distance to 0.1, the number of neighbours to 30 and a random seed for reproducibility. For both UMAP and t-SNE parameters are chosen from within the standard ranges used in the literature to balance local and global structure. Both UMAP and t-SNE are stochastic and sensitive to parameter choices, so when used in practice, replotting and parameter testing is advised.

Figure 4, shows the application of t-SNE and UMAP to each set of feature vectors derived from the data used. In general, figure 4 demonstrates that DINOv3 feature vectors contain biologically meaningful information, as classes generally plot close to each other, in relatively well-defined groupings. This plot demonstrates why some of our datasets are easier to classify than others. Some datasets such as radiolaria, pollen and to an extent the RFID break into discrete groupings organised into classes. Whereas, others like the foraminifera and, in particular, the fossil tracks do not break into distinct groups in the same way. Additionally, when viewing the UMAP of the radiolaria, figure 4e, we can see three distinct groups that may relate to the three evolutionary lineages in the data [12].

**Figure 4:**
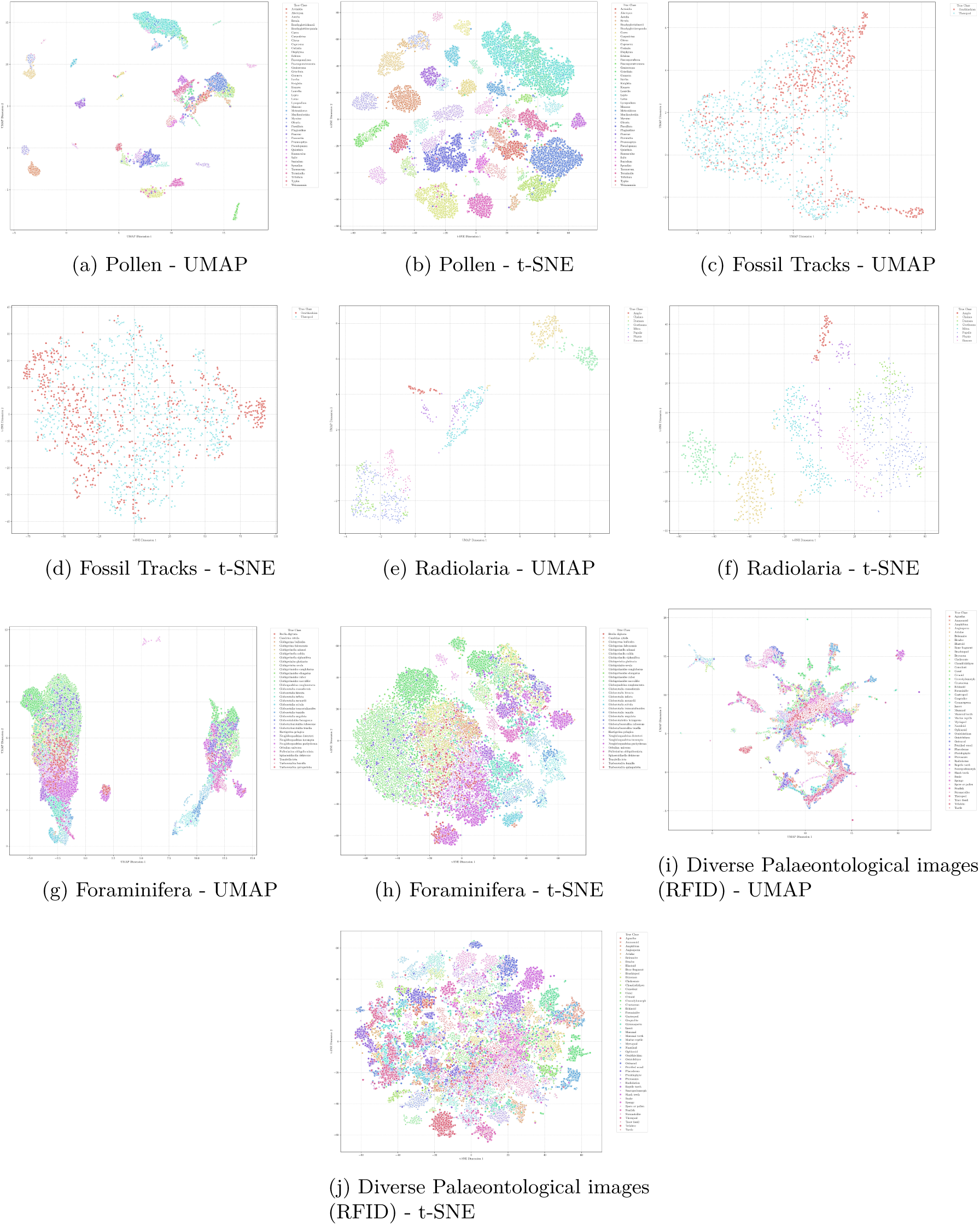
Non-linear dimensionality reduction methods applied to feature vectors for all images across all datasets, points labelled according to their true class. Larger version of images and colour label correspondence given in the SI, with original images in the code repository.

When working with feature vectors non-taxonomic groupings may appear due to patterns in the image data unrelated to biology. The fossil tracks dataset embeds into a single group highlighting the ambiguous region where it is difficult to categorise images. Meanwhile, the foraminifera groups into two large macroscopic clusters as a result of the fact that the dataset contains images of two distinct backgrounds and colour grades, so the feature vectors group objects of similar species independently in each of these two clusters. This may explain why we required more examples to classify the data well in section 3. This short demonstration, figure 4, highlights the utility of the dimensionality reduction as a data visualisation and inspection tool.

### 6.2 Clustering Methodology

To further test how biologically relevant information can be extracted from feature vectors, we tested several unsupervised clustering algorithms and compared the extracted clusters with biological labels. Before clustering, we reduce the dimensionality of the data, as unsupervised clustering algorithms based on distances can be computationally costly, and distances are less meaningful, in high dimensions. This is done using PCA and UMAP [52, 53] on *l*^2^ (euclidean distance) normalised feature vectors. When using PCA to reduce dimension for clustering, we reduce dimension to the minimum of 50 and the number of components needed to explain 95% of the variance. In practice, as the feature vectors are not conducive to linear dimension reduction, all datasets are reduced to 50 dimensions. UMAP is a powerful preprocessing step for clustering, but should be treated with care and used an exploratory method as the non-linear reduction in dimensionality does not preserve distances between points. When using UMAP for dimensionality reduction we step the target dimension to 15, the number of neighbours to 30, the distance metric to cosine, the minimum distance to 0 and a random seed for reproducibility.

We test 4 different methods for unsupervised (or partially unsupervised clustering). Firstly, we use partially unsupervised clustering, k-means clustering where the number of clusters expected, *k*, is known in advance. k-means finds clusters and assigns points such that the distances between points and the centroids of their assigned clusters are minimised. k-means runs very quickly and in some scenarios it may be realistic that there is some knowledge of how many clusters to expect.

The second method, used is k-means with an unknown number of clusters. We fit k-means for varying values of *k* between 2 and min(2 × #classes, 50). In practice, the range would be selected with an educated guess. The best value of *k* is selected using the silhouette score [55], via scikit-learn [50]. This compares the mean intra-cluster distance to the mean distance to the nearest cluster. This also justifies the range selection as silhouette score selection likely tends towards selecting a smaller number of larger clusters.

The third method used in the Bayesian Gaussian Mixture Model (BGMM), again from [50]. This method assumes that the data points in each cluster are drawn from a Gaussian distribution and automatically selects the probable number of clusters using Bayesian inference. We use the default setting from Scikit-learn [50], apart from setting the weight concentration prior to 0.1, a random seed and the number of components to min(2 × #classes, 50).

The final clustering method used is HDBSCAN [56], which performs density-based clustering across scales and aims to identify the most stable clusters. This algorithm automatically determines the number of clusters and assigns points away from dense regions as noise. We use the default settings of the implementation given by the package [56], aside from setting the minimum cluster size to be max(5, 0.02 × # samples) rounded to the nearest integer.

Unsupervised clustering is more difficult than supervised learning as it relies on the classes being well separated in feature space. Additionally, it is sensitive to parameter choice and objects may cluster at different scales. In practice, this would be dealt with by repeated tests with different parameters and algorithms, trying to determine which clusters are robust and what biological features they correspond to. As this work is a proof-of-concept and we simply focus on demonstrating how methodology works, we run the algorithms once and plot the resulting clusters on a visualisation UMAP, which reduces to 2 dimensions using number of neighbours equal to 30, minimum distance 0.1, a random seed and cosine metric.

In the main text, we only show the clusters from the exploratory UMAP pipeline, as they generally cluster much better than PCA as the DINOv3 embeddings have an intrinsically non-linear structure. In the supplementary information, we give detailed metrics for each clustering algorithm and dimensionality reduction method. Clustering methods can be evaluated by comparing the computed clusters with known labels or independently of the labels. For this, we use a variety of metrics from [50].

To compare the clusters with the labels, we use the Adjusted Rand Index (ARI) and Adjusted Mutual Information (AMI) to compare the similarity of the clusters with the true labels while adjusting for overlap from random clustering. We also measure the purity score, which indicates the fraction of points in each cluster which share the most common label in that cluster. To evaluate the clusters without reference to the underlying labels, we use the Silhouette Score, which compares intra-cluster distances to distances to other clusters, and the Davies–Bouldin index, which compares average intra-cluster spread to cluster separation, with lower values being better and indicating tighter, well-separated clusters.

### 6.3 Clustering Results

The results of clustering of the data are shown in figure 5. It is clear that different algorithms pick out different groupings are different scales, however, that meaningful groupings are found when comparing the plots to those by true class, figure 4. Full discussions of the quantitative clustering metrics for each dataset are given in the SI.

**Figure 5:**
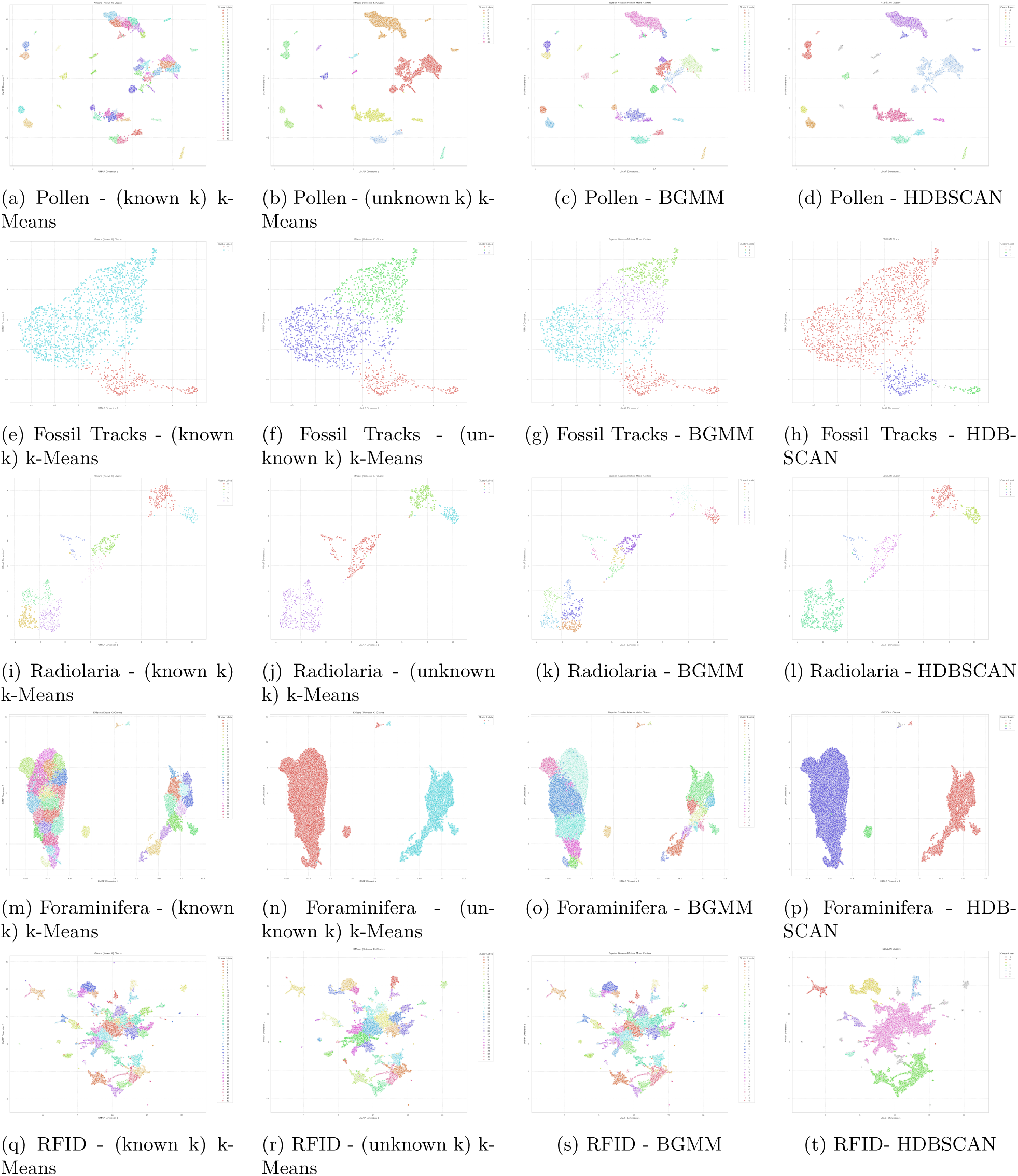
Comparison of Exploratory Unsupervised Clustering. Larger version of images and colour label correspondence given in the SI, with original images in the code repository.

Clustering of the pollen data is shown in figures 5a to 5b. Well-defined clusters are generally found; however, algorithms that infer the number of clusters generally cluster above the level of species. The fossil tracks, figures 5e to figures 5h, do not clearly separate in feature space. Hence, the algorithms break the objects into various subgroups. These represent genuine morphological variation, but do not precisely align with given labels. Clustering results the radiolaria data is generally successful, figures 5i to 5l, with clusters aligning with species or groups of similar species depending on the scale, with larger scale groupings aligning with the three evolutionary lineages in the data [12]. When clustering foraminifera, figures 5m to 5p, some of the algorithms focus on the differences in image backgrounds and colour scales, creating two large macroscopic clusters. Meanwhile, other approaches like the BGMM, figure 5o, find clusters which partially overlap with the known labels. Clustering the RFID, figure 5q to 5t, we again find generally well-defined clusters across different scales.

The results of figure 5 and supporting quantitative metrics in the supplementary information show that well-defined groupings can be extracted directly from palaeontological image data using DINOv3, unlocking analysis and interpretation of large unlabelled datasets.

## 7 Visualising Features Captured by Model

A useful step to confirm that the DINOv3 model is behaving as expected and focusing on relevant image regions is to visualise the intermediate outputs of the model. We present two ways to do this. Firstly, one can construct an attention map using the register tokens (additional learnable parts of the model) [27]. Secondly, one can take outputs at patch-level, reduce them to 3 dimensions, and convert to RGB values for visualisation, as demonstrated in [4].

In the first approach using register tokens, we pass an image through the preprocessor and the model, configured so that it also outputs the attentions. We then extract all the register tokens across all attention heads (parts of the transformer architecture). We take the median value per patch, to reduce noise and outliers. We then reshape this vector of median register attentions per patch into a square array for visualisation. For visualisation, we then apply a Gaussian blur (with sigma set to 2), clip extreme values (below the 5th and above the 95th percentile), normalise the values between 0 and 1 and resize to the same dimensions as the original image. The attention map can then be overlaid on the original image to give an indication of the parts of the image on which the model is focusing. This, however, is only indicative and not necessarily a true representation of how the model projects objects into feature space. The attention maps involve aggregation across heads, different tokens can give different representations and subsequent non-linearities are applied before the final output. Together, these aspects muddy the direct link between maps and explanation.

The second approach using patch-level RGB PCA works as follows. An image is passed through the preprocessor and model configured to output the hidden states, allowing us to extract the patch embeddings. The patch embeddings are then reduced to 3 dimensions using PCA, normalised between 0 and 1 and then reshaped into a square array per colour channel. To each colour channel, we apply a Gaussian blur (with sigma set to 1). Then we multiply each value by a colour multiplier (set to 2) and pass through a sigmoid function to increase colour vibrancy and allow for better differentiation between patches. We can then resize to match the original image and visualise as an RGB image.

This gives an indication of how the model separates parts of the image into different structures and how it views the image. Again, this comes with caveats: the PCA of the patch-level embeddings loses information; the layer this is applied to affects the representation; the sigmoid and steps to make the visualisation more appealing distort the raw differences in values between patches. However, this still gives a useful check to ensure that the model is segmenting the image in a sensible way.

When applying the feature visualisation methods to a selection of images from the datasets, figure 6, we find that results are positive. Showing that DINOv3 generalises well to the palaeontology domain. In the attention maps, attention is concentrated on the objects of interest. Furthermore, for the RGB PCA, the objects of interest are distinct and plotted in a different colour from the background. Even for the complex museum image, figures 6e and 6f, the attention map focuses directly on the object of interest while the PCA segments out the fossil.

**Figure 6:**
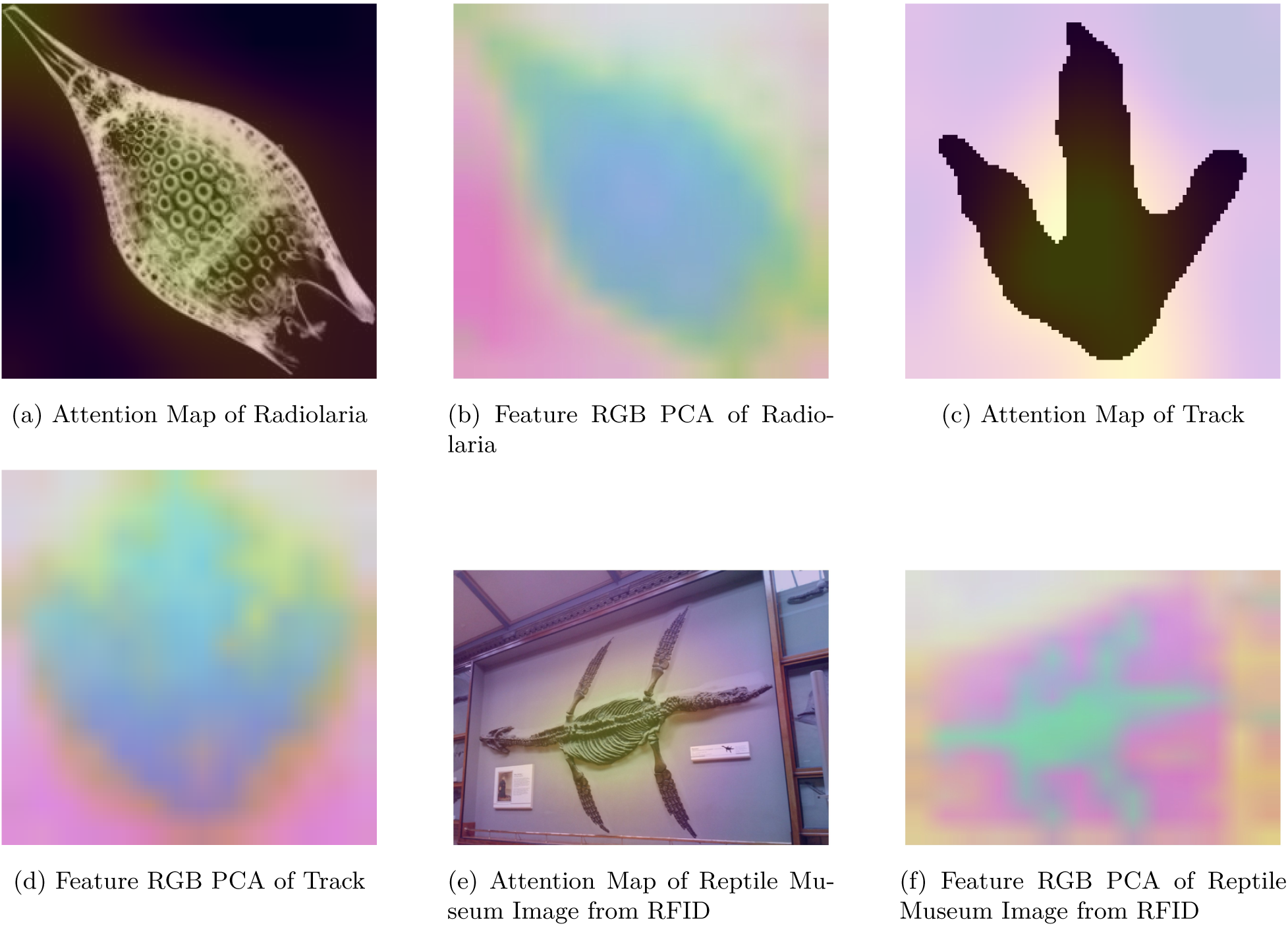
Comparison of techniques for visualising model understanding applied to a sample of images including Radiolaria *Podocyrtis Papalis* from [12], a described theropod footprint from [13] and an image from the marine reptile category in [15].

The results of figure 6 support our observation that these models perform well in the classification of palaeontological data and provide a methodology to interpret, as far as possible, the model performance on novel palaeontological datasets.

## 8 Image-Based Disparity Analysis

Given that by following this pipeline, we have access to the set of feature vectors which describe the objects. This raises the question as to whether these vectors can be used to quantify the morphological disparity of the objects they represent. This part of the work is more speculative in nature. However, this is justified for three reasons. Firstly, if you have followed the pipeline, the feature vectors are already computed and the disparity metrics are relatively simple to calculate, so there is very little cost associated with this method. Secondly, feature vector-based diversity measurements have previously been proposed for CNN feature vectors derived from studying pollen [57], where [57] used an entropy-based measure of diversity. Thirdly, this would represent a simple, fast and standardised way to capture some form of morphological disparity across a wide array of use cases.

As the primary purpose of this work is methodological and focuses on supervised learning and unsupervised clustering, we merely propose a set of metrics which can naturally be constructed from the feature vectors. If this method is employed in future work on other datasets with more metadata, it can be tested whether the prosed metrics and biologically informative variables correlate. The large caveat with this method is that care needs to be taken to ensure that the origin of the difference between vectors is due to the morphology of the objects and not due to differences in the background or imaging conditions, something that may need to be tested in detail on other datasets, as this is outside the scope of this work.

### 8.1 Methodology

A wide array of methods can be constructed to measure the spread, concentration and variation within a set of vectors. Hence, we propose four simple metrics, which are relatively simple to compute, relate to known concepts in palaeontology and are interpretable.

The first metric we propose is the mean and standard deviation of the pairwise distances, measured using a cosine distance metric. This metric reflects the overall morphological disparity by measuring how far apart objects are from each other in feature space.

The next proposed metric is the mean and standard deviations of the cosine distance from all samples to the centroid, the *l*^2^ normalised mean sample. This measures the spread of the objects and how much they deviate from the average object and spread across morphospace.

The next proposed metric is the variance sum, which is defined as

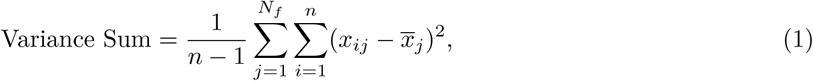

where *n* is the number of samples, *N_f_* is the number of features (1024 in our case) and *x̅_j_* is the mean value of a given feature vector entry. This measures how much the feature vector set spreads out over morphospace. Depending on the choice of metric and normalisation, the value of this, may be identical or directly proportional to the mean pairwise distance. As we use a cosine distance metric and *l*^2^ normalised feature vectors our measured mean pairwise distance is numerically identical to our variance sum. However, we include both measures as they prompt different conceptualisations.

The final metric proposed is the effective rank which is a relatively easy to compute information-theoretic measure of the effective dimensionality of the morphospace. The effective rank depends on the eigenvalues, λ*_i_* of the covariance matrix, estimated using the Ledoit–Wolf method to ensure stability for small sample numbers. The effective rank is defined as,

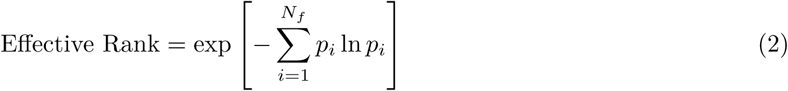

where

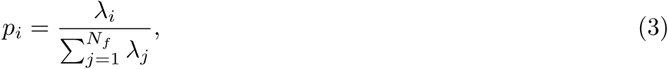

are the normalised eigenvalues of the covariance matrix. The effective rank is the exponentiation of the Shannon entropy of the normalised eigenvalue spectrum of the covariance matrix. When the probability density is concentrated in a single value (*p*_1_ = 1, ∀*i* ≠ 1, *p_i_* = 0) the effective rank is 1. When the probability is uniformly distributed 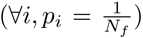 the effective rank is equal to *N_f_*. Hence, demonstrating why it acts like an effective dimensionality and gives an indication of the how filled the morphospace is. However, care should be taken when interpreting the effective rank for mixed datasets as between-group differences can lead to large eigenvalues which decrease the effective rank even though the disparity between samples has increased.

### 8.2 Results

The first way we can analyse the feature vector-based disparity is by inspecting the histograms of the pairwise distances between objects, figure 7. This shows that datasets which contain similar objects, such as the pollen, foraminifera and radiolaria, have generally smaller distances reflecting the fact that the vectors likely point to similar regions of feature space. For the RFID, the average distance is larger and nearer to one, reflecting the fact that the objects are diverse and there is much more variety in the direction of vectors.

**Figure 7:**
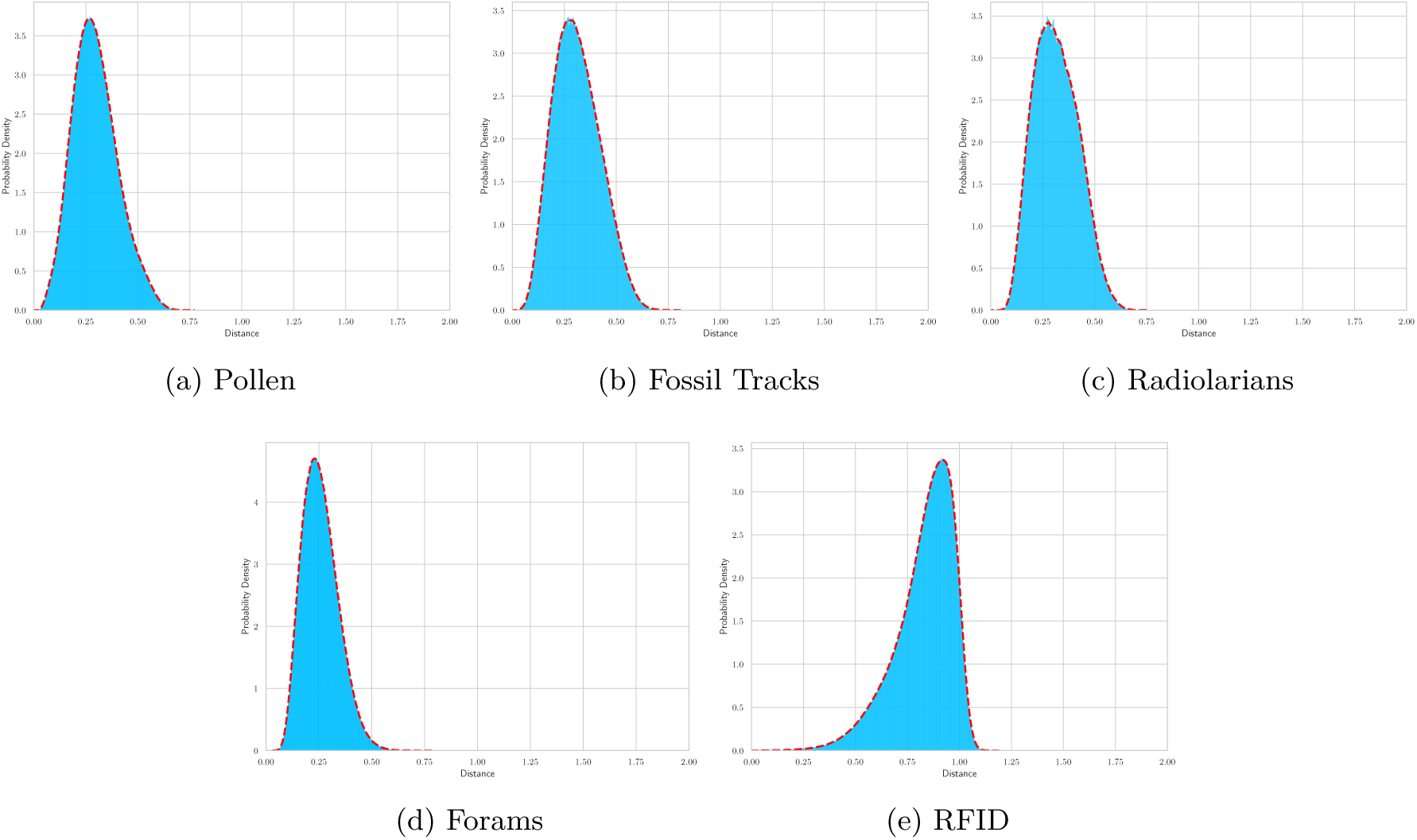
Histograms of Pairwise Distances (Cosine Distance between feature vectors)

The proposed disparity measurements are computed for the datasets in table 2. Table 2, shows that as expected on all counts of the diverse palaeontological images are much more disparate than the other datasets. It also shows that differences between other datasets are generally very subtle. Based on this, we would advise these metrics to be used with care, and only in situations where the morphological disparity is either directly comparable or the differences are large enough that the comparison is unambiguous. For example, it is quite fair to say that the diverse set of web image is more disparate than the a dataset of foraminifera or to compare two dataset of foraminifera with different distributions of species. However, it is ambiguous and potentially inadvisable to place stock in the fact that the dataset of radiolaria has a different disparity than a dataset of pollen based on minor differences. The results of table 2, show that the proposed metrics measure different aspects of the data as the ordering of the datasets is not preserved between metrics.

**Table 2:** Proposed Morphological Disparity Metrics Applied to Datasets used in this work.

| Dataset | No. of Samples | Mean Pairwise Dist. | std Pairwise Dist. | Mean Centroid Dist. | std Centroid Dist. | Variance Sum | Effective Rank |
| --- | --- | --- | --- | --- | --- | --- | --- |
| Pollen | 19667 | 0.2965 | 0.1090 | 0.1613 | 0.0608 | 0.2965 | 42.0603 |
| Fossil Tracks | 1671 | 0.3110 | 0.1116 | 0.1699 | 0.0661 | 0.3110 | 48.3432 |
| Radiolarians | 1085 | 0.3154 | 0.1061 | 0.1724 | 0.0563 | 0.3154 | 54.7914 |
| Endless Forams Original | 27729 | 0.2578 | 0.0839 | 0.1385 | 0.0547 | 0.2578 | 76.3005 |
| Diverse Fossils (RFID) | 59244 | 0.8301 | 0.1408 | 0.5878 | 0.1563 | 0.8301 | 197.3126 |

## 9 Discussion

The choice of using a given ML framework is not a simple analysis of the efficacy of the method in classification. It is a series of trade-offs between efficacy, accessibility, robustness and computational costs. The combined analysis of these factors demonstrates the utility of our pipeline and the potential for it to become the default approach to image classification problems in palaeontology, as was previously the case with CNNs [3]. In section 5, we demonstrated how the use of our methodology gives comparable results with much less data compared to the standard approach of using CNNs. This result is unsurprising, as previous work has found that DINO models [4, 6] generalise well to diverse unseen domains and compare favourably against previous methodologies. Therefore, there is little expectation that palaeontology should behave differently given the known robustness of these methods. Our work also differs from most proposed ML methods in palaeontology in that we propose a general workflow which is data agnostic and benchmark on diverse datasets as opposed to proposing a bespoke methodology and testing it on a single dataset.

Even if our methodology consistently underperformed previous methods, we would argue that it would still be worth adopting, particularly for proof-of-concept studies, as the fact that it can be used without fine-tuning, using a simple classifier, in a few lines of code, using standard well-documented libraries, on smaller amounts of data, with automated preparation is extremely valuable to palaeontology. The flexibility of this approach also means the same methods can be applied for both small and large datasets.

This methodology addresses reproducibility issues in palaeontological ML studies [3] in three ways. Firstly, there are simply fewer parameters required as there is no complex fine-tuning, the data augmentation step can be omitted and the preprocessing is handled automatically. Secondly, as the process of writing the code is faster, and the runtimes shorter, it is more realistic to take the data provided in a study and implement yourself, rather than relying on the code written by others. Thirdly, the large model can be downloaded and accessed directly from Hugging Face [9] negating the need for this to be shared. Only the simple classifier needs to be shared, and even then this could be simply retrained from the data.

Another key benefit of our approach is that, despite our emphasis on DINOv3 [4], it does not actually rely on a particular model and is hence future-proof and flexible. If a new and improved feature extractor is released, it is simply the case of changing the Hugging Face model Id [9] to the new model (or any older model you would like to switch to) and running the pipeline.

A particular benefit to palaeontology is the relative success of few-shot learning and learning on very small datasets. This could unlock new potential applications by allowing ML to be used in small data domains or by giving researchers the confidence to invest resources in a data acquisition programme given promising early results, preventing ideas being abandoned due to result uncertainty.

Fitting a logistic regression or kNN classifier is not a computationally expensive process, and fitting classifier to a dataset of tens of thousands of feature vectors is possible in minutes on a laptop. The runtime of the feature extraction process, which only needs to be run once, will depend on the hardware available. The work on this study performed using an Apple MacBook Pro M4 Max. Using batched image loading and the Apple MPS GPU the smallest data set feature vectors could be extracted in a few minutes and the largest in less than 30 minutes. Unsupervised clustering can become runtime intensive depending on the amount of dimensionality reduction, method and dataset size but this must be evaluated on a case-by-case basis. Removing the requirement for CNN fine-tuning also makes ML in palaeontology more sustainable by reducing the computational costs and need for large amounts of GPU time.

The fact that the off-the-shelf models are so powerful allows insight to be gained into datasets without the need for labels as shown in section 6 on unsupervised learning and in section 8. However, with the caveats that unsupervised learning can be sensitive to methodology setup and structure of the dataset. There may be occasions when classes are distinct, tightly clustered and separable in the embedding space. Or other cases where the classes are close enough such that it may be possible for supervised learning to classify them, but not for distinct groupings to be revealed without labels.

Unsupervised learning on robust feature vectors is likely to group images reflective of actual features of the image data; however, it is difficult to know a priori the cause of the separation if it is taxonomic, and of which rank, if it is morphological or if it is some aspect of image composition. In this work, we mainly focus on a demonstration pipeline, but in practice you would run the unsupervised algorithm using multiple parameter sets and rerun it on large clusters fully explore the structure of the dataset in collaboration with a domain expert.

When performing unsupervised learning in a non-exploratory manner, great care needs to be taken when using non-linear dimension reduction methods such as UMAP [52, 53] as they can distort the underlying distances in the space. Unfortunately, the feature vectors outputted by the models on the datasets do not admit an adequate low-dimensional representation that can be found by linear methods, such as PCA.

Despite the caveats, it is hoped that the unsupervised exploration of these datasets can potentially provide novel biological insight. Another potential unsupervised method, unexplored in this work, is using the embeddings for outlier detection. This could be used for dataset management, finding outliers caused by data contamination or labelling errors, or for the detection of novel objects.

Given the accessibility of this methodology, it may even be useful in situations where ML-based classification is not the overall goal. Just as a PCA of morphological properties can be used as a figure to visualise a dataset in palaeontological papers, one can imagine taking all the images of the collected objects, converting to feature vectors and then plotting using non-linear dimensionality reduction to visualise. Additionally, an argument that objects are morphologically and visually distinct can be supported with relatively little effort by showing that this classification pipeline can achieve near-perfect accuracy on the dataset.

The final speculative proposal of this work is to use the DINOv3 [4] generated feature vectors to quantify the disparity of a dataset. A similar concept was proposed in [57], where it was suggested to measure the diversity of pollen data using entropy-derived measurements based on CNN feature vectors [57]. This should work in the case of a dataset with a number of highly diverse classes in that it will give a quantifiable difference in disparity metrics compared to a dataset composed of a small number of similar classes, if the datasets can be reasonably compared. However, further work and suitable dataset selection is needed to test if this methodology can be useful and correlation with biologically relevant parameters found.

The methods we proposed in section 8, do not claim to be exhaustive or a uniquely optimal choice as given a set of vectors there is a whole host of statistics and measures that could be defined. However, our proposal has merit given the readily accessible nature of the feature vectors and the relative simplicity of the calculations. Given this, it is an almost cost-free addition to a suite of measurements if the utility can be proven. The caveats with this approach are that metrics based on feature vector distances are not intuitively interpretable, and separating effects of shape, texture, pose, colour and other image artifacts may be difficult in practice. However, there may be situations where a fast way to quantify the disparity between large unlabelled assemblages is required, and this provides a solution to this use case.

A wide range of future work could be enabled by this methodology. Coupling this methodology to either DINOv3 or another transformer-based segmentation pipeline could enable the automated processing of large datasets using robotics [47] or thin-section imaging [58]. Feature vectors could be used to track changes in morphology over time, in evolution or ontogeny, enabling differentiation of continuous changes from discrete groupings. Another proposal would be to compare the feature vectors derived distances to phylogenetic distances, comparing phylogenetic trees to trees built from hierarchical clustering in feature vector space. It can be imagined that, this methodology could be useful in activities such as building a fossil identification app requiring fewer training examples and a single model, robust to background.

One key avenue of future work would be to look into how we can use these models to capture the true three-dimensional structure of objects by combining the feature vectors for different views appropriately. Given the simplicity of our classification pipeline, with appropriate normalisation, it may also be possible to concatenate the feature vector with other metadata enabling multi-modal learning. Moreover, image embeddings could be combined with a text embedding model, which embeds descriptions of the object. Feature vectors could also be used to search museum collections for images similar to unknown objects based on similarity scores. Finally, more complex classifiers than those presented here could be used, or the model itself fine-tuned, where the need for accuracy or data merits this.

In conclusion, we have shown how a simple pipeline utilising state-of-the-art pre-trained self-supervised models can match the performance of CNN-based methods and allow unsupervised insight into palaeontological data. In future, it is hoped that this proposed pipeline or variants of it can become widely used, allowing state-of-the-art ML to be accessible to non-specialists, and potentially contributing to new discoveries in palaeontology.

## 10 Data and Code Availability

Code and access links to data for this version of the work are uploaded to GitHub at https://github.com/nrodgers1/Self-Supervised-Learning-for-Palaeontology, on acceptance for publication a final version of this will be placed in a permanent repository with DOI. Potential users of DinoV3 [4] should consult the custom Meta AI license agreement.

## 11 Funding and Acknowledgements

Niall Rodgers acknowledges the support of the Human Frontiers Science Program under grant number RGP006/2024. The author thanks Sean McMahon, Henderson Cleaves and Mark van Zuilen for acquiring this funding and allowing the time to work on this manuscript. The author thanks Sean McMahon for useful discussions and feedback on this work.

For the purpose of open access, the author has applied a Creative Commons Attribution (CC BY) licence to any Author Accepted Manuscript version arising from this submission.

