## Supplementary Information for "Complexity Begets Simplicity: Self-Supervised Learning for Palaeontological Images with Few or No Labels"

In this supplementary information, we provide more detailed classification and clustering metrics, plot confusion matrices for each dataset and show how PCA can be used for feature vector dimension reduction. At the end of the supplementary information we also repeat the plots dimension reduction given in the main text which can be used as reference for the colour label correspondence and examined in more detail.

### S1.1 Supplementary Methods

#### S1.1.1 Classification Metrics

Accuracy alone is not enough to fully understand the behaviour of a classifier; however, it does give a good overall impression of how a classifier performs. As accuracy is the main metric used to illustrate performance in the literature, we focus on this for our literature comparison, but for completeness list other relevant metrics in this supplementary information. The additional metrics used are described below. Firstly, we measure top-2 and top-3 accuracy which measure the fraction of predictions where the true labels are in the top-2 or top-3 predictions, respectively. This is useful when working with a large number of classes as it gives a sense if the errors are between similar classes and if the classifier has learned the structure of the data or if the predictions are more random. We also measure the balanced accuracy, which re-weights the accuracy to account for class imbalance. We also measure the standard metrics of precision, recall, specificity and F1 scores, for each of these metrics we provide the 'macro' version from scikit-learn [50] which does not account for class imbalance aside from the F1 score where we also measure the 'weighted' version which does. The precision is the ratio of the True Positives to the sum of True Positives and False Positives, averaged across classes which gives the fraction of positive classifications which are correct. The recall score is the ratio of the True Positives to the sum of the True Positives and False Negatives, which is the fraction of positives identified. The specificity is the ratio of the True Negatives over the sum of the True Negatives and False Positives, which gives the fraction of negative classifications identified. While, the F1 score is the harmonic mean of the precision and recall.

#### S1.1.2 Confusion Matrices

Confusion matrices are a way to visualise the performance across classes, allowing the per-class accuracy to be visualised and combinations of classes that are most likely to be misclassified to be understood. All the confusion matrices plotted are the result of cross-validated fitting of class-balanced logistic regression, passing the full datasets into this cross-validation pipeline. This is done using five folds generated using the stratified fold function in scikit-learn [50] which are passed the cross-validation predict function which predicts and fits across folds, allowing a normalised confusion matrix to be constructed from the predictions on the held-back test sets.

#### S1.1.3 Clustering Metrics

For the sake of brevity, we do not repeat the explanation the clustering metrics stated in the main text. However, highlight that all the metrics are scikit-learn [50] functions, so definitions can be read directly from the documentation. We do, however, remind the reader of the acronyms Adjusted Rand Index (ARI), Adjusted Mutual Information (AMI) and Davies–Bouldin index (DBI).

### S1.2 Feature Visualisation with PCA

In figure S1, we show the application of PCA to reduce the dimensions of the feature vectors. In general, points of the same class are placed together; however, it is difficult to separate the classes as the classes cluster too tightly together. In the case of radiolaria, figure S1c, as classes are relatively easy to classify and there are a relatively small number of classes, the PCA is able to separate out some of the classes. However, for larger datasets with more classes the PCA struggles to be interpretable. In general, we recommend that non-linear dimensionality reduction techniques are likely more effectively used on the DINOv3 vectors; however, there may be cases where PCA gives adequate insight into the data.

### S1.3 Pollen Dataset

In this section, we refer to Pollen Dataset as the data of New Zealand Pollen compromising approximately 19,000 images and 46 classes taken from [14], described in the main text.

#### S1.3.1 Classification Results

Table S1: Pollen Classification results with mean ± standard deviation over folds and repeats

| Train Fraction | Classifier | Top-1 Accuracy | Top-2 Accuracy | Top-3 Accuracy | F1 Macro | F1 Weighted | Precision Macro | Recall Macro | Specificity Macro | Balanced Accuracy |
|---|---|---|---|---|---|---|---|---|---|---|
| 0.10 | Logistic Regression | 0.951 ± 0.013 | 0.989 ± 0.005 | 0.995 ± 0.003 | 0.931 ± 0.022 | 0.950 ± 0.013 | 0.943 ± 0.020 | 0.931 ± 0.022 | 0.999 ± 0.000 | 0.931 ± 0.022 |
| 0.10 | k-Nearest Neighbours | 0.887 ± 0.016 | 0.961 ± 0.009 | 0.973 ± 0.008 | 0.825 ± 0.027 | 0.879 ± 0.018 | 0.864 ± 0.027 | 0.816 ± 0.027 | 0.997 ± 0.000 | 0.816 ± 0.027 |
| 0.30 | Logistic Regression | 0.972 ± 0.005 | 0.995 ± 0.002 | 0.998 ± 0.001 | 0.967 ± 0.008 | 0.972 ± 0.005 | 0.971 ± 0.007 | 0.966 ± 0.008 | 0.999 ± 0.000 | 0.966 ± 0.008 |
| 0.30 | k-Nearest Neighbours | 0.925 ± 0.007 | 0.977 ± 0.004 | 0.983 ± 0.003 | 0.900 ± 0.013 | 0.924 ± 0.007 | 0.933 ± 0.011 | 0.885 ± 0.015 | 0.998 ± 0.000 | 0.885 ± 0.015 |
| 0.50 | Logistic Regression | 0.978 ± 0.003 | 0.997 ± 0.001 | **0.999 ± 0.001** | 0.973 ± 0.005 | 0.978 ± 0.003 | 0.975 ± 0.005 | 0.972 ± 0.005 | 0.999 ± 0.000 | 0.972 ± 0.005 |
| 0.50 | k-Nearest Neighbours | 0.938 ± 0.005 | 0.981 ± 0.003 | 0.986 ± 0.003 | 0.921 ± 0.010 | 0.938 ± 0.005 | 0.945 ± 0.009 | 0.907 ± 0.011 | 0.999 ± 0.000 | 0.907 ± 0.011 |
| 0.70 | Logistic Regression | 0.981 ± 0.003 | 0.997 ± 0.001 | 0.999 ± 0.001 | 0.977 ± 0.005 | 0.981 ± 0.003 | 0.978 ± 0.005 | 0.977 ± 0.005 | **1.000 ± 0.000** | 0.977 ± 0.005 |
| 0.70 | k-Nearest Neighbours | 0.945 ± 0.004 | 0.984 ± 0.002 | 0.988 ± 0.002 | 0.930 ± 0.008 | 0.944 ± 0.004 | 0.950 ± 0.007 | 0.918 ± 0.009 | 0.999 ± 0.000 | 0.918 ± 0.009 |
| 1.00 | Logistic Regression | **0.983 ± 0.002** | **0.998 ± 0.001** | 0.999 ± 0.001 | **0.980 ± 0.003** | **0.983 ± 0.002** | **0.981 ± 0.003** | **0.980 ± 0.003** | 1.000 ± 0.000 | **0.980 ± 0.003** |
| 1.00 | k-Nearest Neighbours | 0.951 ± 0.003 | 0.986 ± 0.002 | 0.989 ± 0.002 | 0.938 ± 0.005 | 0.950 ± 0.003 | 0.954 ± 0.004 | 0.928 ± 0.005 | 0.999 ± 0.000 | 0.928 ± 0.005 |

The Pollen Dataset, table S1, shows extremely strong results of the classification metrics. With only 10% of the data, all metrics are above 90% and the top-3 accuracy is close to 100% for logistic regression. When the whole dataset is passed to the pipeline, all metrics approach 100% for logistic regression, with kNN also performing very well. This demonstrates that for well separated and distinguishable classes our proposed pipeline can reach the upper limits of possible performance, negating the need for a more complex pipeline.

The confusion matrix for the pollen, figure S2, also shows excellent performance. Most classes have performance in the high 90% range. The lowest performing class *Salix* is confused mostly for *Weinmannia* which looks very similar in certain images, showing errors tend not to be random in nature.

Again, the few-shot results are very strong on the pollen dataset, above 50% in all metrics can be gained with a single example per class. There is a rapid increase in performance with more data, table S2. This highlights the utility of this method for small datasets and proof-of-concept studies.

#### S1.3.2 Clustering Metrics

When clustering the pollen dataset, table S3, UMAP is generally much better than using PCA as a preprocessing step. Meanwhile, the performance of the clustering method depends on the scale at which it chooses to cluster. If the method chooses a number of clusters which is broadly similar to the true number of classes the metrics which depend on the true labels are generally better, while if the algorithm clusters at a higher level of organisation then the label-independent measure of performance are best. This demonstrates how unsupervised learning can quite successfully identify structure in the pollen dataset; however, there can be variation between algorithms on the scale of clusters detected.

### S1.4 Fossil Tracks Dataset

In this section, we discuss the data of [13], described in the main text, which contains approximately 1700 binary images of fossil tracks broken into ornithischians and theropod classes. This dataset additionally contains many images which are fundamentally ambiguous, and it is unclear to which class they should belong.

#### S1.4.1 Classification Results

When classifying the fossil tracks, table S4, we do not give a top-2 or top-3 accuracy score, as this does not make sense for a binary classification problem. The performance of the classifier is relatively consistent as more data is added as a result of the ambiguity between classes. All metrics plateau near 80%, which is still a reasonably good performance due to the ambiguity. These results highlight that this pipeline cannot

Table S2: Few Shot Classification Results - Pollen Dataset

| Few-Shot k | Classifier | Repeats Successful | Top-1 Accuracy | Top-2 Accuracy | Top-3 Accuracy | F1 Macro | F1 Weighted | Precision Macro | Recall Macro | Specificity Macro | Balanced Accuracy |
|---|---|---|---|---|---|---|---|---|---|---|---|
| 1 | k-Nearest Neighbours | 50 | 0.540 ± 0.030 | 0.546 ± 0.029 | 0.558 ± 0.030 | 0.511 ± 0.026 | 0.536 ± 0.033 | 0.572 ± 0.023 | 0.550 ± 0.024 | 0.990 ± 0.001 | 0.550 ± 0.024 |
| 1 | Logistic Regression | 50 | 0.570 ± 0.035 | 0.720 ± 0.032 | 0.795 ± 0.029 | 0.538 ± 0.029 | 0.564 ± 0.036 | 0.573 ± 0.024 | 0.580 ± 0.030 | 0.990 ± 0.001 | 0.580 ± 0.030 |
| 2 | k-Nearest Neighbours | 50 | 0.533 ± 0.023 | 0.767 ± 0.025 | 0.771 ± 0.024 | 0.470 ± 0.024 | 0.500 ± 0.034 | 0.581 ± 0.020 | 0.514 ± 0.023 | 0.989 ± 0.001 | 0.514 ± 0.023 |
| 2 | Logistic Regression | 50 | 0.702 ± 0.021 | 0.838 ± 0.022 | 0.894 ± 0.018 | 0.682 ± 0.018 | 0.702 ± 0.022 | 0.689 ± 0.018 | 0.719 ± 0.014 | 0.993 ± 0.001 | 0.719 ± 0.014 |
| 3 | k-Nearest Neighbours | 50 | 0.605 ± 0.024 | 0.797 ± 0.021 | 0.863 ± 0.018 | 0.572 ± 0.022 | 0.596 ± 0.029 | 0.643 ± 0.020 | 0.611 ± 0.020 | 0.991 ± 0.001 | 0.611 ± 0.020 |
| 3 | Logistic Regression | 50 | 0.760 ± 0.018 | 0.881 ± 0.018 | 0.925 ± 0.016 | 0.746 ± 0.015 | 0.759 ± 0.018 | 0.743 ± 0.015 | 0.779 ± 0.012 | 0.995 ± 0.000 | 0.779 ± 0.012 |
| 5 | k-Nearest Neighbours | 50 | 0.695 ± 0.019 | 0.846 ± 0.017 | 0.907 ± 0.013 | 0.660 ± 0.014 | 0.678 ± 0.020 | 0.694 ± 0.012 | 0.693 ± 0.013 | 0.993 ± 0.000 | 0.693 ± 0.013 |
| 5 | Logistic Regression | 50 | 0.818 ± 0.014 | 0.923 ± 0.013 | 0.956 ± 0.010 | 0.809 ± 0.012 | 0.818 ± 0.015 | 0.800 ± 0.013 | 0.839 ± 0.009 | 0.996 ± 0.000 | 0.839 ± 0.009 |
| 10 | k-Nearest Neighbours | 50 | 0.784 ± 0.011 | 0.909 ± 0.009 | 0.945 ± 0.008 | 0.762 ± 0.011 | 0.775 ± 0.014 | 0.769 ± 0.010 | 0.791 ± 0.009 | 0.995 ± 0.000 | 0.791 ± 0.009 |
| 10 | Logistic Regression | 50 | 0.884 ± 0.010 | 0.964 ± 0.006 | 0.983 ± 0.004 | 0.880 ± 0.008 | 0.884 ± 0.010 | 0.869 ± 0.010 | 0.899 ± 0.006 | 0.997 ± 0.000 | 0.899 ± 0.006 |
| 20 | k-Nearest Neighbours | 50 | 0.849 ± 0.007 | 0.946 ± 0.004 | 0.967 ± 0.004 | 0.831 ± 0.007 | 0.844 ± 0.008 | 0.826 ± 0.007 | 0.857 ± 0.006 | 0.996 ± 0.000 | 0.857 ± 0.006 |
| 20 | Logistic Regression | 50 | **0.926 ± 0.005** | **0.983 ± 0.003** | **0.992 ± 0.002** | **0.923 ± 0.005** | **0.926 ± 0.005** | **0.914 ± 0.006** | **0.935 ± 0.004** | **0.998 ± 0.000** | **0.935 ± 0.004** |

necessarily remove fundamental ambiguities in the data and that DINOv3 models trained on a large complex dataset can generalise quite well to binary shapes.

The confusion matrix, figure S3, is unsurprising given what has been observed about this binary classification problem. We classify most of each class correctly, but with a still significant amount of errors falsely placing the object in the opposite class.

Few-shot classification of the tracks paints a similar picture to previous results, table S4. Where again we omit the top-2 and top-3 accuracy. Due to the difficulty of the classification problem, it takes more data to improve beyond the baseline of predicting the most frequent class. The results still demonstrate that few-shot learning is possible and much better than would be possible in a CNN-based approach but that the success of the approach depends on characteristics of the data in question.

#### S1.4.2 Clustering Metrics

Due to the difficulty of the fossil tracks separability, clustering results are quite poor, table S6. Some algorithms find clusters which are reasonable without respect to the labelling, but due to the fact the data is continuous and does not fall into discrete groups, finding high-quality separable clusters is not possible. It is expected that the found clusters do represent genuine morphological groupings in the data, just ones that are continuous at the boundaries and do not align with the given labels. This data highlights the difficulties of unsupervised clustering, but also how it can inform you about the structure of the data.

### S1.5 Radiolaria

The radiolaria data we refer to in the SI is the dataset of [12] referred to in the main text. This data contains 8 classes and 1085 images.

#### S1.5.1 Classification Results

The radiolaria data is generally very well classified even when using a small fraction of the data, table S7. Using the maximum amount of data both the logistic regression and the kNN classifiers score above 90% in

Table S3: Clustering results - Pollen Dataset

| Pipeline | Algorithm | No. of Clusters | ARI | AMI | Purity | Silhouette | DBI | Noise Fraction | Components Used |
|---|---|---|---|---|---|---|---|---|---|
| PCA | KMeans (Known K) | 46 | 0.4805 | 0.7568 | 0.7633 | 0.1407 | 1.9950 | 0.000000 | 50 |
| PCA | KMeans (Unknown K) | 3 | 0.0909 | 0.3589 | 0.1942 | 0.1689 | 1.8855 | 0.000000 | 50 |
| PCA | Bayesian GMM | 50 | 0.5625 | 0.8234 | 0.8363 | 0.1330 | 2.0443 | 0.000000 | 50 |
| PCA | HDBSCAN | 3 | 0.0122 | 0.2044 | 0.0933 | 0.4010 | 1.0315 | 0.787105 | 50 |
| Exploratory UMAP | KMeans (Known K) | 46 | 0.6298 | 0.8711 | **0.8753** | 0.6332 | 0.5032 | 0.000000 | 15 |
| Exploratory UMAP | KMeans (Unknown K) | 12 | 0.3312 | 0.7203 | 0.4450 | **0.7527** | **0.2995** | 0.000000 | 15 |
| Exploratory UMAP | Bayesian GMM | 29 | **0.6833** | **0.8915** | 0.7258 | 0.6702 | 0.3459 | 0.000000 | 15 |
| Exploratory UMAP | HDBSCAN | 12 | 0.3801 | 0.7470 | 0.4757 | 0.6863 | 0.3199 | 0.089948 | 15 |

Table S4: Classification results with mean ± standard deviation over folds and repeats - Fossil Tracks Dataset

| Train Fraction | Classifier | Top-1 Accuracy | Top-2 Accuracy | Top-3 Accuracy | F1 Macro | F1 Weighted | Precision Macro | Recall Macro | Specificity Macro | Balanced Accuracy |
|---|---|---|---|---|---|---|---|---|---|---|
| 0.10 | Logistic Regression | 0.775 ± 0.057 | N/A | N/A | 0.765 ± 0.063 | 0.773 ± 0.060 | 0.773 ± 0.058 | 0.767 ± 0.064 | 0.767 ± 0.064 | 0.767 ± 0.064 |
| 0.10 | k-Nearest Neighbours | 0.755 ± 0.074 | N/A | N/A | 0.741 ± 0.079 | 0.751 ± 0.076 | 0.758 ± 0.083 | 0.740 ± 0.077 | 0.740 ± 0.077 | 0.740 ± 0.077 |
| 0.30 | Logistic Regression | 0.801 ± 0.046 | N/A | N/A | 0.794 ± 0.047 | 0.800 ± 0.046 | 0.797 ± 0.048 | 0.795 ± 0.046 | 0.795 ± 0.046 | 0.795 ± 0.046 |
| 0.30 | k-Nearest Neighbours | 0.772 ± 0.044 | N/A | N/A | 0.760 ± 0.048 | 0.769 ± 0.046 | 0.771 ± 0.048 | 0.757 ± 0.047 | 0.757 ± 0.047 | 0.757 ± 0.047 |
| 0.50 | Logistic Regression | 0.812 ± 0.028 | N/A | N/A | 0.806 ± 0.029 | 0.812 ± 0.028 | **0.808 ± 0.029** | 0.807 ± 0.028 | 0.807 ± 0.028 | 0.807 ± 0.028 |
| 0.50 | k-Nearest Neighbours | 0.784 ± 0.028 | N/A | N/A | 0.773 ± 0.030 | 0.781 ± 0.029 | 0.782 ± 0.030 | 0.769 ± 0.030 | 0.769 ± 0.030 | 0.769 ± 0.030 |
| 0.70 | Logistic Regression | 0.812 ± 0.024 | N/A | N/A | **0.807 ± 0.025** | 0.812 ± 0.024 | 0.807 ± 0.025 | **0.808 ± 0.026** | **0.808 ± 0.026** | **0.808 ± 0.026** |
| 0.70 | k-Nearest Neighbours | 0.790 ± 0.027 | N/A | N/A | 0.780 ± 0.028 | 0.788 ± 0.027 | 0.788 ± 0.029 | 0.776 ± 0.028 | 0.776 ± 0.028 | 0.776 ± 0.028 |
| 1.00 | Logistic Regression | **0.813 ± 0.022** | N/A | N/A | 0.807 ± 0.023 | **0.813 ± 0.022** | 0.808 ± 0.023 | 0.808 ± 0.022 | 0.808 ± 0.022 | 0.808 ± 0.022 |
| 1.00 | k-Nearest Neighbours | 0.799 ± 0.024 | N/A | N/A | 0.789 ± 0.025 | 0.797 ± 0.024 | 0.796 ± 0.025 | 0.786 ± 0.026 | 0.786 ± 0.026 | 0.786 ± 0.026 |

all metrics and score begin to plateau near the maximum value after using only half the data. This shows our pipeline performs well on distinct objects in small data scenarios.

The confusion matrix for the radiolaria data, figure S4, shows strong performance across all classes aside from *Diamesa* which is confused with *Papalis*. This is unsurprising as, particularly in certain images, both these classes look very similar, demonstrating that the classifier has separated them into a distinct region of feature space.

The few-shot classification performance on the radiolaria dataset, table S8, is generally very good with reasonable results across all metrics with only a few examples and a smooth increase in performance with increased data. In particular, the top-2 accuracy is above 90% for only 5 examples in logistic regression showing how the DINOv3 model separates the groups of classes in feature space.

#### S1.5.2 Clustering Metrics

Again, when clustering the radiolarian dataset, table S7, the UMAP dimension reduction performs much more consistently than the PCA. Clustering results on the radiolaria data are generally solid, however, with the caveat that different algorithms may group more some similar species but not others creating a wide variety of possible groupings.

### S1.6 Foraminifera

This section of SI refers to the dataset of Foraminifera images taken from [11, 49] which was derived from [10] which contains approximately 27,000 images spread over 35 classes.

Table S5: Few Shot Classification Results - Fossils Tracks Dataset

| Few-Shot k | Classifier | Repeats Successful | Top-1 Accuracy | Top-2 Accuracy | Top-3 Accuracy | F1 Macro | F1 Weighted | Precision Macro | Recall Macro | Specificity Macro | Balanced Accuracy |
|---|---|---|---|---|---|---|---|---|---|---|---|
| 1 | k-Nearest Neighbours | 50 | 0.561 ± 0.104 | N/A | N/A | 0.515 ± 0.102 | 0.531 ± 0.109 | 0.564 ± 0.107 | 0.541 ± 0.087 | 0.541 ± 0.087 | 0.541 ± 0.087 |
| 1 | Logistic Regression | 50 | 0.590 ± 0.093 | N/A | N/A | 0.550 ± 0.101 | 0.565 ± 0.104 | 0.600 ± 0.100 | 0.573 ± 0.082 | 0.573 ± 0.082 | 0.573 ± 0.082 |
| 2 | k-Nearest Neighbours | 50 | 0.609 ± 0.052 | N/A | N/A | 0.468 ± 0.115 | 0.449 ± 0.132 | 0.567 ± 0.093 | 0.554 ± 0.066 | 0.554 ± 0.066 | 0.554 ± 0.066 |
| 2 | Logistic Regression | 50 | 0.605 ± 0.093 | N/A | N/A | 0.584 ± 0.098 | 0.591 ± 0.102 | 0.613 ± 0.091 | 0.601 ± 0.085 | 0.601 ± 0.085 | 0.601 ± 0.085 |
| 3 | k-Nearest Neighbours | 50 | 0.601 ± 0.077 | N/A | N/A | 0.568 ± 0.081 | 0.580 ± 0.084 | 0.614 ± 0.078 | 0.588 ± 0.066 | 0.588 ± 0.066 | 0.588 ± 0.066 |
| 3 | Logistic Regression | 50 | 0.625 ± 0.068 | N/A | N/A | 0.608 ± 0.068 | 0.619 ± 0.068 | 0.623 ± 0.070 | 0.614 ± 0.064 | 0.614 ± 0.064 | 0.614 ± 0.064 |
| 5 | k-Nearest Neighbours | 50 | 0.625 ± 0.066 | N/A | N/A | 0.593 ± 0.077 | 0.608 ± 0.075 | 0.628 ± 0.070 | 0.606 ± 0.061 | 0.606 ± 0.061 | 0.606 ± 0.061 |
| 5 | Logistic Regression | 50 | 0.642 ± 0.061 | N/A | N/A | 0.629 ± 0.060 | 0.638 ± 0.063 | 0.645 ± 0.057 | 0.637 ± 0.052 | 0.637 ± 0.052 | 0.637 ± 0.052 |
| 10 | k-Nearest Neighbours | 50 | 0.641 ± 0.048 | N/A | N/A | 0.625 ± 0.048 | 0.637 ± 0.048 | 0.635 ± 0.048 | 0.629 ± 0.045 | 0.629 ± 0.045 | 0.629 ± 0.045 |
| 10 | Logistic Regression | 50 | 0.674 ± 0.058 | N/A | N/A | 0.668 ± 0.056 | 0.674 ± 0.057 | 0.674 ± 0.054 | 0.674 ± 0.053 | 0.674 ± 0.053 | 0.674 ± 0.053 |
| 20 | k-Nearest Neighbours | 50 | 0.667 ± 0.050 | N/A | N/A | 0.658 ± 0.046 | 0.666 ± 0.049 | 0.666 ± 0.044 | 0.663 ± 0.041 | 0.663 ± 0.041 | 0.663 ± 0.041 |
| 20 | Logistic Regression | 50 | **0.718 ± 0.031** | N/A | N/A | **0.713 ± 0.030** | **0.719 ± 0.031** | **0.716 ± 0.029** | **0.719 ± 0.028** | **0.719 ± 0.028** | **0.719 ± 0.028** |

##### S1.6.1 Classification Results

As the foraminifera dataset is a relatively large, performance, table S10, is good with only a small fraction of the data used, and the high top-3 accuracy shows it can quickly learn to group similar foraminifera. This shows that supervised classification can still perform well despite differing views of the foraminifera and differences in image lighting conditions.

The confusion matrix, figure S5, shows generally good performance across most classes but a few outliers with very poor performance. The very poorly performing classes generally have a very small number of images, which explains why the pipeline struggles to separate them as effectively as the larger classes.

Few-shot performance, table S11, is weaker on the foraminifera dataset since it takes more examples for the pipeline to build a representative set of images for each class. However, top-3 accuracy does become reasonable, showing how some understanding of the data is found. Few-shot performance is still indicative that these objects could be classified given more data and could be useful in a proof-of-concept approach.

##### S1.6.2 Clustering Metrics

The outcome of the clustering of the foraminifera data, table S12, depends heavily on the chosen algorithm and dimension reduction method. UMAP is generally better than PCA, with PCA based HDBSCAN failing completely. Some clustering methods, such as unknown k k-means and HDBSCAN, focus on the two large clusters caused by imaging artifacts. While, the Bayesian Gaussian mixture model finds a set of clusters which partially align with the underlying labels. This shows how care needs to be taken not to identify image artifacts with this methodology; however, this could be remedied by clustering on the large macroscopic clusters again individually and inspecting the clusters to find alignment.

#### S1.7 Diverse Palaeontological Images Dataset (RFID)

This section in the SI concerns the diverse RFID [15], described in the main text, which contains approximately 60,000 images taken from the Internet comprising 50 classes.

Table S6: Clustering results - Fossil Tracks Dataset

| Pipeline | Algorithm | No. of Clusters | ARI | AMI | Purity | Silhouette | DBI | Noise Fraction | Components Used |
|---|---|---|---|---|---|---|---|---|---|
| PCA | KMeans (Known K) | 2 | 0.0385 | 0.0256 | 0.5990 | 0.1640 | 2.1381 | 0.000000 | 50 |
| PCA | KMeans (Unknown K) | 2 | 0.0385 | 0.0256 | 0.5990 | 0.1640 | 2.1381 | 0.000000 | 50 |
| PCA | Bayesian GMM | 4 | 0.0493 | 0.0378 | 0.6290 | 0.1187 | 2.4838 | 0.000000 | 50 |
| PCA | HDBSCAN | 2 | 0.0003 | 0.0489 | 0.1305 | 0.4298 | 0.7397 | 0.833633 | 50 |
| Exploratory UMAP | KMeans (Known K) | 2 | 0.0102 | 0.0019 | 0.5859 | 0.5333 | 0.6047 | 0.000000 | 15 |
| Exploratory UMAP | KMeans (Unknown K) | 3 | **0.0973** | 0.0612 | **0.6517** | **0.5396** | 0.6413 | 0.000000 | 15 |
| Exploratory UMAP | Bayesian GMM | 4 | 0.0817 | 0.0500 | 0.6349 | 0.4360 | 0.7782 | 0.000000 | 15 |
| Exploratory UMAP | HDBSCAN | 3 | 0.0251 | **0.0660** | 0.6236 | 0.4906 | **0.4666** | 0.008977 | 15 |

Table S7: Classification results with mean ± standard deviation over folds and repeats - Radiolaria Dataset

| Train Fraction | Classifier | Top-1 Accuracy | Top-2 Accuracy | Top-3 Accuracy | F1 Macro | F1 Weighted | Precision Macro | Recall Macro | Specificity Macro | Balanced Accuracy |
|---|---|---|---|---|---|---|---|---|---|---|
| 0.10 | Logistic Regression | 0.859 ± 0.068 | 0.969 ± 0.037 | 0.989 ± 0.022 | 0.803 ± 0.094 | 0.840 ± 0.072 | 0.815 ± 0.099 | 0.824 ± 0.084 | 0.979 ± 0.010 | 0.824 ± 0.084 |
| 0.10 | k-Nearest Neighbours | 0.775 ± 0.069 | 0.924 ± 0.053 | 0.960 ± 0.045 | 0.645 ± 0.120 | 0.734 ± 0.091 | 0.647 ± 0.139 | 0.670 ± 0.108 | 0.965 ± 0.012 | 0.670 ± 0.108 |
| 0.30 | Logistic Regression | 0.917 ± 0.029 | 0.990 ± 0.010 | 0.998 ± 0.005 | 0.904 ± 0.037 | 0.915 ± 0.030 | 0.922 ± 0.033 | 0.900 ± 0.038 | 0.987 ± 0.004 | 0.900 ± 0.038 |
| 0.30 | k-Nearest Neighbours | 0.871 ± 0.035 | 0.975 ± 0.020 | 0.986 ± 0.013 | 0.816 ± 0.058 | 0.852 ± 0.041 | 0.875 ± 0.061 | 0.804 ± 0.057 | 0.979 ± 0.006 | 0.804 ± 0.057 |
| 0.50 | Logistic Regression | 0.932 ± 0.019 | 0.992 ± 0.009 | **0.999 ± 0.003** | 0.926 ± 0.023 | 0.931 ± 0.020 | 0.932 ± 0.021 | 0.924 ± 0.026 | 0.990 ± 0.003 | 0.924 ± 0.026 |
| 0.50 | k-Nearest Neighbours | 0.895 ± 0.030 | 0.981 ± 0.015 | 0.989 ± 0.010 | 0.866 ± 0.041 | 0.887 ± 0.032 | 0.907 ± 0.039 | 0.852 ± 0.042 | 0.983 ± 0.005 | 0.852 ± 0.042 |
| 0.70 | Logistic Regression | 0.938 ± 0.019 | 0.993 ± 0.006 | 0.999 ± 0.003 | 0.935 ± 0.021 | 0.938 ± 0.019 | 0.940 ± 0.019 | 0.934 ± 0.024 | **0.991 ± 0.003** | 0.934 ± 0.024 |
| 0.70 | k-Nearest Neighbours | 0.910 ± 0.020 | 0.984 ± 0.008 | 0.991 ± 0.006 | 0.888 ± 0.029 | 0.902 ± 0.023 | 0.919 ± 0.020 | 0.875 ± 0.031 | 0.985 ± 0.003 | 0.875 ± 0.031 |
| 1.00 | Logistic Regression | **0.943 ± 0.015** | **0.994 ± 0.005** | 0.999 ± 0.002 | **0.940 ± 0.016** | **0.943 ± 0.015** | **0.942 ± 0.017** | **0.940 ± 0.017** | 0.991 ± 0.002 | **0.940 ± 0.017** |
| 1.00 | k-Nearest Neighbours | 0.921 ± 0.018 | 0.985 ± 0.008 | 0.991 ± 0.005 | 0.901 ± 0.023 | 0.913 ± 0.018 | 0.924 ± 0.018 | 0.888 ± 0.025 | 0.987 ± 0.003 | 0.888 ± 0.025 |

#### S1.7.1 Classification Results - Diverse Palaeontological Images

Classification performance is generally very good on the RFID, table S13, given the variations in image quality in this dataset. However, as DINOv3 is trained on such a large volume of images, this is the scenario in which it is expected to perform well. Interestingly, on this data the kNN classifier and the logistic regression perform very similarly, with kNN even exceeding the performance of the logistic regression on some metrics. This is likely due to how the DINOv3 feature vectors separate the diverse objects in feature space.

The confusion matrix, figure S6, shows generally very good performance across most classes, with performance being weakest on classes which lack strong distinct structures such as bone fragments, bryozoans, sponges and trace fossils. The classifier also struggles to distinguish bivavles and brachiopods which demonstrates that the classifier behaves in a somewhat predictable manner as these are both classes which are difficult for people to distinguish.

Given the diversity of image types within each class, it takes several examples for few-shot performance, table 3e, to begin to increase to reasonable levels. However, with enough examples we see adequate few shot performance even on a dataset containing as much diversity this.

#### S1.7.2 Clustering Metrics

Clustering of the diverse images, table S15, again shows UMAP performs better than PCA, with PCA HDBSCAN failing. The algorithms are able to find moderately good clusters. However, these do not necessarily reflect the labelling, as in a dataset as diverse, in species and representation, it means that there will be many ways to group the objects.

Table S8: Few-Shot Classification Results - Radiolaria Dataset

| Few-Shot k | Classifier | Repeats Successful | Top-1 Accuracy | Top-2 Accuracy | Top-3 Accuracy | F1 Macro | F1 Weighted | Precision Macro | Recall Macro | Specificity Macro | Balanced Accuracy |
|---|---|---|---|---|---|---|---|---|---|---|---|
| 1 | k-Nearest Neighbours | 50 | 0.562 ± 0.065 | 0.598 ± 0.071 | 0.635 ± 0.068 | 0.536 ± 0.053 | 0.564 ± 0.069 | 0.590 ± 0.048 | 0.568 ± 0.050 | 0.938 ± 0.009 | 0.568 ± 0.050 |
| 1 | Logistic Regression | 50 | 0.579 ± 0.063 | 0.780 ± 0.057 | 0.885 ± 0.040 | 0.550 ± 0.051 | 0.580 ± 0.064 | 0.586 ± 0.047 | 0.583 ± 0.045 | 0.940 ± 0.008 | 0.583 ± 0.045 |
| 2 | k-Nearest Neighbours | 50 | 0.522 ± 0.060 | 0.802 ± 0.053 | 0.820 ± 0.052 | 0.498 ± 0.066 | 0.524 ± 0.074 | 0.589 ± 0.057 | 0.553 ± 0.048 | 0.934 ± 0.009 | 0.553 ± 0.048 |
| 2 | Logistic Regression | 50 | 0.654 ± 0.043 | 0.843 ± 0.038 | 0.926 ± 0.029 | 0.638 ± 0.040 | 0.659 ± 0.042 | 0.656 ± 0.040 | 0.667 ± 0.039 | 0.951 ± 0.006 | 0.667 ± 0.039 |
| 3 | k-Nearest Neighbours | 50 | 0.589 ± 0.046 | 0.788 ± 0.048 | 0.913 ± 0.034 | 0.582 ± 0.037 | 0.603 ± 0.049 | 0.644 ± 0.043 | 0.631 ± 0.029 | 0.944 ± 0.005 | 0.631 ± 0.029 |
| 3 | Logistic Regression | 50 | 0.709 ± 0.044 | 0.883 ± 0.036 | 0.951 ± 0.023 | 0.701 ± 0.041 | 0.715 ± 0.044 | 0.711 ± 0.039 | 0.730 ± 0.031 | 0.959 ± 0.006 | 0.730 ± 0.031 |
| 5 | k-Nearest Neighbours | 50 | 0.678 ± 0.036 | 0.876 ± 0.030 | 0.954 ± 0.016 | 0.658 ± 0.036 | 0.680 ± 0.039 | 0.688 ± 0.042 | 0.686 ± 0.030 | 0.953 ± 0.005 | 0.686 ± 0.030 |
| 5 | Logistic Regression | 50 | 0.768 ± 0.032 | 0.925 ± 0.024 | 0.975 ± 0.013 | 0.763 ± 0.030 | 0.775 ± 0.032 | 0.764 ± 0.029 | 0.788 ± 0.028 | 0.967 ± 0.004 | 0.788 ± 0.028 |
| 10 | k-Nearest Neighbours | 50 | 0.764 ± 0.029 | 0.926 ± 0.020 | 0.973 ± 0.010 | 0.755 ± 0.023 | 0.765 ± 0.029 | 0.777 ± 0.026 | 0.776 ± 0.021 | 0.965 ± 0.004 | 0.776 ± 0.021 |
| 10 | Logistic Regression | 50 | 0.837 ± 0.029 | 0.965 ± 0.014 | 0.991 ± 0.005 | 0.838 ± 0.026 | 0.843 ± 0.027 | 0.834 ± 0.025 | 0.858 ± 0.022 | 0.977 ± 0.004 | 0.858 ± 0.022 |
| 20 | k-Nearest Neighbours | 50 | 0.812 ± 0.021 | 0.951 ± 0.013 | 0.980 ± 0.009 | 0.816 ± 0.018 | 0.819 ± 0.023 | 0.824 ± 0.017 | 0.841 ± 0.020 | 0.973 ± 0.004 | 0.841 ± 0.020 |
| 20 | Logistic Regression | 50 | **0.882 ± 0.017** | **0.982 ± 0.005** | **0.996 ± 0.002** | **0.883 ± 0.015** | **0.887 ± 0.015** | **0.877 ± 0.014** | **0.900 ± 0.016** | **0.983 ± 0.002** | **0.900 ± 0.016** |

### S1.8 Larger Images from the Main Text

In this section, we repeat the dimension reduction figures given in the main text but now given each figure its own page so that details can be observed more clearly and so that the correspondence between colours and labels can be clearly read off from the figure legends.

#### S1.8.1 Dimension Reduction with True Labels from Main Text

Non-linear dimensionality reduction methods applied to feature vectors for all images across all datasets with points labelled according to their true class.

#### S1.8.2 Clustering Plots from Main Text

In this section, we replot the clustering results described in the main text, in a larger format.

Table S9: Clustering Results - Radiolaria Dataset

| Pipeline | Algorithm | No. of Clusters | ARI | AMI | Purity | Silhouette | DBI | Noise Fraction | Components Used |
|---|---|---|---|---|---|---|---|---|---|
| PCA | KMeans (Known K) | 8 | 0.4768 | 0.6210 | 0.7327 | 0.1387 | 2.0850 | 0.000000 | 50 |
| PCA | KMeans (Unknown K) | 3 | 0.4019 | 0.5681 | 0.5456 | 0.1979 | 1.7619 | 0.000000 | 50 |
| PCA | Bayesian GMM | 16 | 0.3288 | 0.5685 | 0.7558 | 0.0842 | 2.6096 | 0.000000 | 50 |
| PCA | HDBSCAN | 2 | 0.0719 | 0.1634 | 0.2378 | 0.2303 | 1.2640 | 0.456221 | 50 |
| Exploratory UMAP | KMeans (Known K) | 8 | 0.5193 | 0.6740 | 0.7475 | 0.6255 | 0.5220 | 0.000000 | 15 |
| Exploratory UMAP | KMeans (Unknown K) | 4 | 0.5643 | 0.7252 | 0.7051 | **0.7234** | **0.3704** | 0.000000 | 15 |
| Exploratory UMAP | Bayesian GMM | 15 | 0.4907 | 0.6895 | **0.8544** | 0.4300 | 0.8286 | 0.000000 | 15 |
| Exploratory UMAP | HDBSCAN | 5 | **0.6122** | **0.7607** | 0.7475 | 0.6809 | 0.4270 | 0.000000 | 15 |

Table S10: Classification results with mean ± standard deviation over folds and repeats - Foraminifera Dataset

| Train Fraction | Classifier | Top-1 Accuracy | Top-2 Accuracy | Top-3 Accuracy | F1 Macro | F1 Weighted | Precision Macro | Recall Macro | Specificity Macro | Balanced Accuracy |
|---|---|---|---|---|---|---|---|---|---|---|
| 0.10 | Logistic Regression | 0.810 ± 0.016 | 0.917 ± 0.012 | 0.955 ± 0.008 | 0.671 ± 0.029 | 0.811 ± 0.015 | 0.683 ± 0.032 | 0.687 ± 0.026 | 0.994 ± 0.001 | 0.708 ± 0.022 |
| 0.10 | k-Nearest Neighbours | 0.711 ± 0.018 | 0.826 ± 0.015 | 0.872 ± 0.014 | 0.506 ± 0.029 | 0.690 ± 0.015 | 0.584 ± 0.041 | 0.488 ± 0.030 | 0.991 ± 0.001 | 0.493 ± 0.030 |
| 0.30 | Logistic Regression | 0.844 ± 0.009 | 0.939 ± 0.005 | 0.968 ± 0.004 | 0.716 ± 0.032 | 0.846 ± 0.009 | 0.717 ± 0.037 | 0.737 ± 0.034 | 0.995 ± 0.000 | 0.751 ± 0.034 |
| 0.30 | k-Nearest Neighbours | 0.756 ± 0.010 | 0.861 ± 0.008 | 0.899 ± 0.007 | 0.549 ± 0.028 | 0.745 ± 0.010 | 0.643 ± 0.041 | 0.517 ± 0.026 | 0.992 ± 0.000 | 0.518 ± 0.027 |
| 0.50 | Logistic Regression | 0.856 ± 0.007 | 0.947 ± 0.004 | 0.973 ± 0.003 | 0.726 ± 0.021 | 0.859 ± 0.007 | 0.715 ± 0.021 | 0.752 ± 0.025 | **0.996 ± 0.000** | 0.755 ± 0.024 |
| 0.50 | k-Nearest Neighbours | 0.777 ± 0.007 | 0.876 ± 0.006 | 0.912 ± 0.006 | 0.568 ± 0.017 | 0.768 ± 0.007 | 0.670 ± 0.027 | 0.530 ± 0.015 | 0.993 ± 0.000 | 0.530 ± 0.015 |
| 0.70 | Logistic Regression | 0.863 ± 0.005 | 0.951 ± 0.004 | 0.976 ± 0.003 | 0.740 ± 0.026 | 0.866 ± 0.005 | 0.724 ± 0.028 | 0.770 ± 0.028 | 0.996 ± 0.000 | 0.770 ± 0.028 |
| 0.70 | k-Nearest Neighbours | 0.787 ± 0.005 | 0.884 ± 0.004 | 0.917 ± 0.004 | 0.584 ± 0.022 | 0.779 ± 0.005 | 0.683 ± 0.029 | 0.546 ± 0.021 | 0.993 ± 0.000 | 0.546 ± 0.021 |
| 1.00 | Logistic Regression | **0.870 ± 0.004** | **0.956 ± 0.002** | **0.979 ± 0.002** | **0.770 ± 0.022** | **0.873 ± 0.004** | **0.748 ± 0.022** | **0.809 ± 0.026** | 0.996 ± 0.000 | **0.809 ± 0.026** |
| 1.00 | k-Nearest Neighbours | 0.797 ± 0.004 | 0.892 ± 0.003 | 0.924 ± 0.003 | 0.603 ± 0.018 | 0.790 ± 0.003 | 0.703 ± 0.024 | 0.564 ± 0.017 | 0.994 ± 0.000 | 0.564 ± 0.017 |

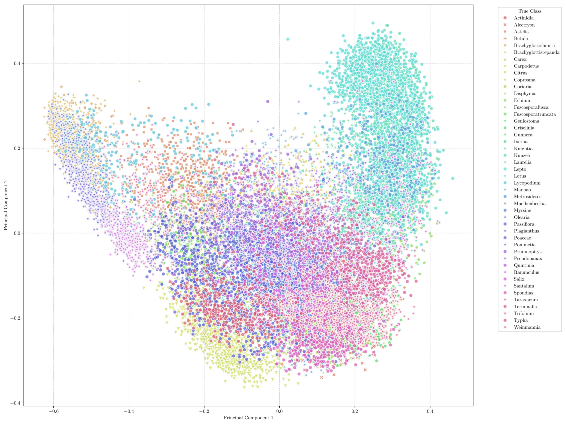

(a) Pollen

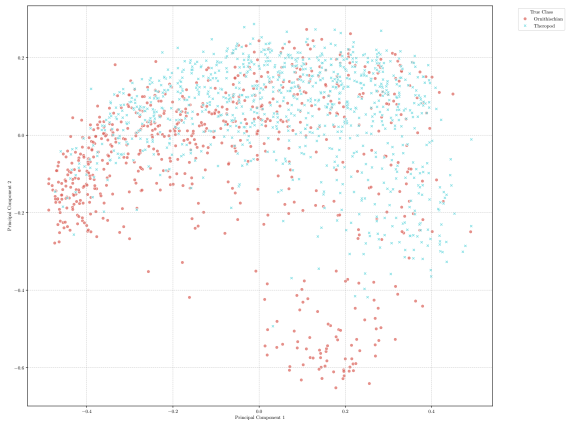

(b) Fossil Tracks

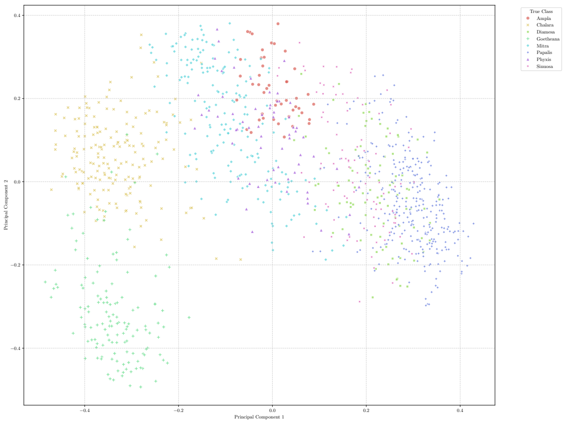

(c) Radiolaria

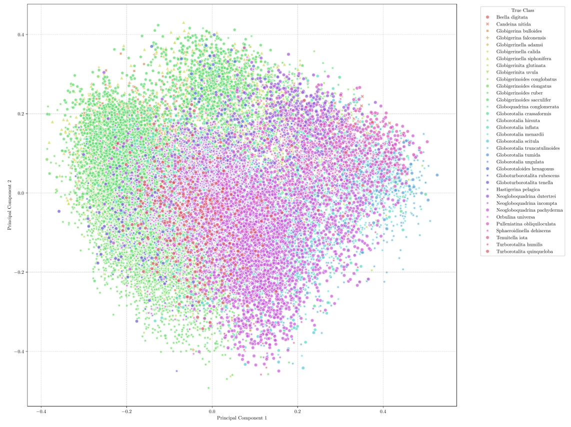

(d) Foraminifera

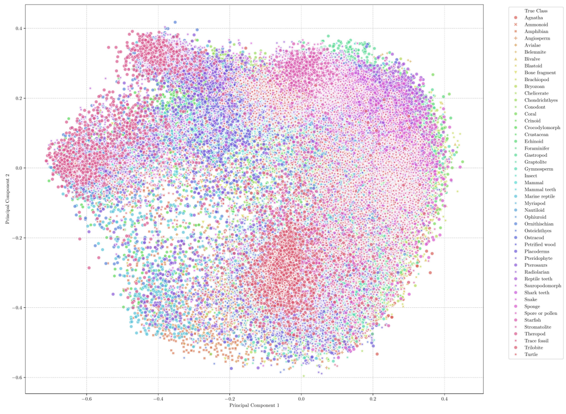

(e) Diverse Palaeontological Images

Figure S1: Dimension reduction of feature vectors by PCA for all Datasets and all Images. Points are labelled by the true class.

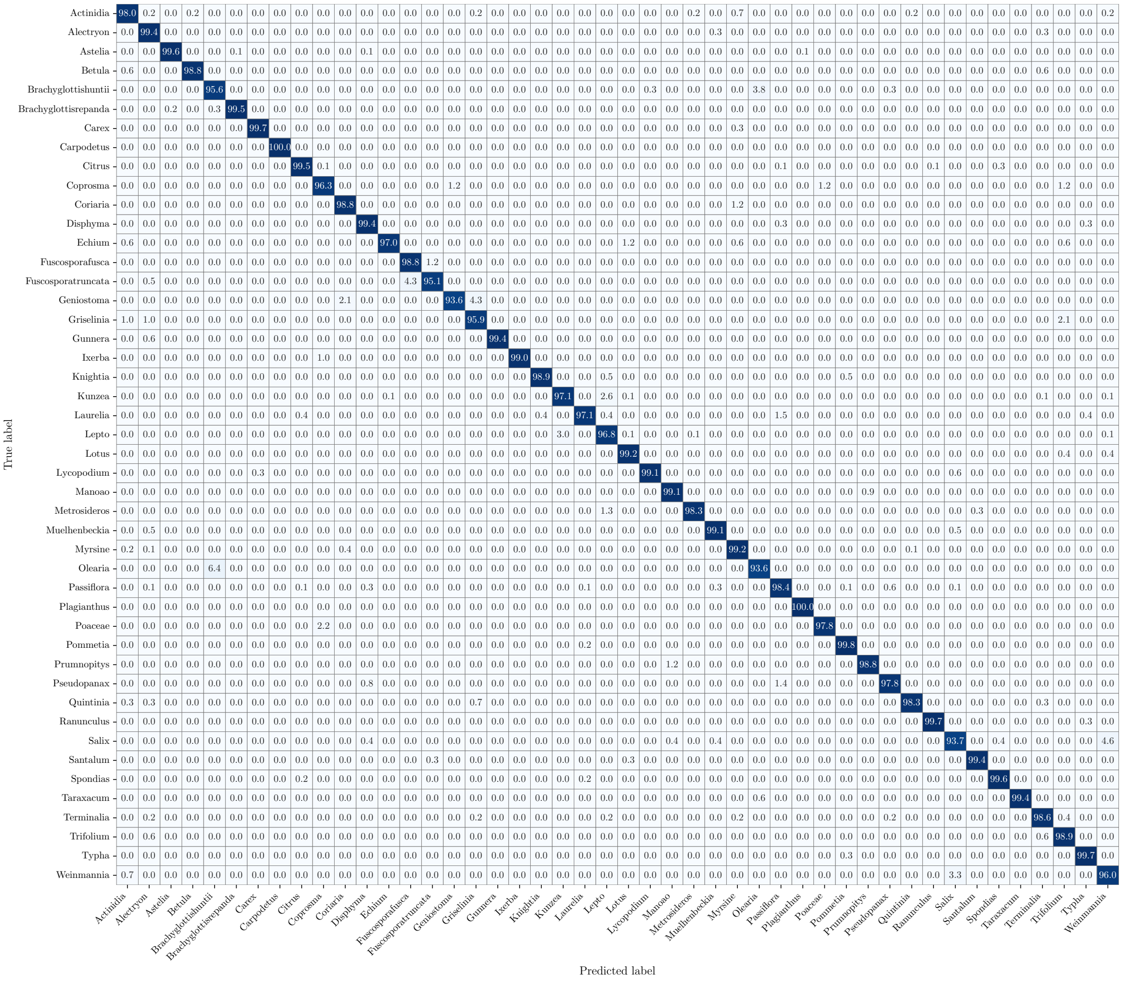

Figure S2: Confusion Matrix, Pollen Dataset

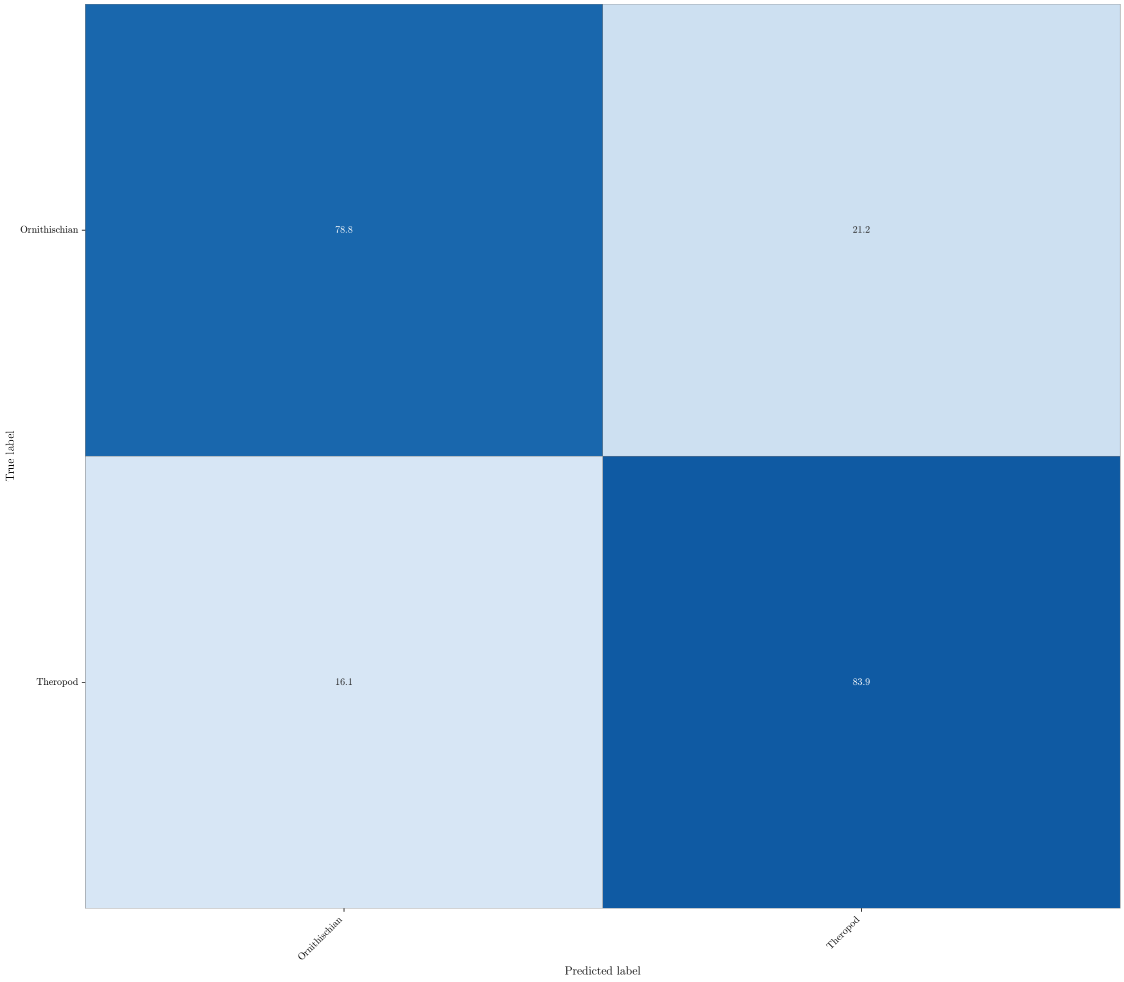

Figure S3: Confusion Matrix - Fossil Tracks Dataset

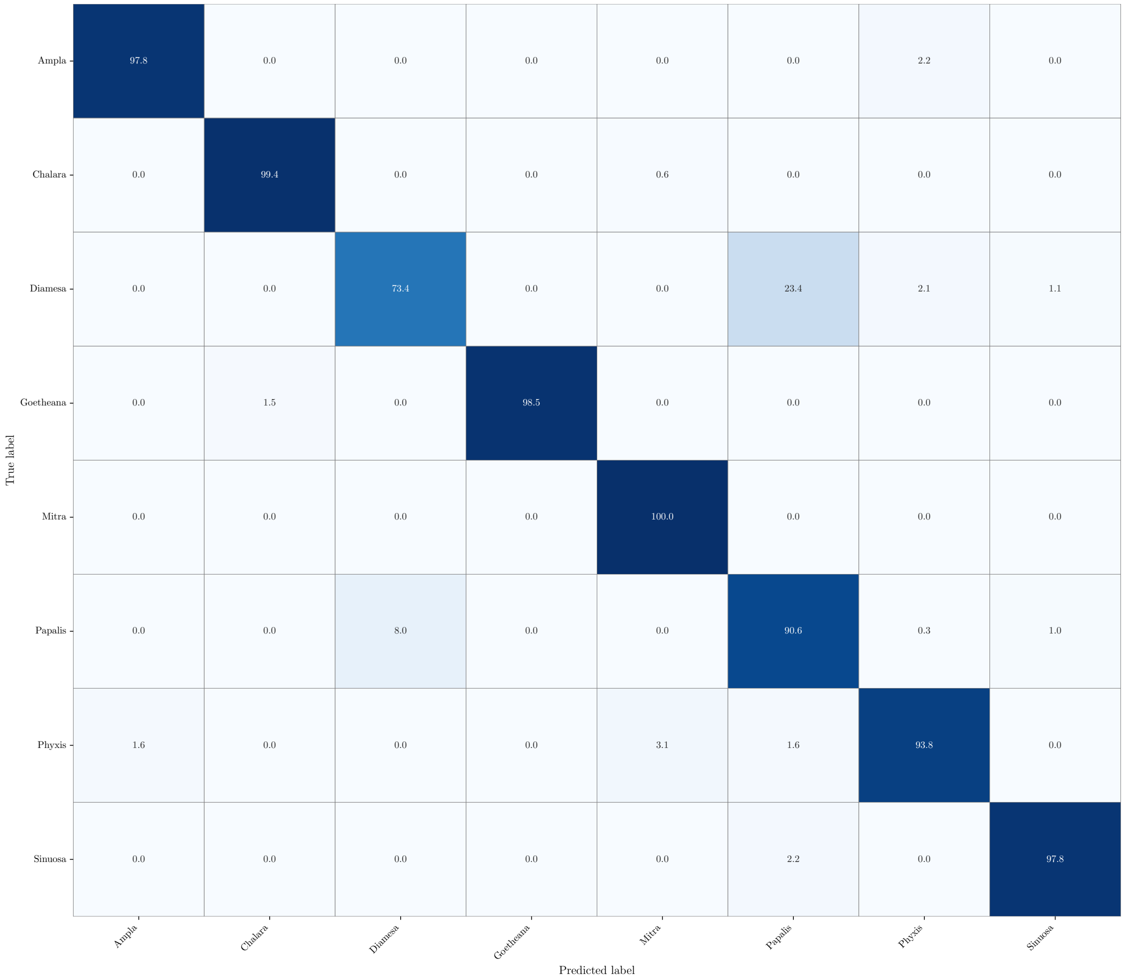

Figure S4: Confusion Matrix - Radiolarians Dataset

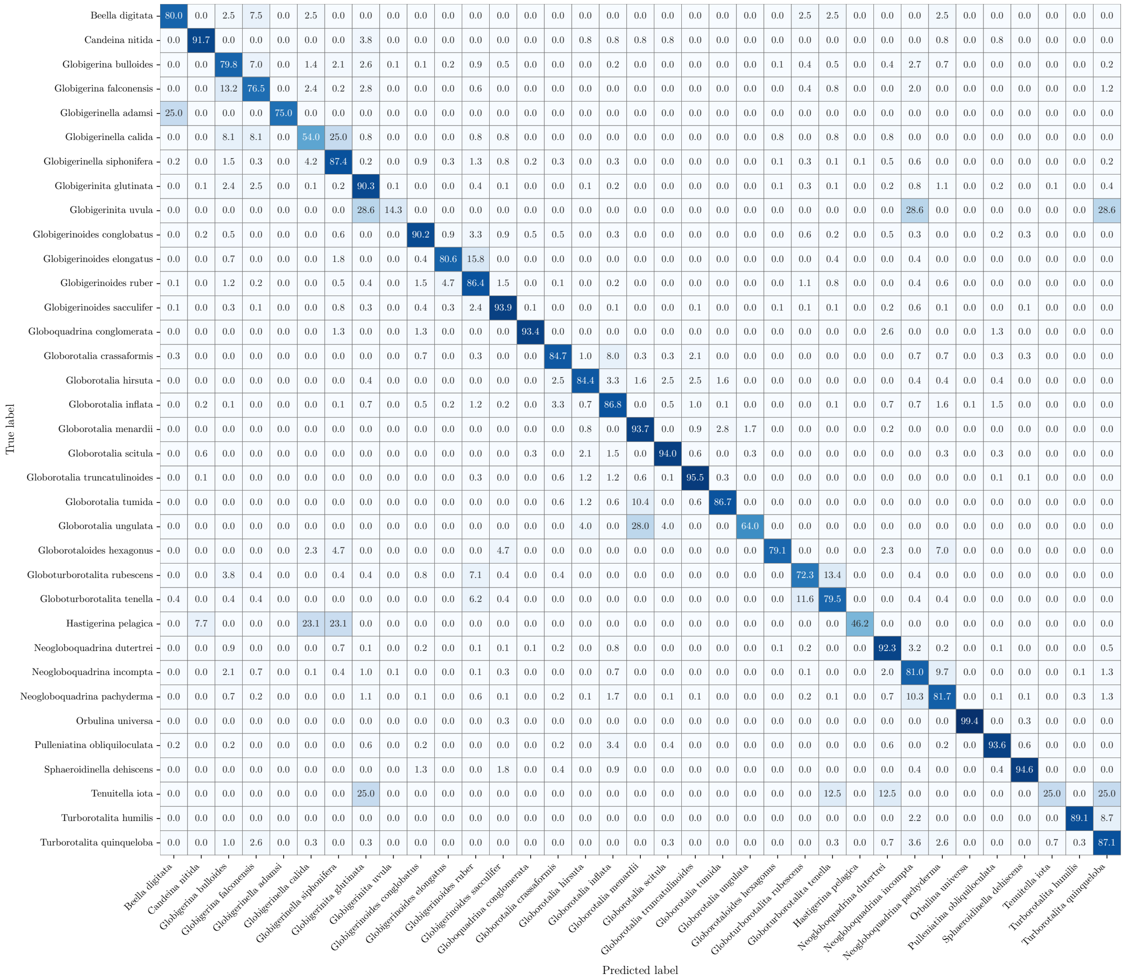

Figure S5: Confusion Matrix - Foraminifera

Table S11: Few-Shot Classification Results - Foraminifera

| Few-Shot k | Classifier | Repeats Successful | Top-1 Accuracy | Top-2 Accuracy | Top-3 Accuracy | F1 Macro | F1 Weighted | Precision Macro | Recall Macro | Specificity Macro | Balanced Accuracy |
|---|---|---|---|---|---|---|---|---|---|---|---|
| 1 | k-Nearest Neighbours | 50 | 0.218 ± 0.032 | 0.254 ± 0.030 | 0.384 ± 0.036 | 0.193 ± 0.016 | 0.238 ± 0.038 | 0.257 ± 0.022 | 0.305 ± 0.023 | 0.977 ± 0.001 | 0.305 ± 0.023 |
| 1 | Logistic Regression | 50 | 0.221 ± 0.033 | 0.338 ± 0.045 | 0.422 ± 0.050 | 0.194 ± 0.017 | 0.245 ± 0.042 | 0.235 ± 0.021 | 0.333 ± 0.021 | 0.977 ± 0.001 | 0.333 ± 0.021 |
| 2 | k-Nearest Neighbours | 50 | 0.224 ± 0.039 | 0.385 ± 0.047 | 0.470 ± 0.034 | 0.170 ± 0.018 | 0.217 ± 0.038 | 0.252 ± 0.024 | 0.279 ± 0.019 | 0.976 ± 0.001 | 0.279 ± 0.019 |
| 2 | Logistic Regression | 50 | 0.310 ± 0.032 | 0.457 ± 0.036 | 0.551 ± 0.036 | 0.265 ± 0.018 | 0.346 ± 0.039 | 0.289 ± 0.021 | 0.441 ± 0.023 | 0.980 ± 0.001 | 0.441 ± 0.023 |
| 3 | k-Nearest Neighbours | 50 | 0.225 ± 0.027 | 0.382 ± 0.032 | 0.512 ± 0.035 | 0.197 ± 0.015 | 0.223 ± 0.035 | 0.289 ± 0.019 | 0.330 ± 0.019 | 0.977 ± 0.001 | 0.330 ± 0.019 |
| 3 | Logistic Regression | 50 | 0.364 ± 0.031 | 0.518 ± 0.037 | 0.614 ± 0.036 | 0.307 ± 0.014 | 0.401 ± 0.036 | 0.323 ± 0.015 | 0.506 ± 0.016 | 0.981 ± 0.001 | 0.506 ± 0.016 |
| 5 | k-Nearest Neighbours | 50 | 0.278 ± 0.029 | 0.439 ± 0.030 | 0.556 ± 0.031 | 0.251 ± 0.015 | 0.304 ± 0.038 | 0.315 ± 0.015 | 0.402 ± 0.022 | 0.979 ± 0.001 | 0.402 ± 0.022 |
| 5 | Logistic Regression | 50 | 0.433 ± 0.028 | 0.599 ± 0.028 | 0.693 ± 0.025 | 0.367 ± 0.013 | 0.475 ± 0.030 | 0.371 ± 0.013 | 0.584 ± 0.018 | 0.983 ± 0.001 | 0.584 ± 0.018 |
| 10 | k-Nearest Neighbours | 50 | 0.358 ± 0.025 | 0.528 ± 0.027 | 0.638 ± 0.026 | 0.321 ± 0.014 | 0.394 ± 0.031 | 0.358 ± 0.013 | 0.499 ± 0.024 | 0.981 ± 0.001 | 0.499 ± 0.024 |
| 10 | Logistic Regression | 50 | 0.534 ± 0.018 | 0.702 ± 0.016 | 0.786 ± 0.014 | 0.437 ± 0.011 | 0.575 ± 0.017 | 0.423 ± 0.010 | 0.669 ± 0.020 | 0.986 ± 0.001 | 0.669 ± 0.020 |
| 20 | k-Nearest Neighbours | 50 | 0.435 ± 0.017 | 0.605 ± 0.015 | 0.706 ± 0.014 | 0.372 ± 0.016 | 0.473 ± 0.020 | 0.391 ± 0.015 | 0.559 ± 0.025 | 0.983 ± 0.001 | 0.559 ± 0.025 |
| 20 | Logistic Regression | 50 | **0.616 ± 0.014** | **0.775 ± 0.013** | **0.848 ± 0.011** | **0.489 ± 0.012** | **0.650 ± 0.014** | **0.465 ± 0.013** | **0.723 ± 0.022** | **0.989 ± 0.000** | **0.723 ± 0.022** |

Table S12: Clustering Results - Foraminifera

| Pipeline | Algorithm | No. of Clusters | ARI | AMI | Purity | Silhouette | DBI | Noise Fraction | Components Used |
|---|---|---|---|---|---|---|---|---|---|
| PCA | KMeans (Known K) | 35 | 0.1788 | 0.4256 | 0.5771 | 0.0711 | 2.5075 | 0.000000 | 50 |
| PCA | KMeans (Unknown K) | 2 | 0.0923 | 0.1556 | 0.2665 | 0.1153 | 2.7967 | 0.000000 | 50 |
| PCA | Bayesian GMM | 50 | 0.1846 | 0.4871 | 0.6574 | 0.0379 | 2.8144 | 0.000000 | 50 |
| PCA | HDBSCAN | 0 | 0.0000 | 0.0000 | 0.0000 | nan | nan | 1.000000 | 50 |
| Exploratory UMAP | KMeans (Known K) | 35 | 0.2505 | 0.5177 | **0.6643** | 0.4040 | 0.8222 | 0.000000 | 15 |
| Exploratory UMAP | KMeans (Unknown K) | 2 | 0.0466 | 0.0951 | 0.2259 | **0.7883** | **0.3012** | 0.000000 | 15 |
| Exploratory UMAP | Bayesian GMM | 38 | **0.4172** | **0.5456** | 0.5886 | -0.0011 | 1.2859 | 0.000000 | 15 |
| Exploratory UMAP | HDBSCAN | 3 | 0.0560 | 0.1359 | 0.2419 | 0.6234 | 0.3518 | 0.006744 | 15 |

Table S13: Classification Results with mean ± standard deviation over folds and repeats - RFID

| Train Fraction | Classifier | Top-1 Accuracy | Top-2 Accuracy | Top-3 Accuracy | F1 Macro | F1 Weighted | Precision Macro | Recall Macro | Specificity Macro | Balanced Accuracy |
|---|---|---|---|---|---|---|---|---|---|---|
| 0.10 | Logistic Regression | 0.788 ± 0.010 | 0.880 ± 0.009 | 0.916 ± 0.008 | 0.790 ± 0.010 | 0.789 ± 0.010 | 0.798 ± 0.010 | 0.789 ± 0.010 | 0.996 ± 0.000 | 0.789 ± 0.010 |
| 0.10 | k-Nearest Neighbours | 0.734 ± 0.013 | 0.827 ± 0.010 | 0.864 ± 0.010 | 0.736 ± 0.011 | 0.735 ± 0.011 | 0.753 ± 0.011 | 0.736 ± 0.012 | 0.995 ± 0.000 | 0.736 ± 0.012 |
| 0.30 | Logistic Regression | 0.817 ± 0.007 | 0.900 ± 0.005 | 0.931 ± 0.004 | 0.818 ± 0.006 | 0.817 ± 0.006 | 0.822 ± 0.006 | 0.817 ± 0.007 | 0.996 ± 0.000 | 0.817 ± 0.007 |
| 0.30 | k-Nearest Neighbours | 0.788 ± 0.008 | 0.866 ± 0.006 | 0.896 ± 0.006 | 0.790 ± 0.007 | 0.789 ± 0.007 | 0.798 ± 0.007 | 0.790 ± 0.007 | 0.996 ± 0.000 | 0.790 ± 0.007 |
| 0.50 | Logistic Regression | 0.825 ± 0.006 | 0.905 ± 0.005 | 0.935 ± 0.004 | 0.827 ± 0.006 | 0.826 ± 0.006 | 0.829 ± 0.005 | 0.826 ± 0.006 | 0.996 ± 0.000 | 0.826 ± 0.006 |
| 0.50 | k-Nearest Neighbours | 0.809 ± 0.005 | 0.881 ± 0.004 | 0.908 ± 0.003 | 0.811 ± 0.005 | 0.810 ± 0.005 | 0.817 ± 0.005 | 0.810 ± 0.005 | 0.996 ± 0.000 | 0.810 ± 0.005 |
| 0.70 | Logistic Regression | 0.829 ± 0.003 | 0.908 ± 0.003 | 0.938 ± 0.002 | 0.831 ± 0.003 | 0.830 ± 0.003 | 0.833 ± 0.003 | 0.830 ± 0.003 | **0.997 ± 0.000** | 0.830 ± 0.003 |
| 0.70 | k-Nearest Neighbours | 0.821 ± 0.004 | 0.890 ± 0.004 | 0.915 ± 0.003 | 0.824 ± 0.004 | 0.822 ± 0.004 | 0.828 ± 0.004 | 0.823 ± 0.004 | 0.996 ± 0.000 | 0.823 ± 0.004 |
| 1.00 | Logistic Regression | 0.833 ± 0.003 | **0.911 ± 0.003** | **0.941 ± 0.002** | 0.835 ± 0.003 | 0.834 ± 0.003 | 0.836 ± 0.003 | 0.834 ± 0.003 | 0.997 ± 0.000 | 0.834 ± 0.003 |
| 1.00 | k-Nearest Neighbours | **0.834 ± 0.003** | 0.899 ± 0.003 | 0.922 ± 0.002 | **0.836 ± 0.003** | **0.835 ± 0.003** | **0.840 ± 0.003** | **0.836 ± 0.003** | 0.997 ± 0.000 | **0.836 ± 0.003** |

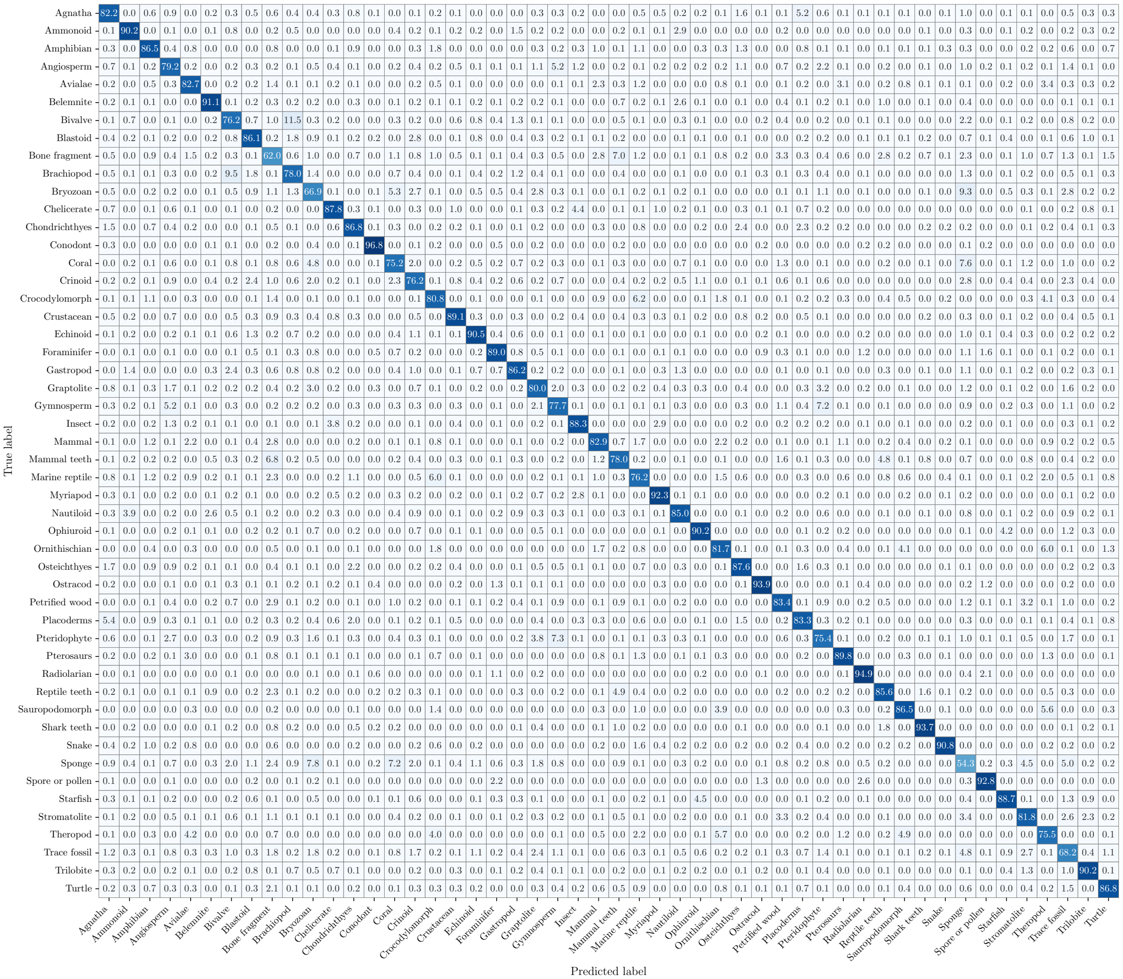

Figure S6: Confusion Matrix - RFID

Table S14: Few-Shot Classification Results - RFID

| Few-Shot k | Classifier | Repeats Successful | Top-1 Accuracy | Top-2 Accuracy | Top-3 Accuracy | F1 Macro | F1 Weighted | Precision Macro | Recall Macro | Specificity Macro | Balanced Accuracy |
|---|---|---|---|---|---|---|---|---|---|---|---|
| 1 | k-Nearest Neighbours | 50 | 0.343 ± 0.028 | 0.357 ± 0.027 | 0.376 ± 0.027 | 0.333 ± 0.028 | 0.334 ± 0.028 | 0.401 ± 0.033 | 0.342 ± 0.028 | 0.987 ± 0.001 | 0.342 ± 0.028 |
| 1 | Logistic Regression | 50 | 0.355 ± 0.025 | 0.473 ± 0.028 | 0.544 ± 0.028 | 0.346 ± 0.025 | 0.346 ± 0.025 | 0.393 ± 0.027 | 0.355 ± 0.024 | 0.987 ± 0.001 | 0.355 ± 0.024 |
| 2 | k-Nearest Neighbours | 50 | 0.345 ± 0.019 | 0.537 ± 0.019 | 0.549 ± 0.018 | 0.321 ± 0.023 | 0.323 ± 0.022 | 0.401 ± 0.025 | 0.336 ± 0.022 | 0.987 ± 0.000 | 0.336 ± 0.022 |
| 2 | Logistic Regression | 50 | 0.461 ± 0.018 | 0.591 ± 0.017 | 0.663 ± 0.017 | 0.455 ± 0.020 | 0.456 ± 0.020 | 0.486 ± 0.022 | 0.461 ± 0.018 | 0.989 ± 0.000 | 0.461 ± 0.018 |
| 3 | k-Nearest Neighbours | 50 | 0.390 ± 0.018 | 0.557 ± 0.017 | 0.652 ± 0.017 | 0.381 ± 0.019 | 0.381 ± 0.019 | 0.476 ± 0.022 | 0.389 ± 0.018 | 0.988 ± 0.000 | 0.389 ± 0.018 |
| 3 | Logistic Regression | 50 | 0.520 ± 0.015 | 0.653 ± 0.015 | 0.720 ± 0.014 | 0.518 ± 0.016 | 0.517 ± 0.016 | 0.542 ± 0.016 | 0.521 ± 0.015 | 0.990 ± 0.000 | 0.521 ± 0.015 |
| 5 | k-Nearest Neighbours | 50 | 0.458 ± 0.012 | 0.601 ± 0.012 | 0.685 ± 0.012 | 0.456 ± 0.013 | 0.456 ± 0.013 | 0.522 ± 0.013 | 0.461 ± 0.012 | 0.989 ± 0.000 | 0.461 ± 0.012 |
| 5 | Logistic Regression | 50 | 0.587 ± 0.013 | 0.717 ± 0.011 | 0.780 ± 0.010 | 0.587 ± 0.014 | 0.586 ± 0.014 | 0.603 ± 0.015 | 0.588 ± 0.013 | 0.992 ± 0.000 | 0.588 ± 0.013 |
| 10 | k-Nearest Neighbours | 50 | 0.548 ± 0.009 | 0.681 ± 0.008 | 0.750 ± 0.008 | 0.553 ± 0.009 | 0.552 ± 0.009 | 0.594 ± 0.010 | 0.554 ± 0.009 | 0.991 ± 0.000 | 0.554 ± 0.009 |
| 10 | Logistic Regression | 50 | 0.663 ± 0.006 | 0.782 ± 0.005 | 0.838 ± 0.005 | 0.665 ± 0.006 | 0.664 ± 0.006 | 0.674 ± 0.007 | 0.664 ± 0.006 | 0.993 ± 0.000 | 0.664 ± 0.006 |
| 20 | k-Nearest Neighbours | 50 | 0.622 ± 0.007 | 0.740 ± 0.005 | 0.797 ± 0.005 | 0.628 ± 0.007 | 0.626 ± 0.007 | 0.652 ± 0.006 | 0.626 ± 0.007 | 0.992 ± 0.000 | 0.626 ± 0.007 |
| 20 | Logistic Regression | 50 | **0.717 ± 0.005** | **0.826 ± 0.004** | **0.873 ± 0.004** | **0.719 ± 0.005** | **0.718 ± 0.005** | **0.726 ± 0.006** | **0.718 ± 0.005** | **0.994 ± 0.000** | **0.718 ± 0.005** |

Table S15: Clustering results - RFID

| Pipeline | Algorithm | No. of Clusters | ARI | AMI | Purity | Silhouette | DBI | Noise Fraction | Components Used |
|---|---|---|---|---|---|---|---|---|---|
| PCA | KMeans (Known K) | 50 | 0.3148 | 0.5708 | 0.4789 | 0.1649 | 2.0605 | 0.000000 | 50 |
| PCA | KMeans (Unknown K) | 39 | 0.2777 | 0.5543 | 0.4198 | 0.1693 | 1.9669 | 0.000000 | 50 |
| PCA | Bayesian GMM | 50 | 0.3092 | 0.5910 | 0.4713 | 0.1262 | 2.2057 | 0.000000 | 50 |
| PCA | HDBSCAN | 0 | 0.0000 | 0.0000 | 0.0000 | nan | nan | 1.000000 | 50 |
| Exploratory UMAP | KMeans (Known K) | 50 | **0.3571** | 0.6390 | 0.5051 | 0.5636 | 0.5652 | 0.000000 | 15 |
| Exploratory UMAP | KMeans (Unknown K) | 37 | 0.3195 | 0.6245 | 0.4353 | **0.5847** | 0.5169 | 0.000000 | 15 |
| Exploratory UMAP | Bayesian GMM | 50 | 0.3497 | **0.6440** | **0.5057** | 0.5197 | 0.9864 | 0.000000 | 15 |
| Exploratory UMAP | HDBSCAN | 7 | 0.0684 | 0.4328 | 0.1355 | 0.5587 | **0.4859** | 0.093309 | 15 |

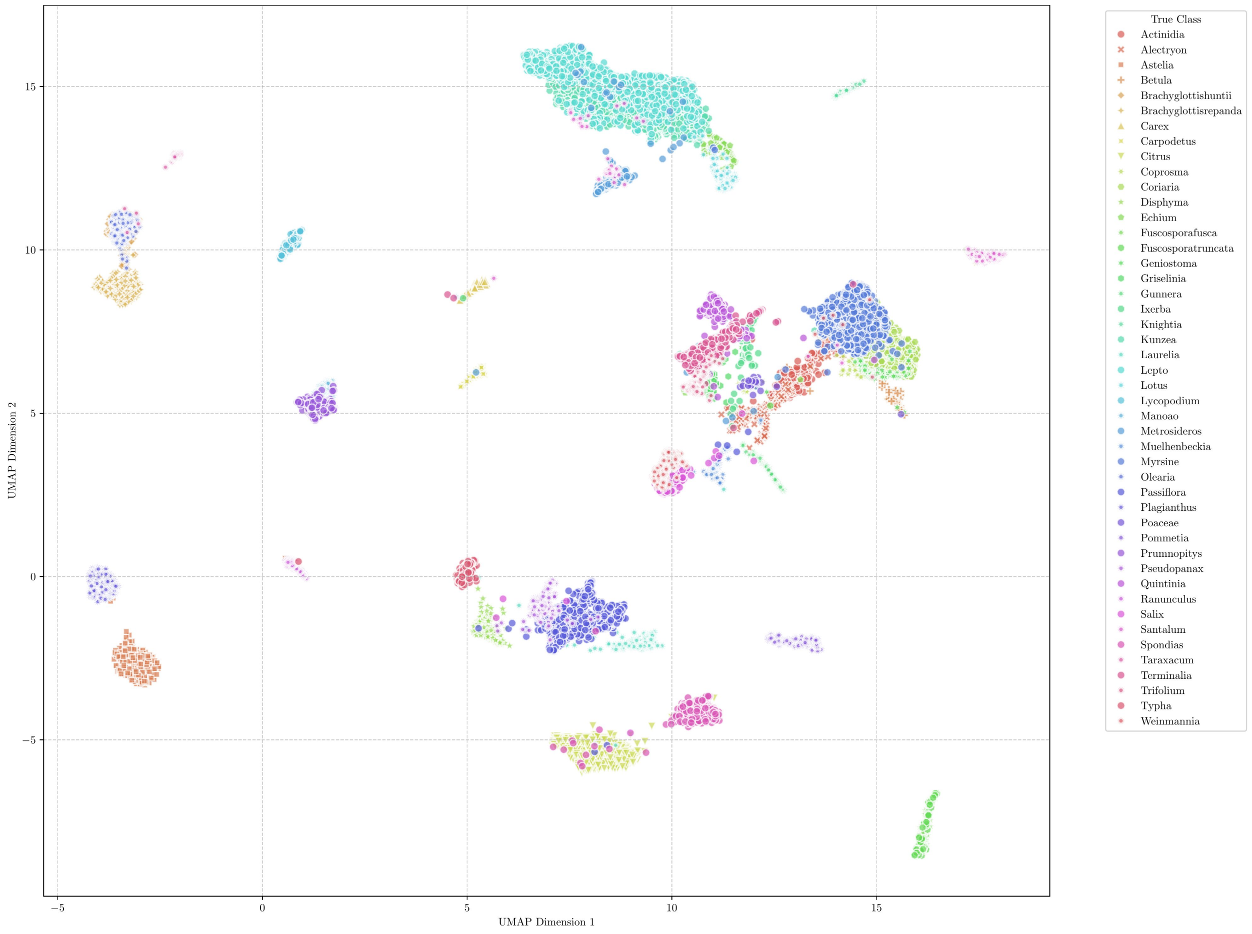

Figure S7: Pollen - UMAP

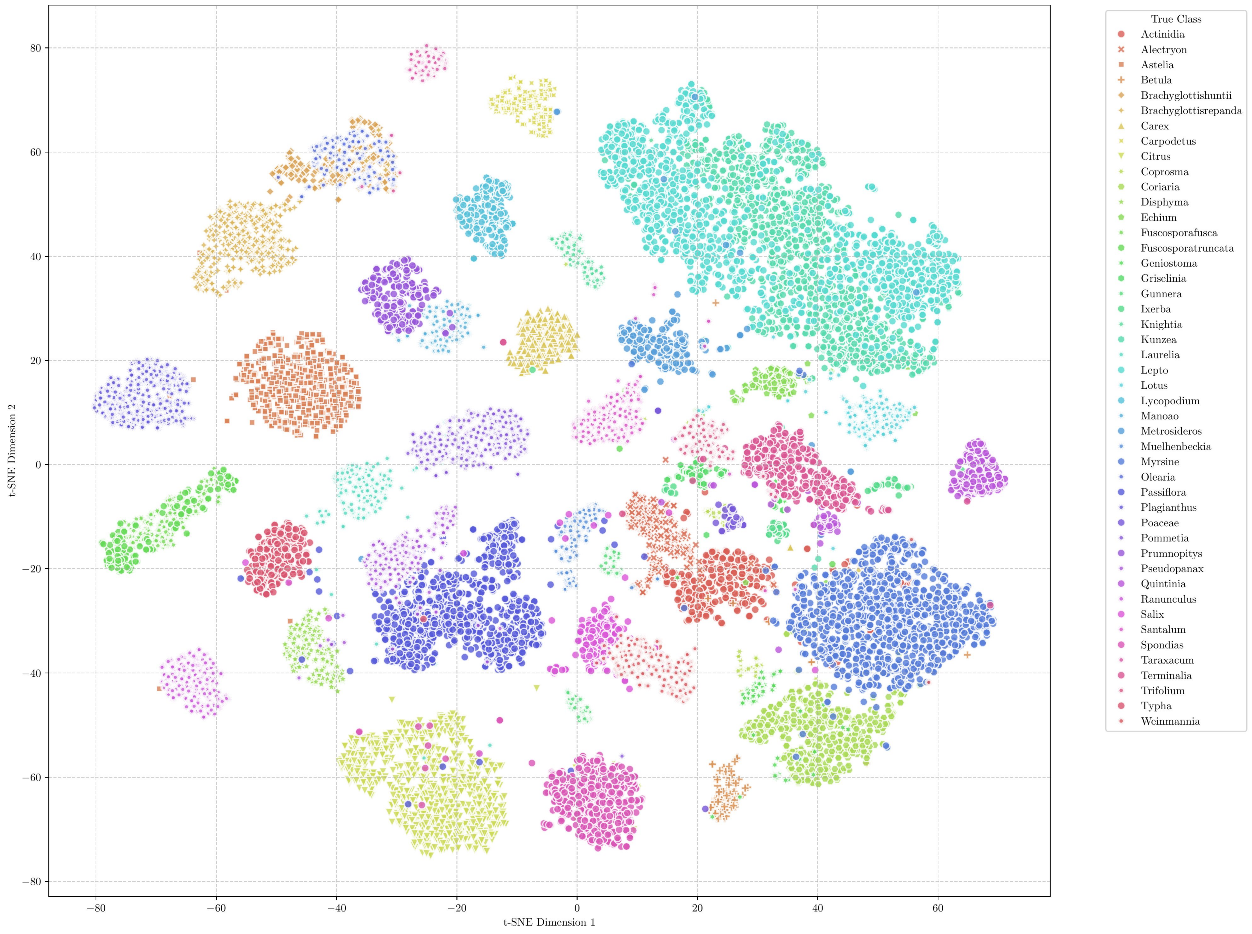

Figure S8: Pollen - t-SNE

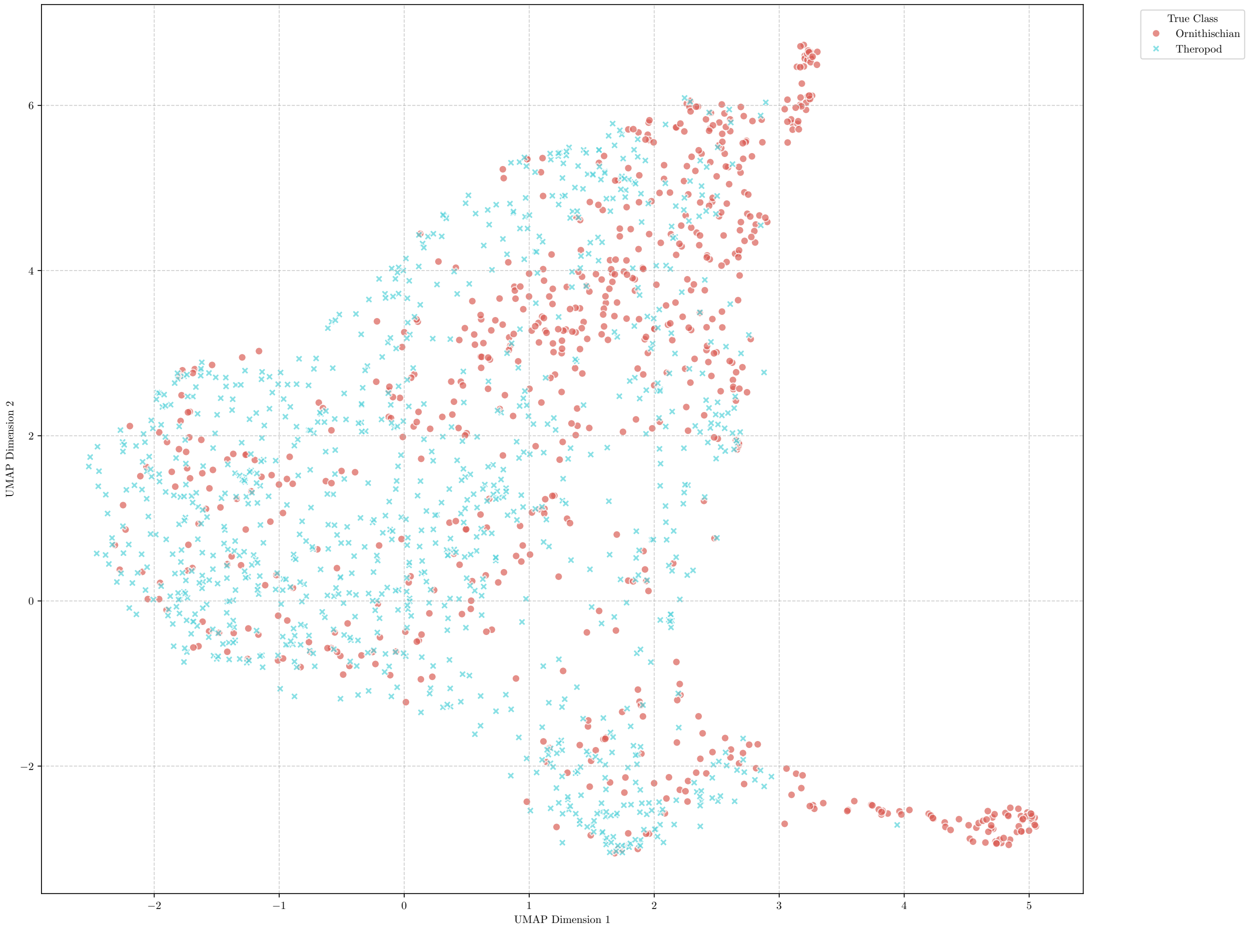

Figure S9: Fossil Tracks - UMAP

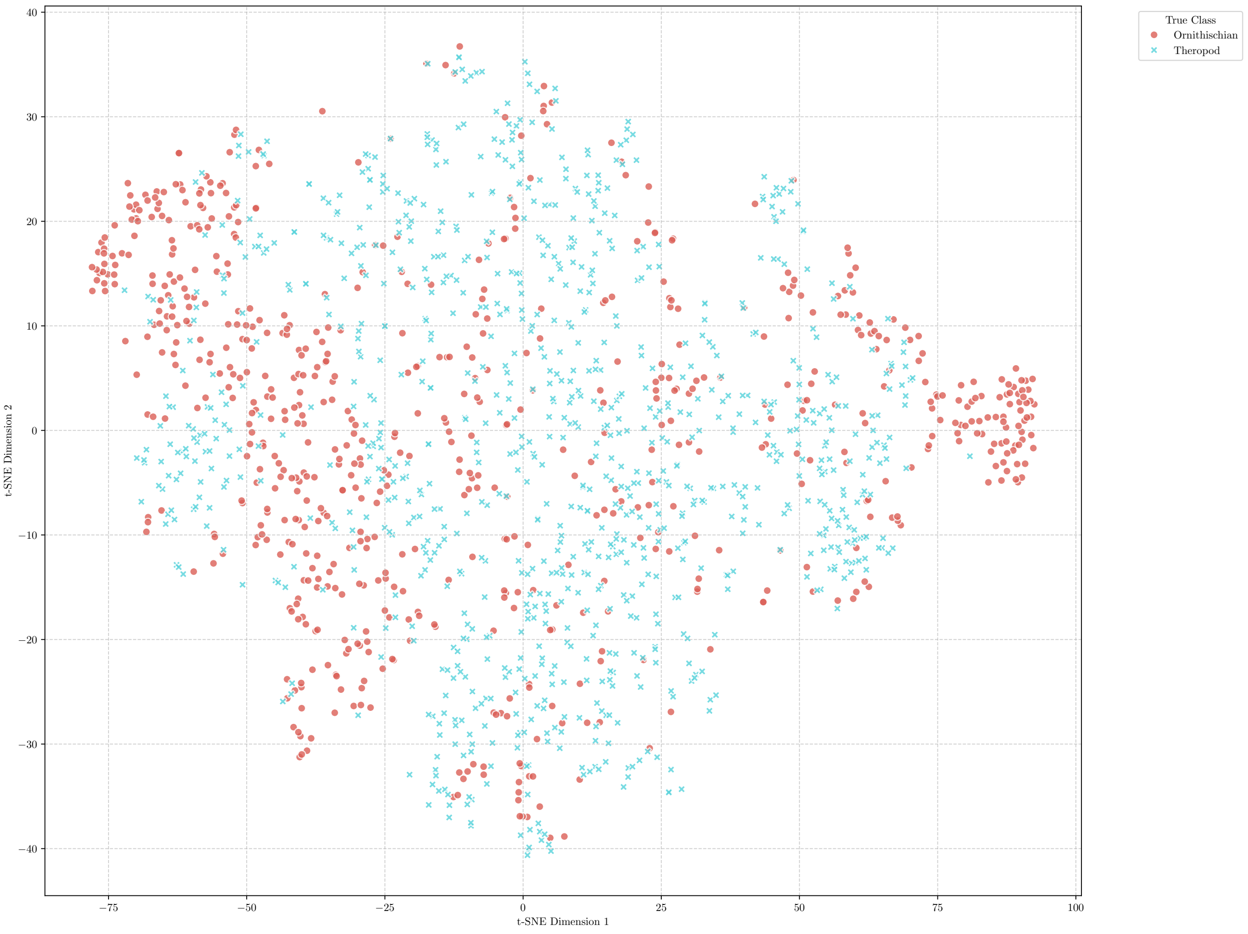

Figure S10: Fossil Tracks - t-SNE

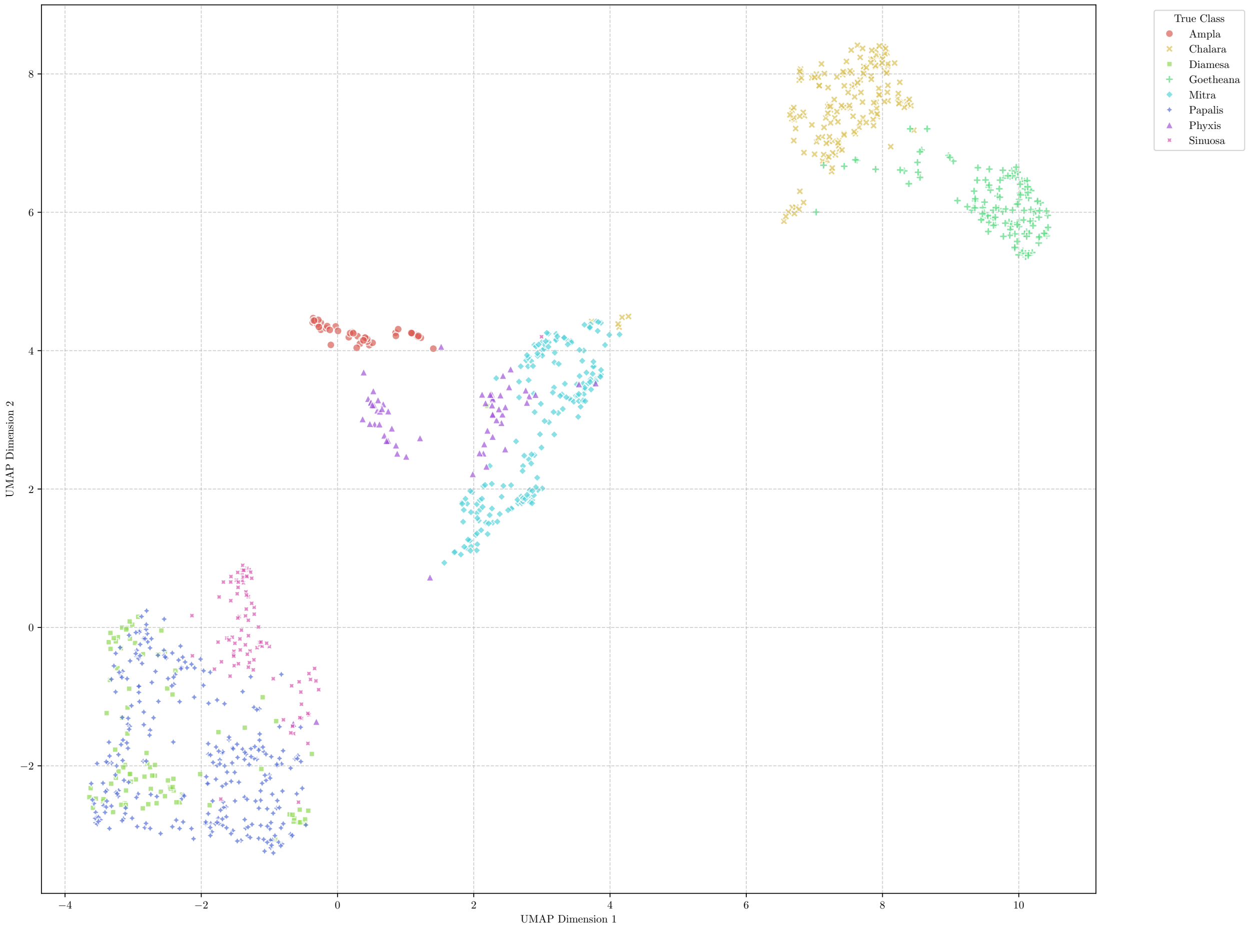

Figure S11: Radiolaria - UMAP

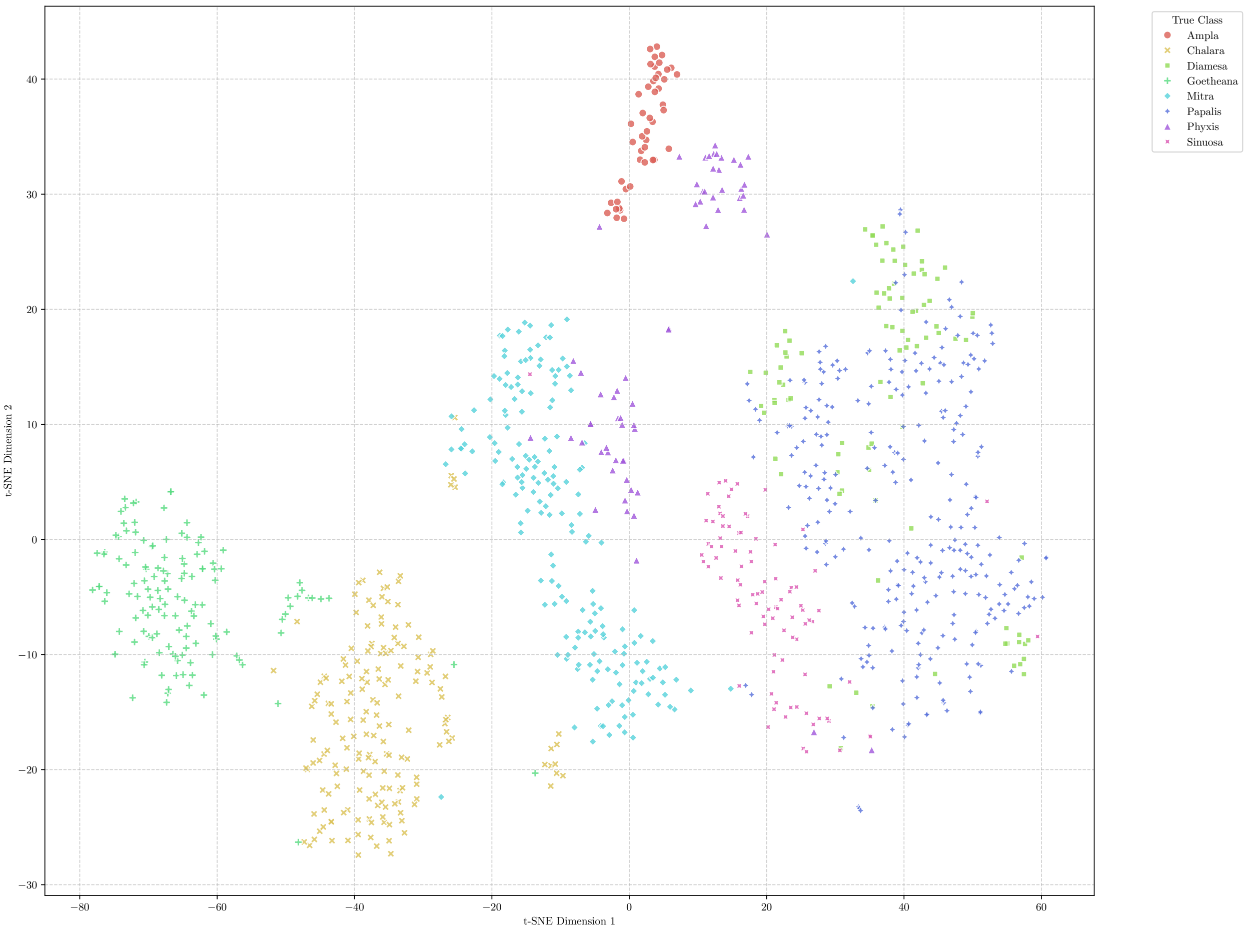

Figure S12: Radiolaria - t-SNE

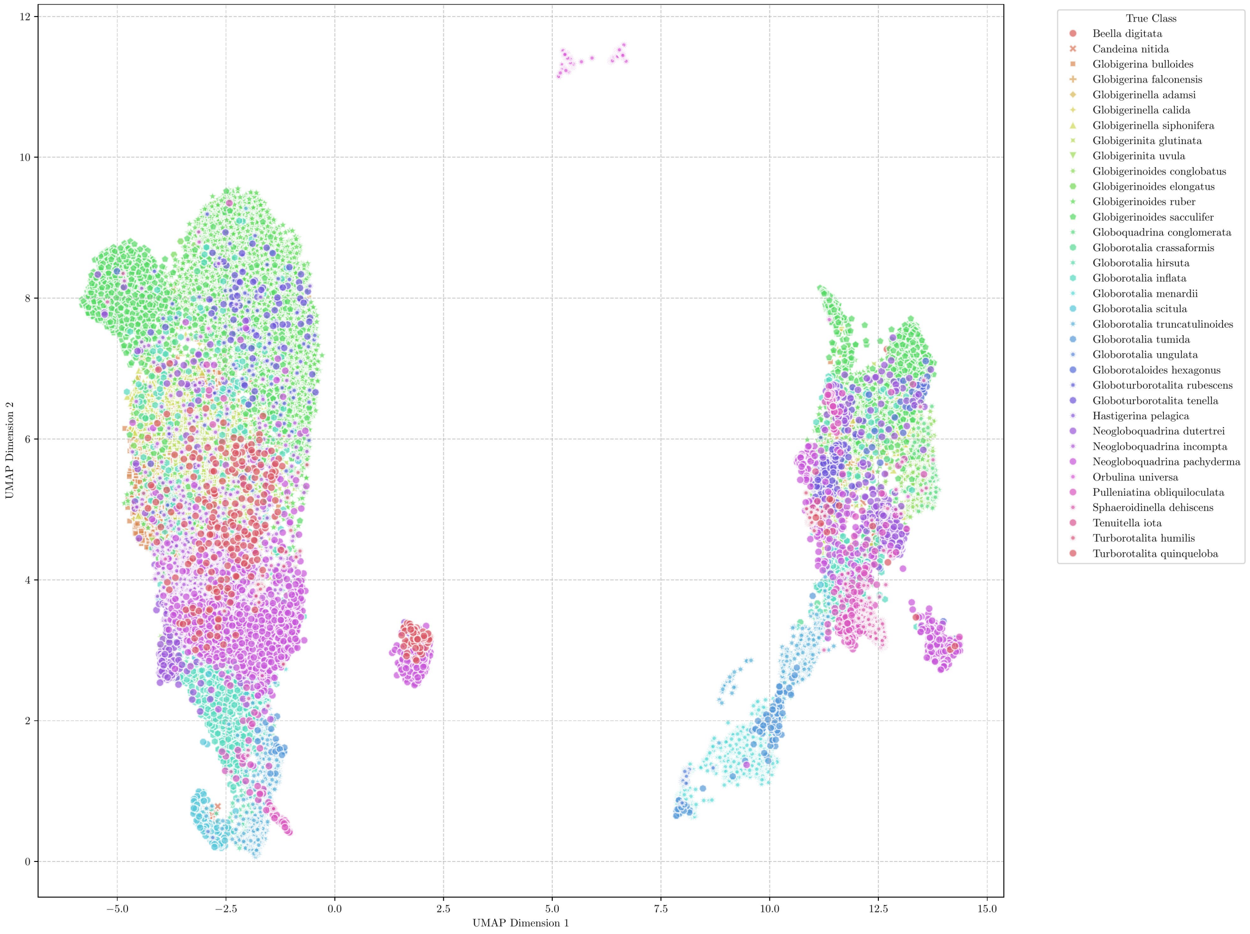

Figure S13: Foraminifera - UMAP

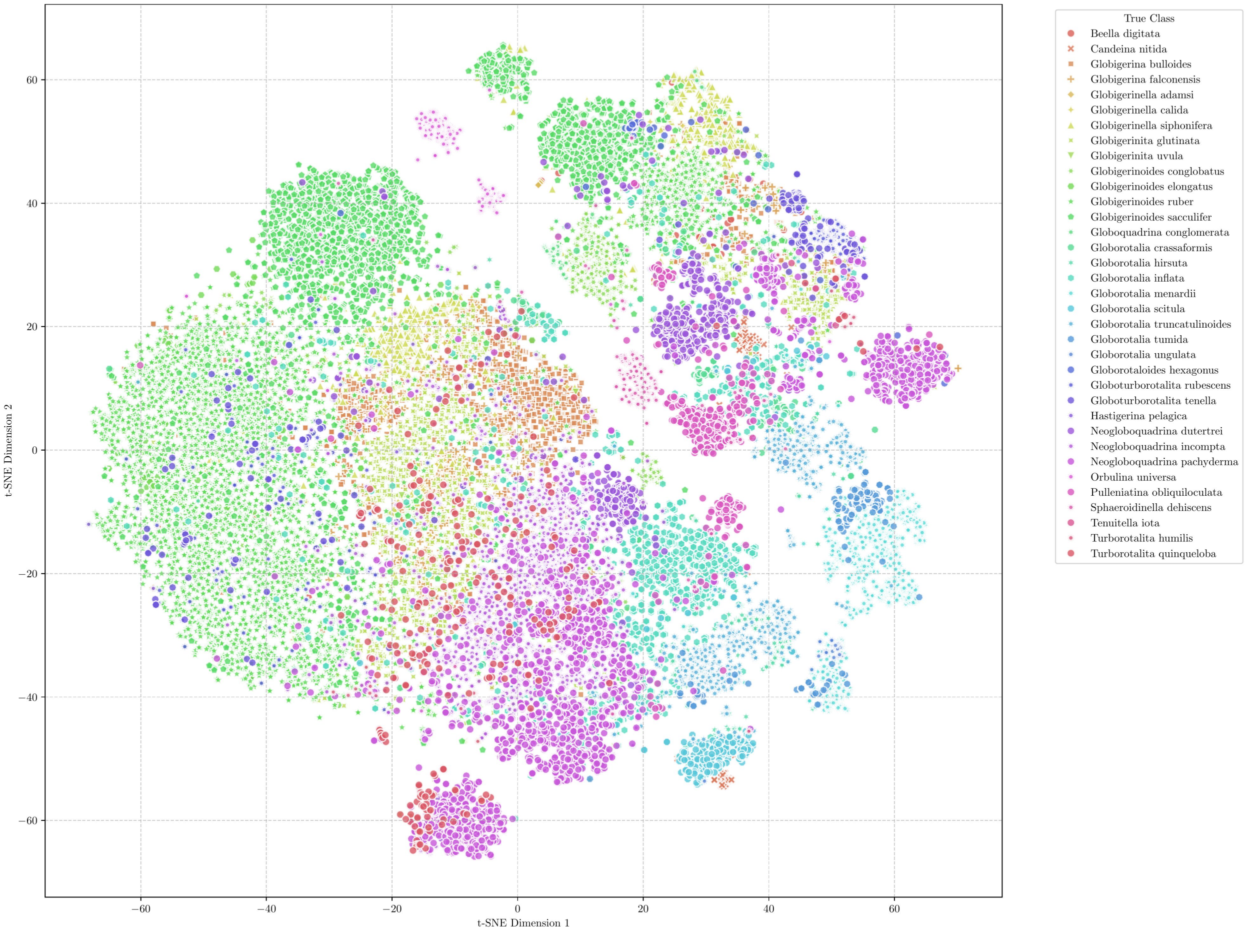

Figure S14: Foraminifera - t-SNE

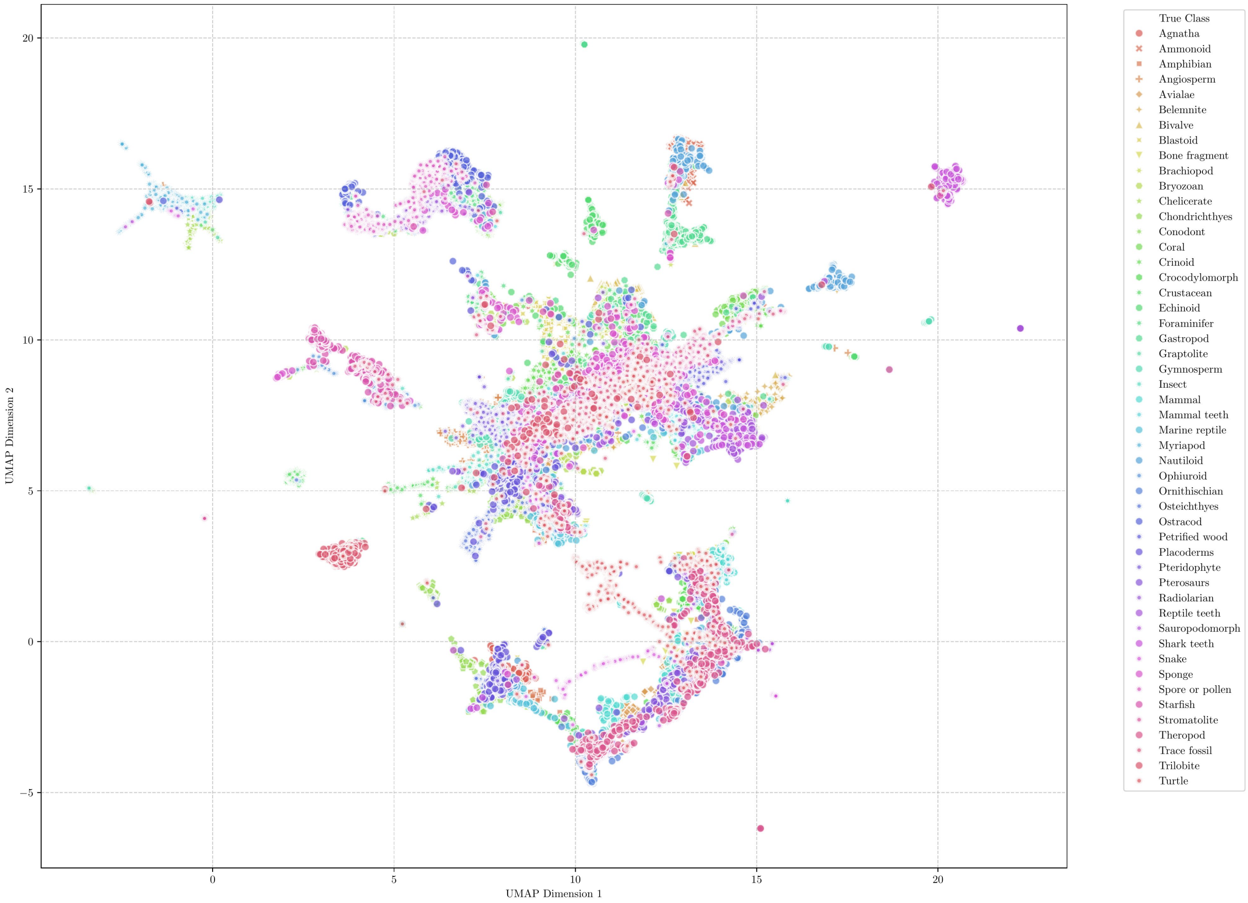

Figure S15: Diverse Palaeontological images (RFID) - UMAP

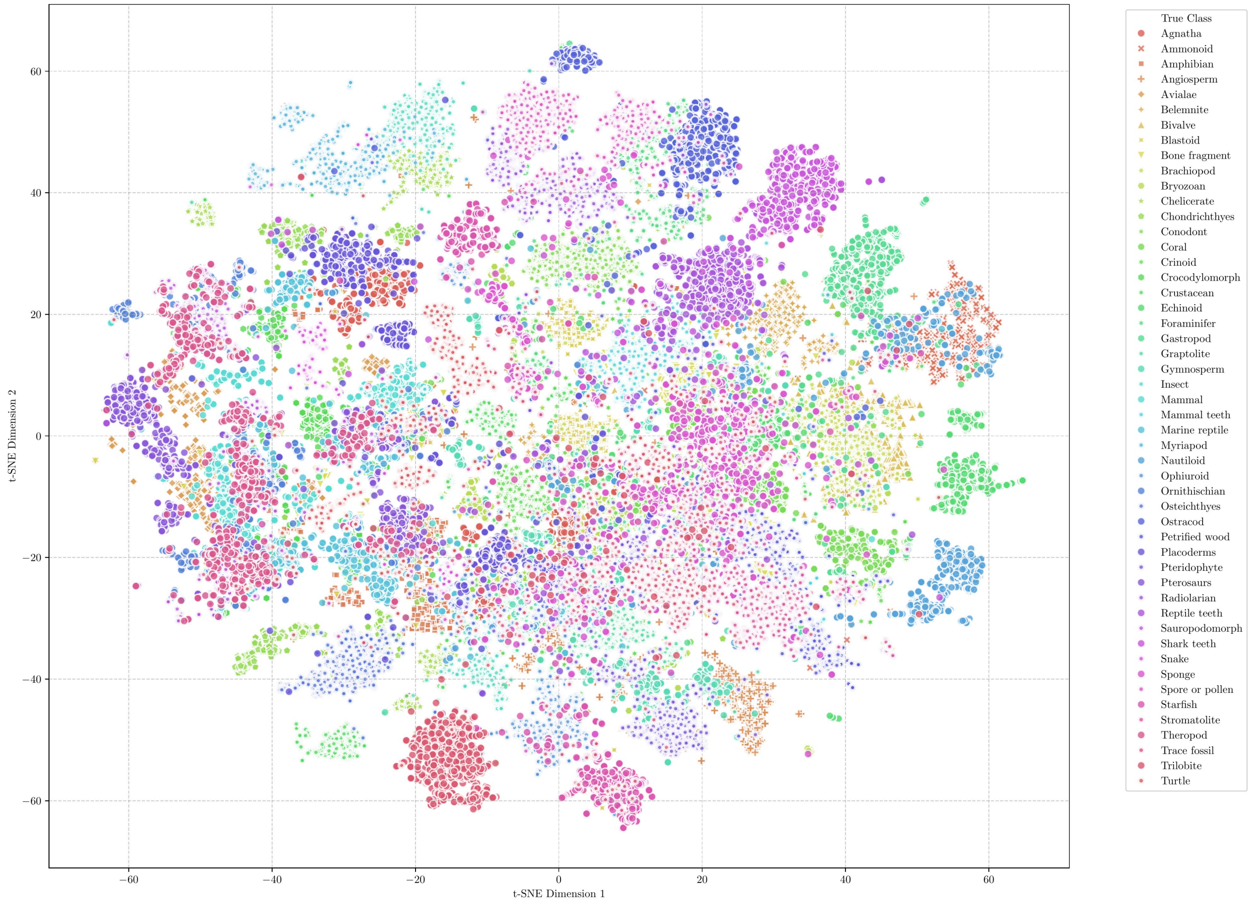

Figure S16: Diverse Palaeontological images (RFID) - t-SNE

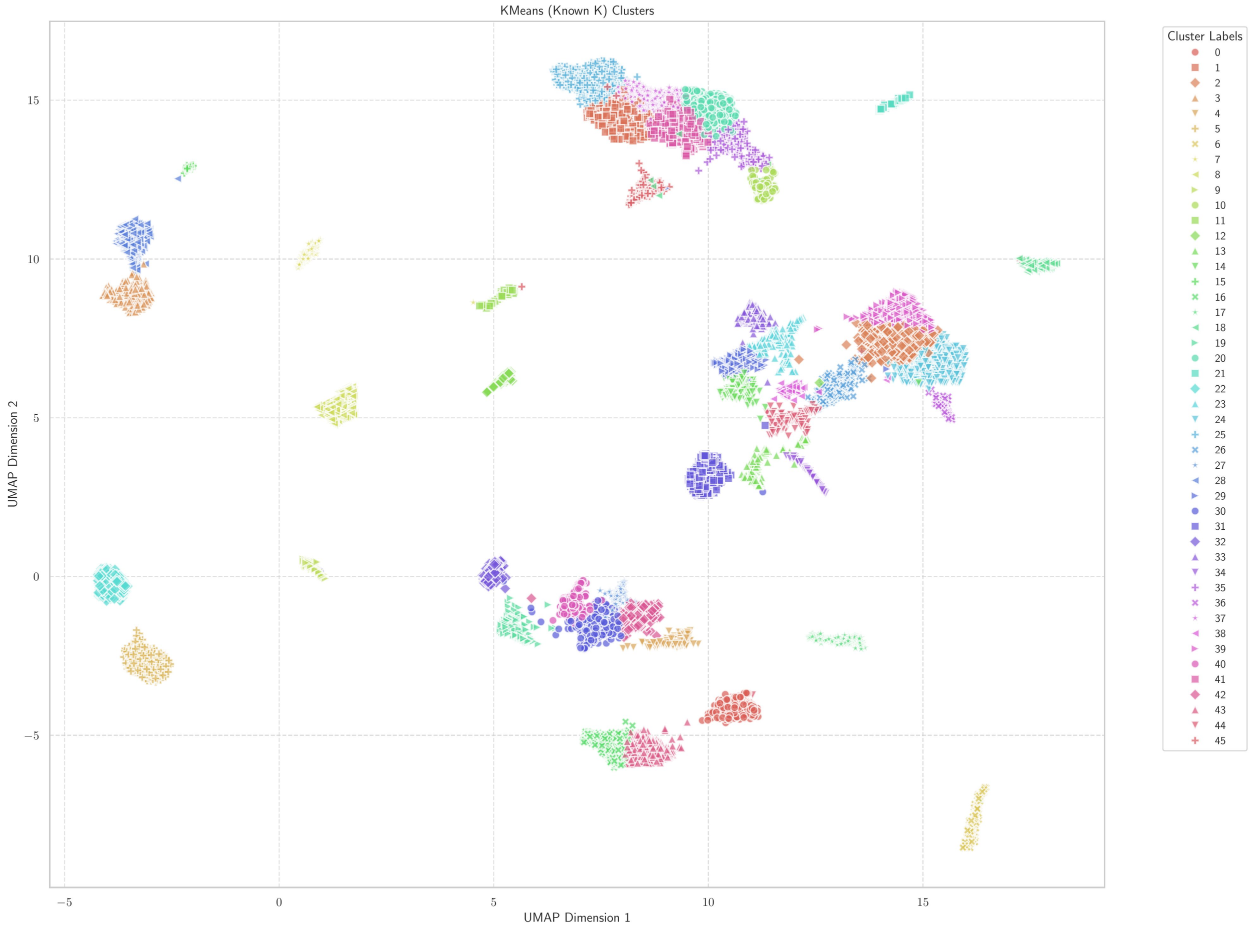

Figure S17: Pollen - (known k) k-Means

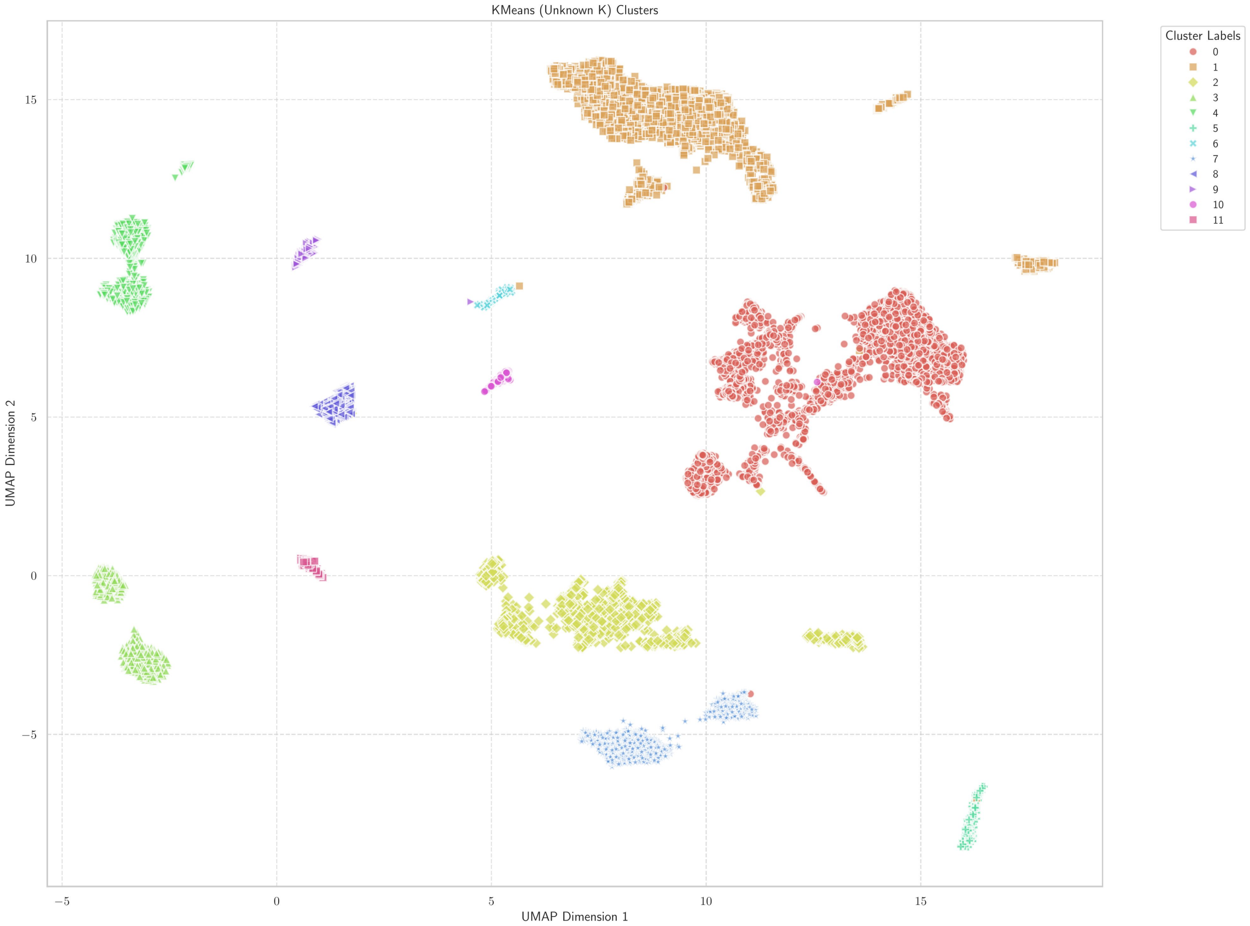

Figure S18: Pollen - (unknown k) k-Means

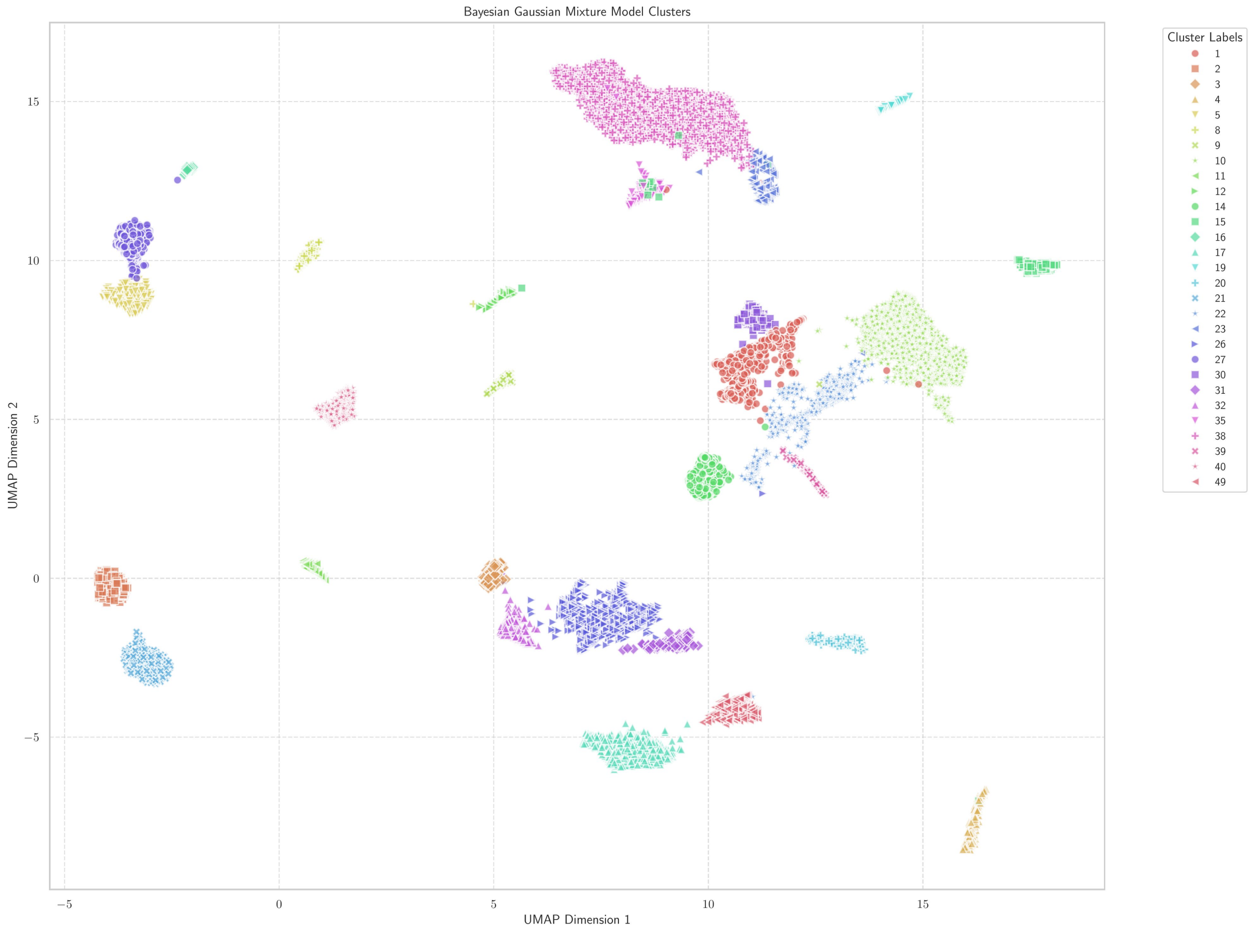

Figure S19: Pollen - BGMM

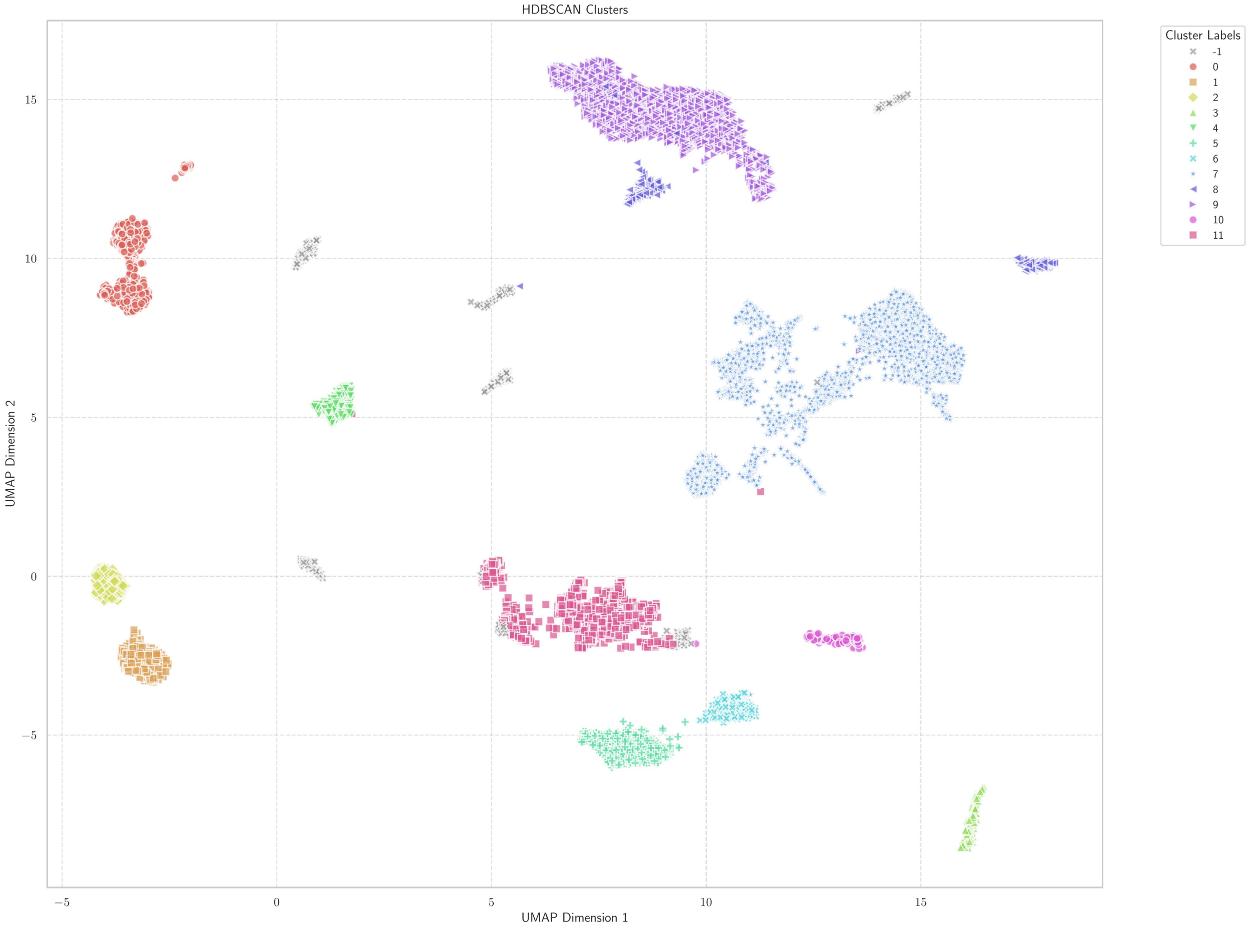

Figure S20: Pollen - HDBSCAN

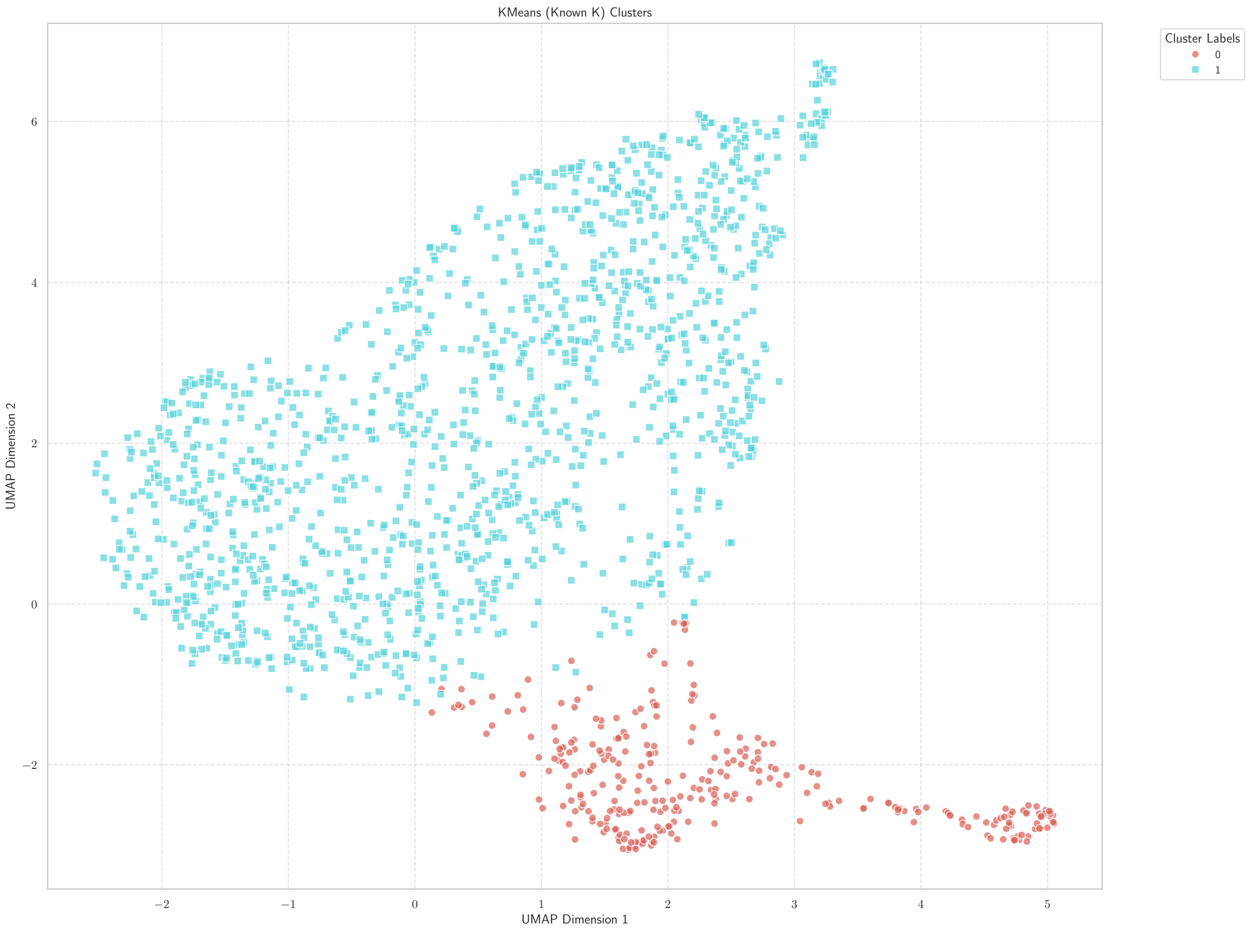

Figure S21: Fossil Tracks - (known k) k-Means

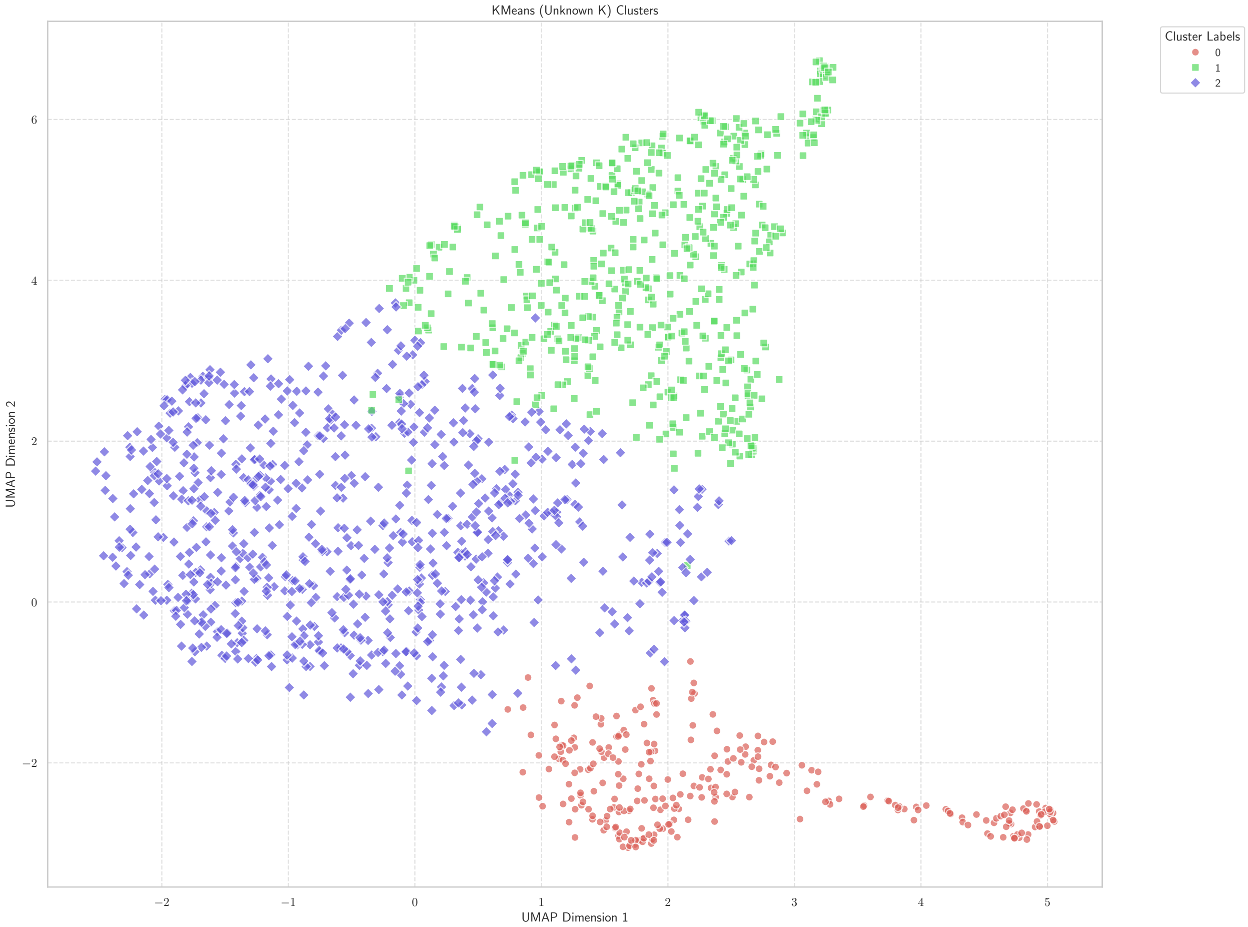

Figure S22: Fossil Tracks - (unknown k) k-Means

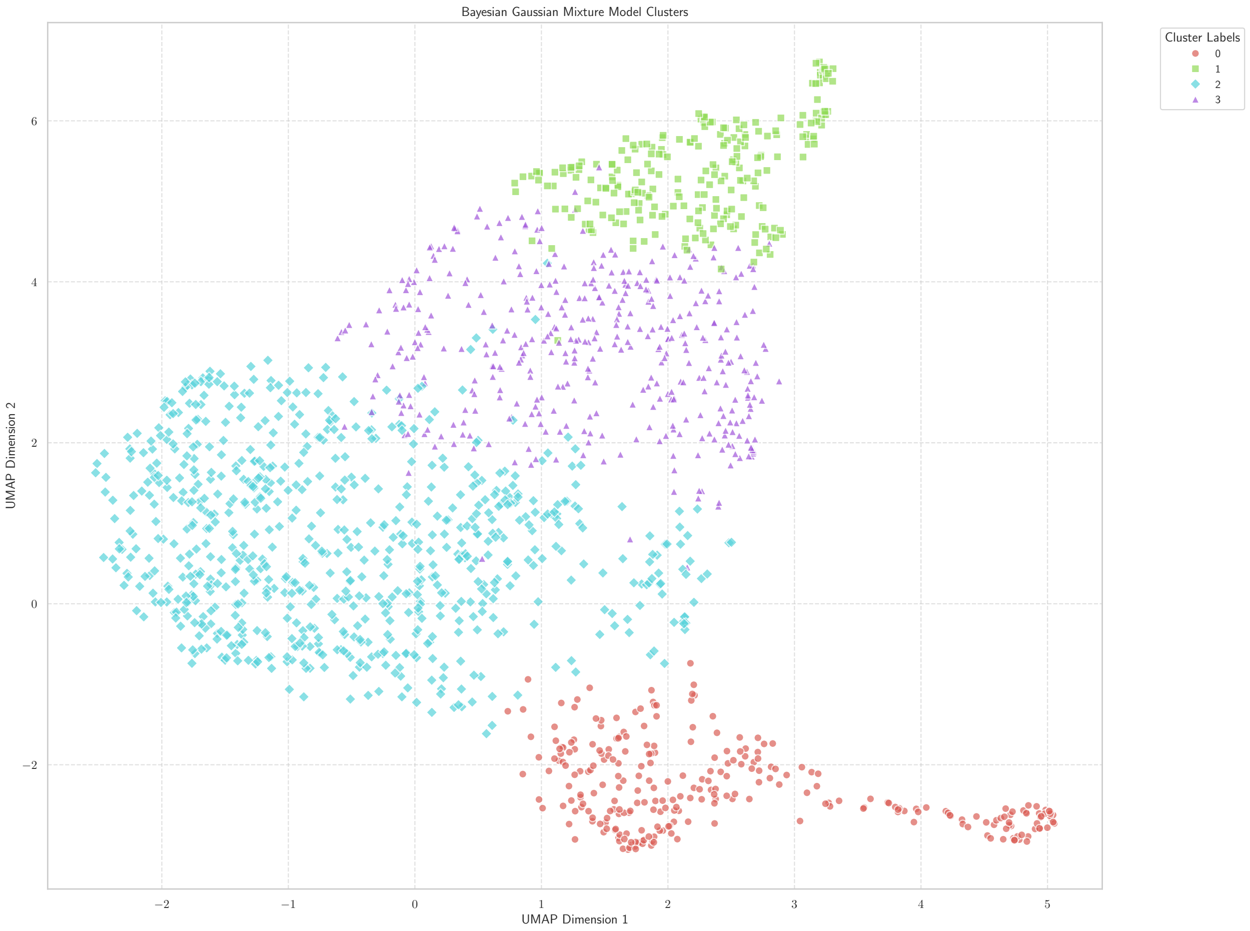

Figure S23: Fossil Tracks - BGMM

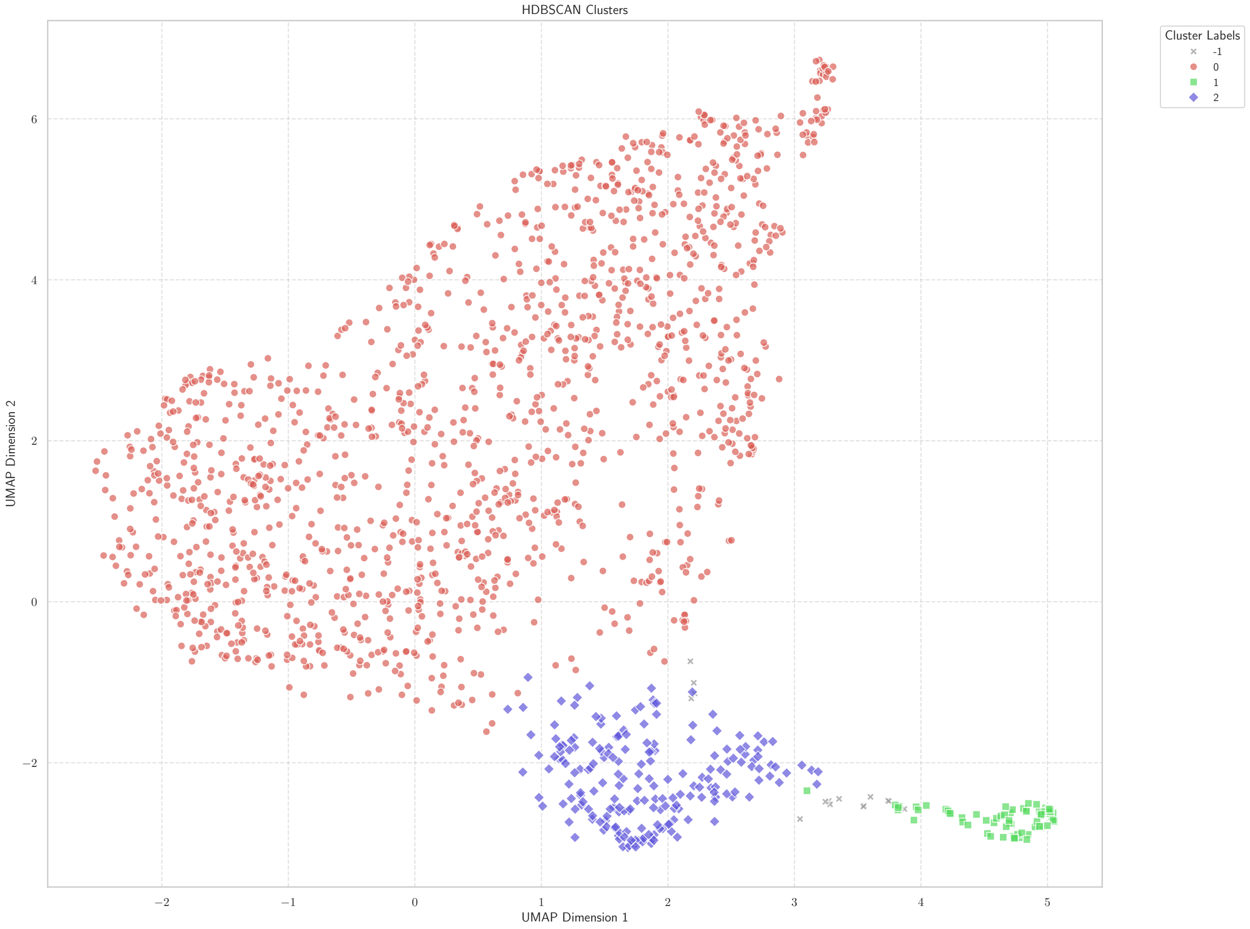

Figure S24: Fossil Tracks - HDBSCAN

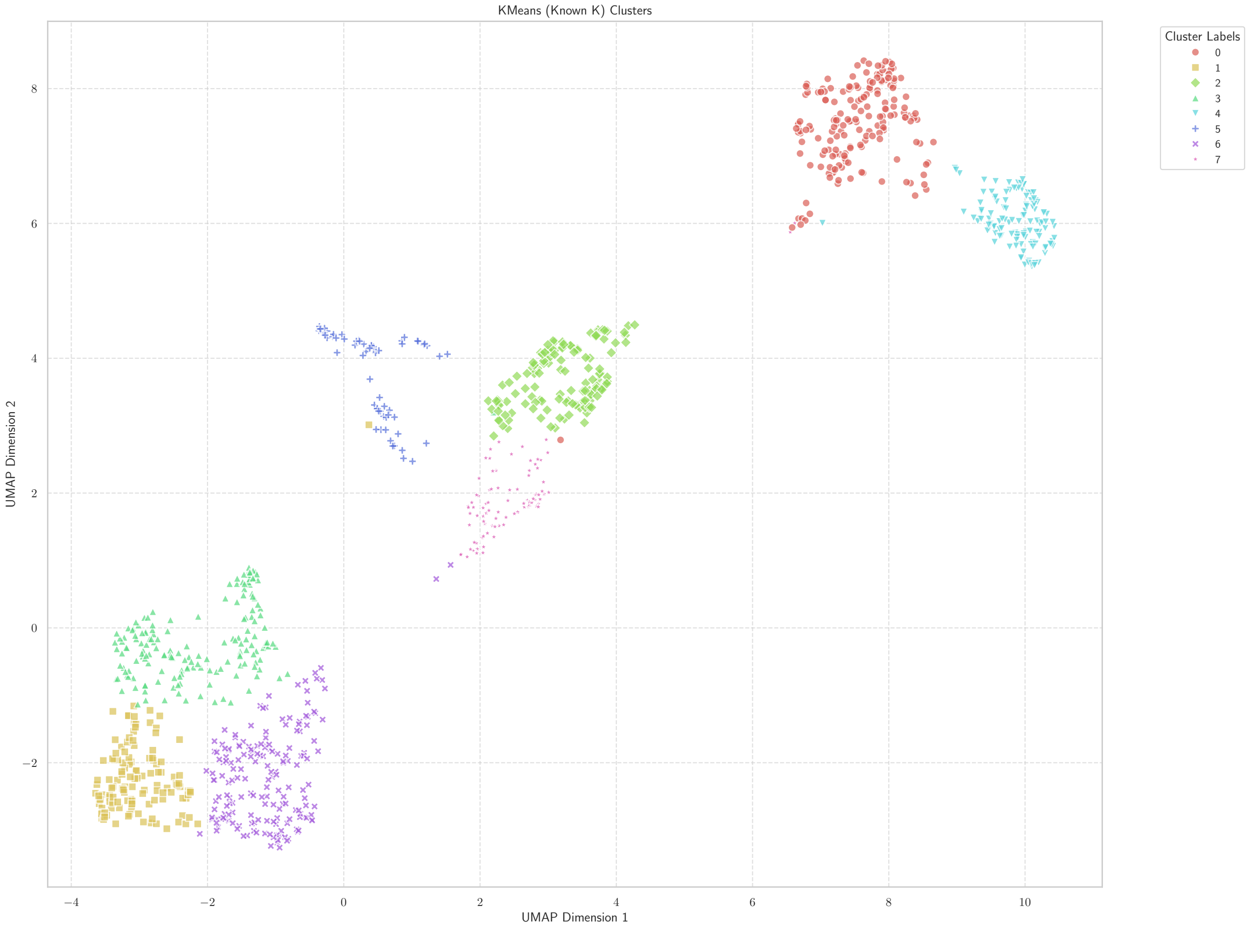

Figure S25: Radiolaria - (known k) k-Means

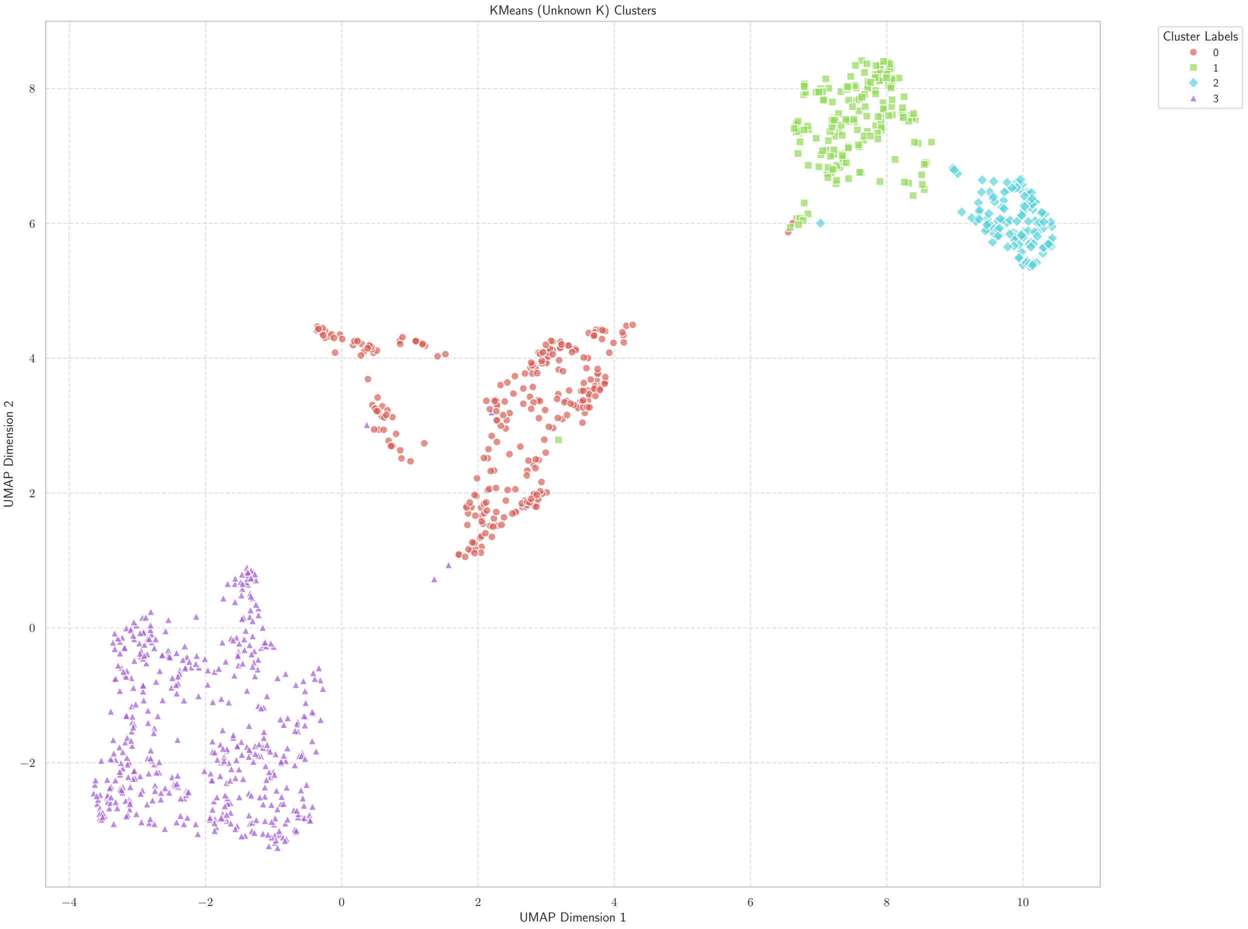

Figure S26: Radiolaria - (unknown k) k-Means

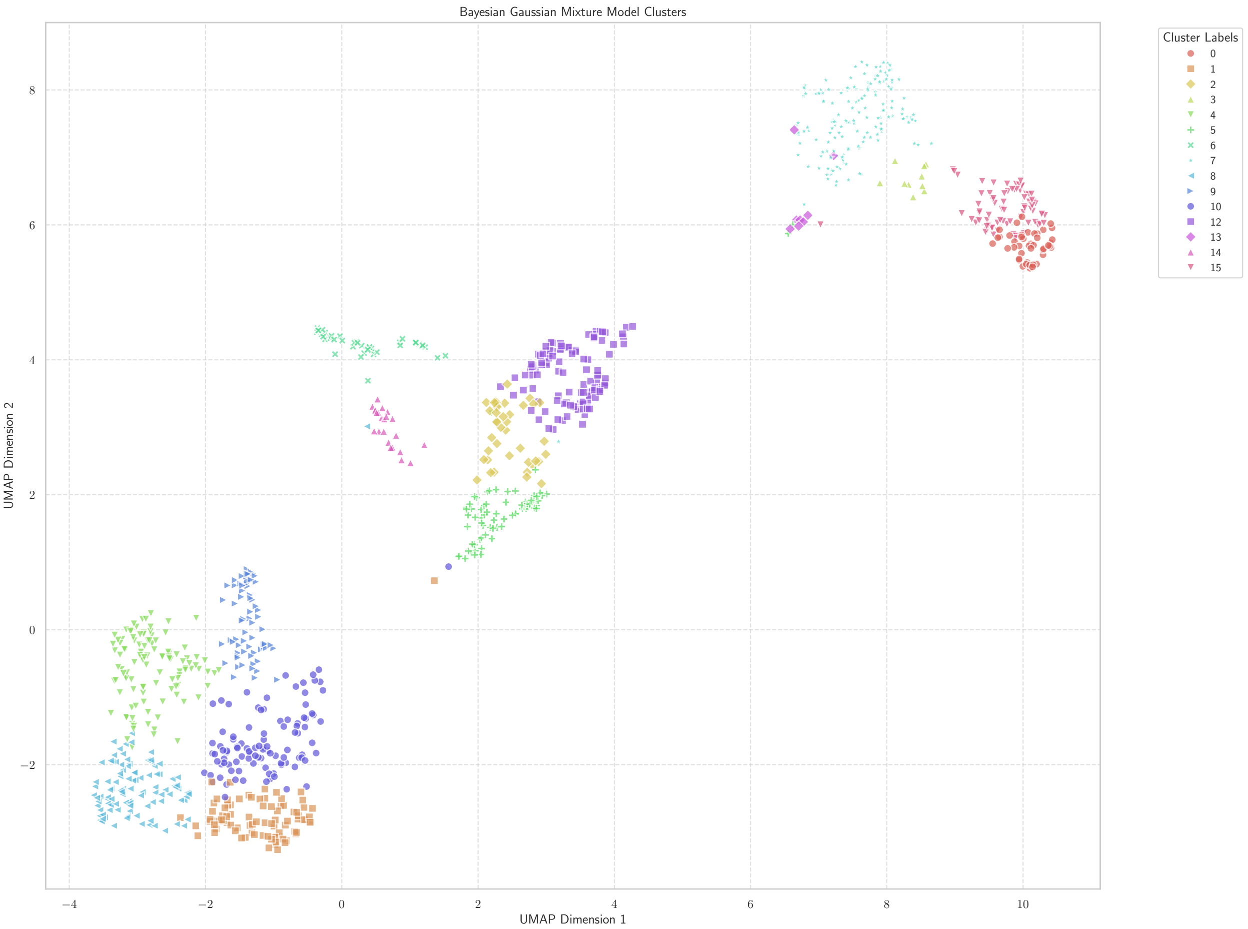

Figure S27: Radiolaria - BGMM

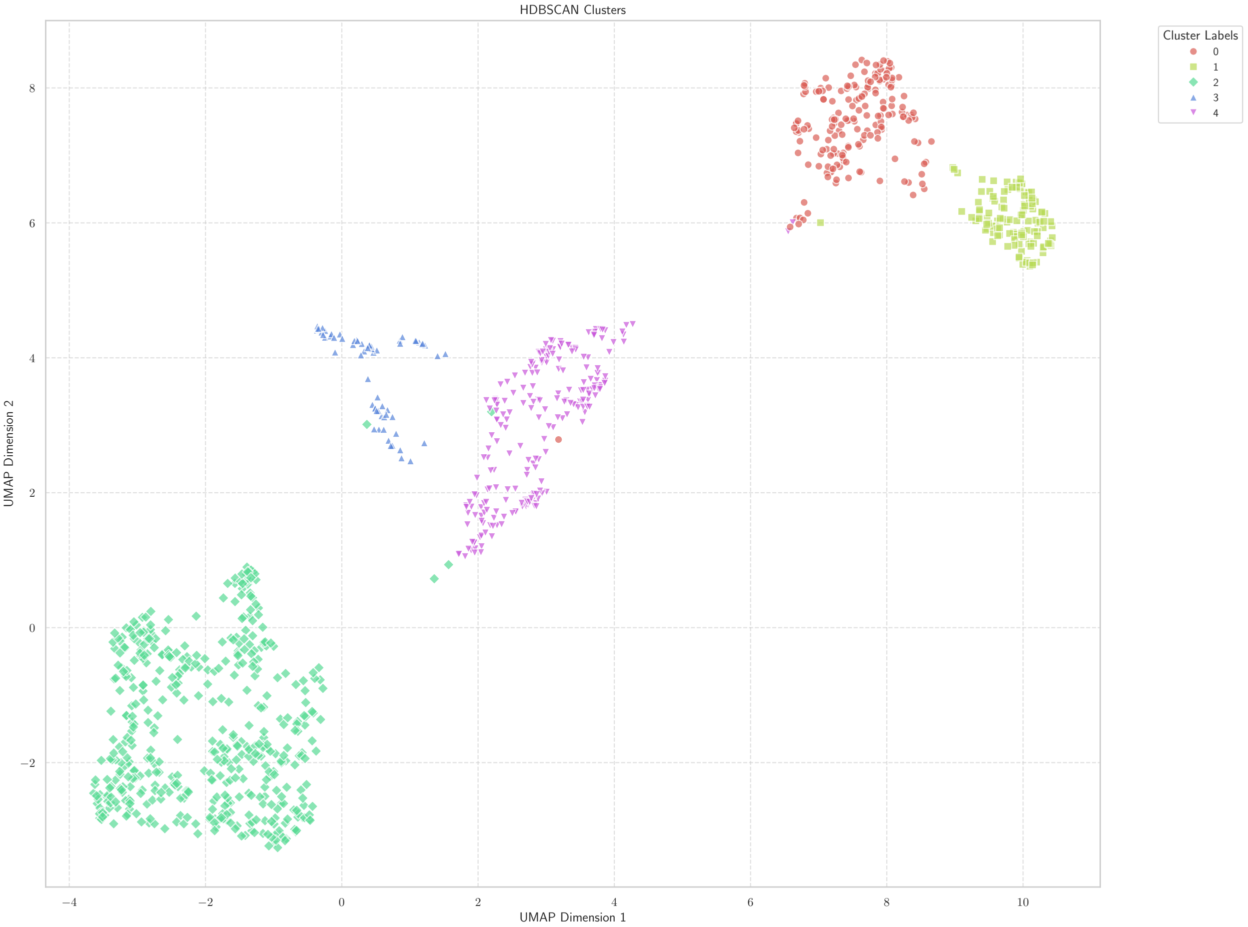

Figure S28: Radiolaria - HDBSCAN

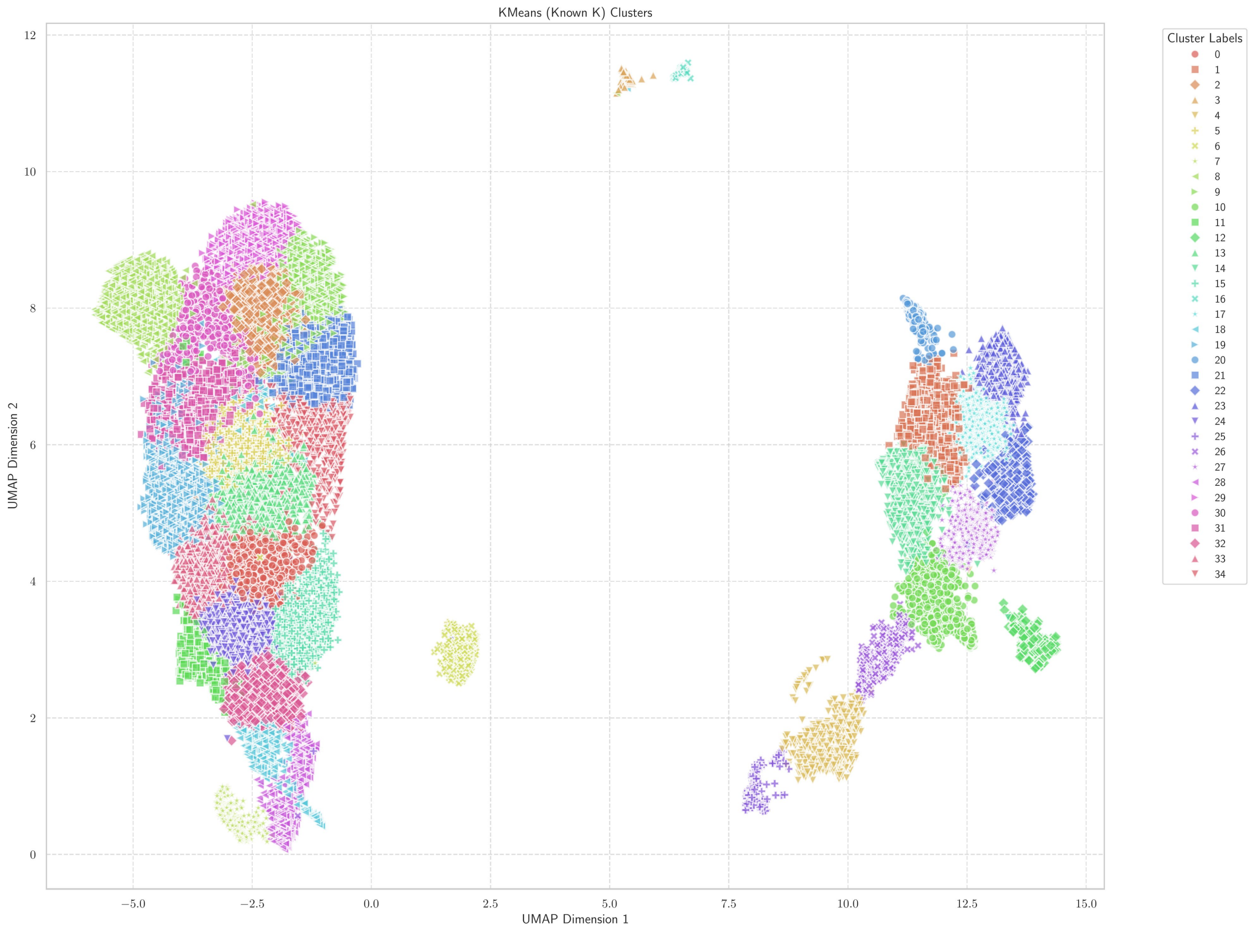

Figure S29: Foraminifera - (known k) k-Means

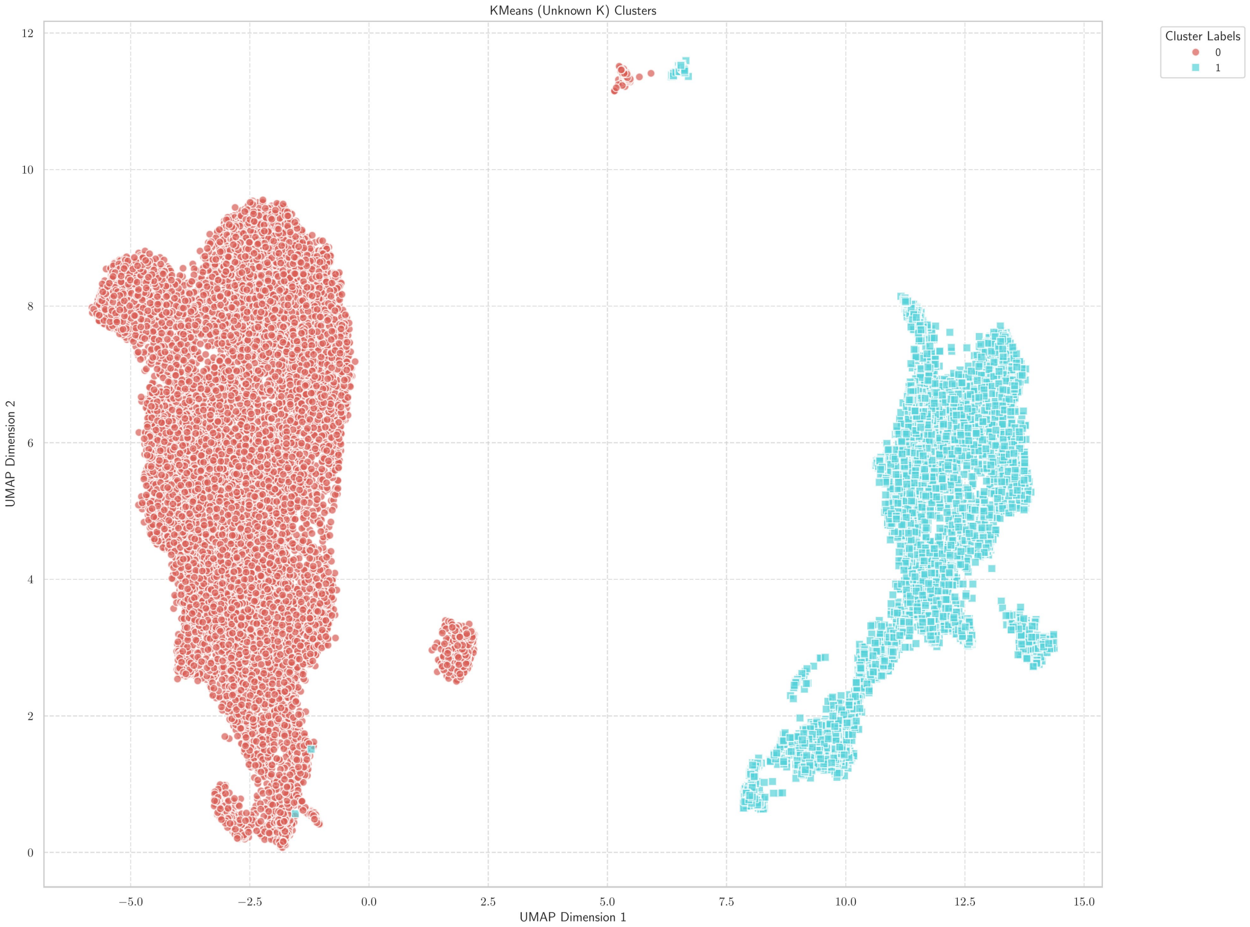

Figure S30: Foraminifera - (unknown k) k-Means

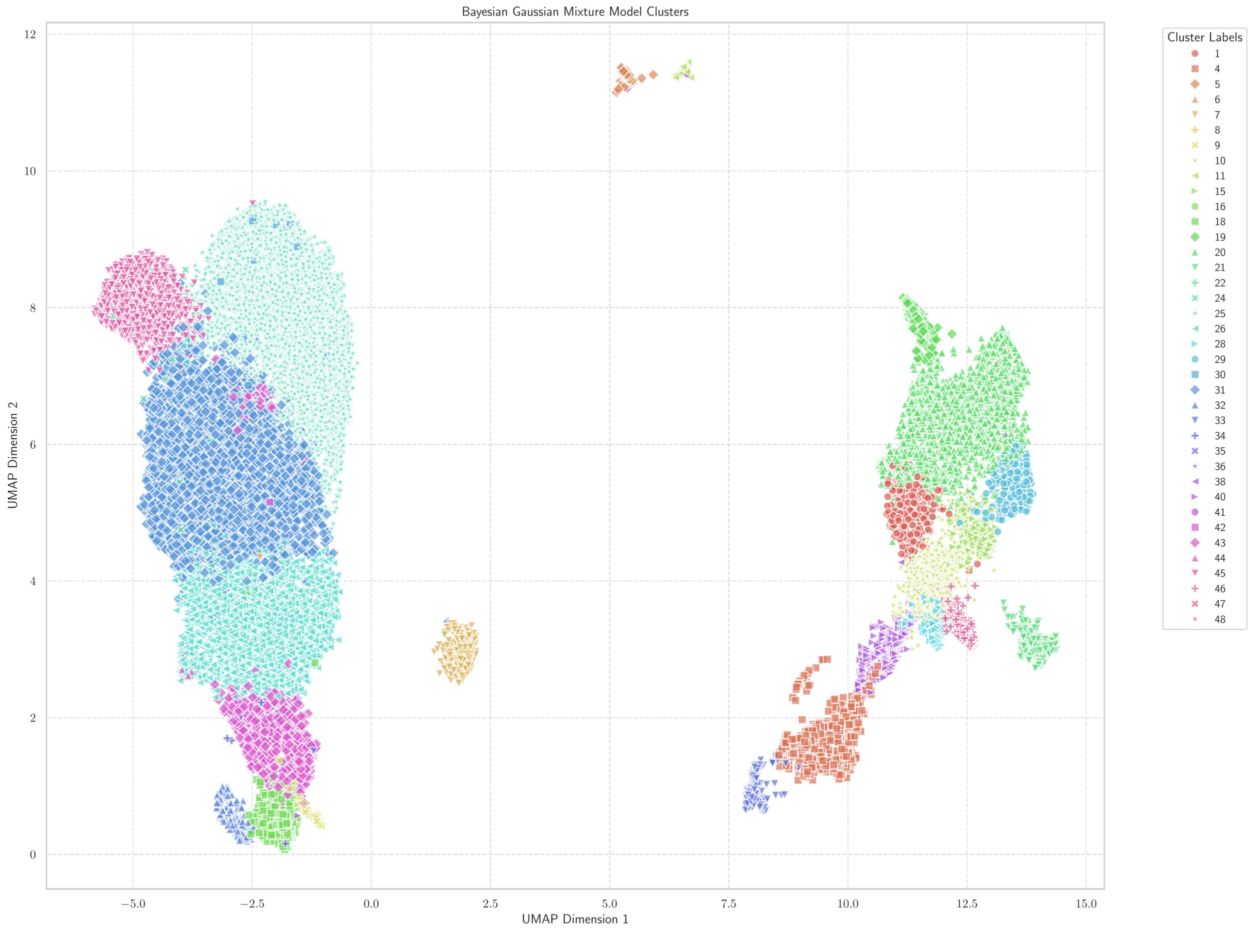

Figure S31: Foraminifera - BGMM

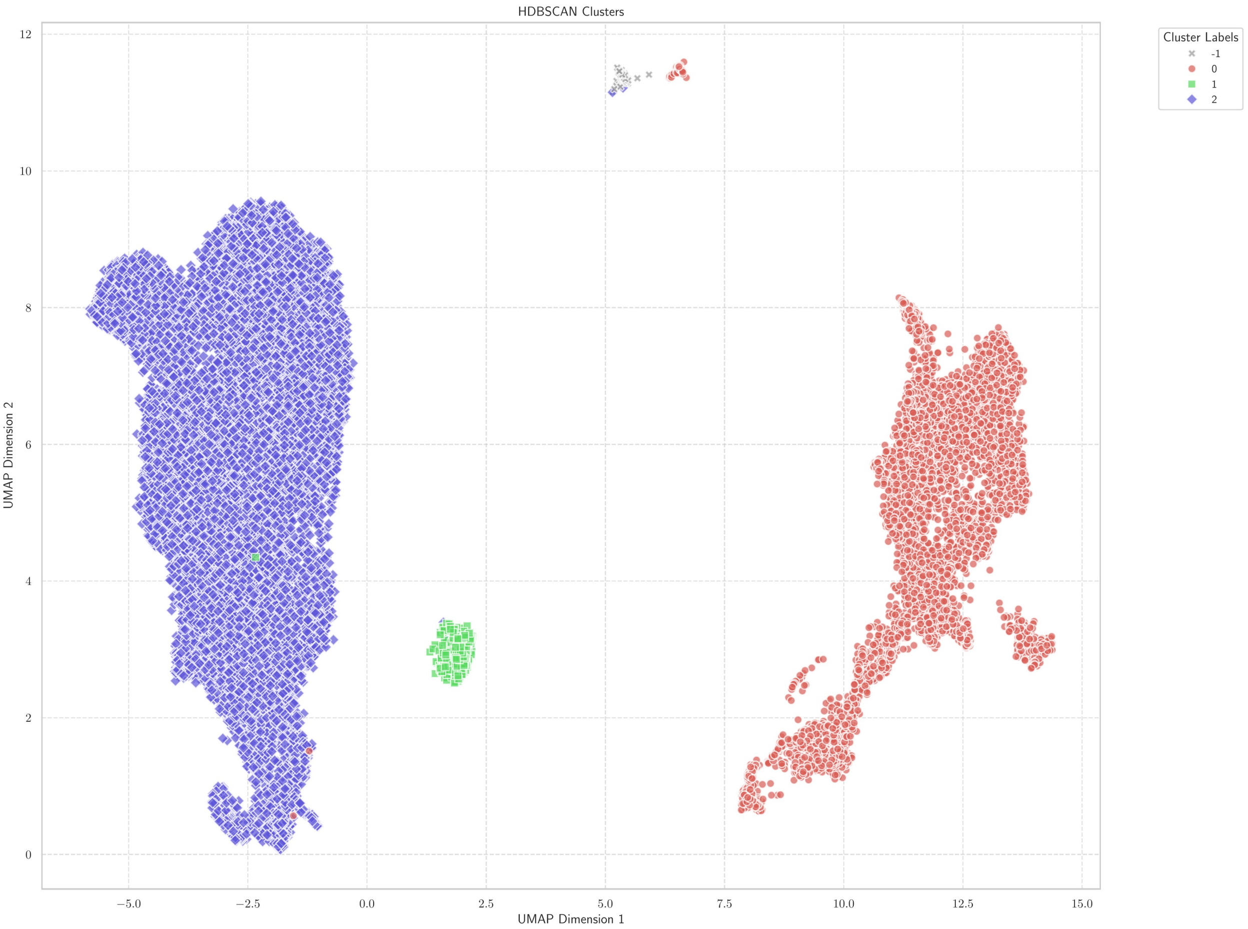

Figure S32: Foraminifera - HDBSCAN

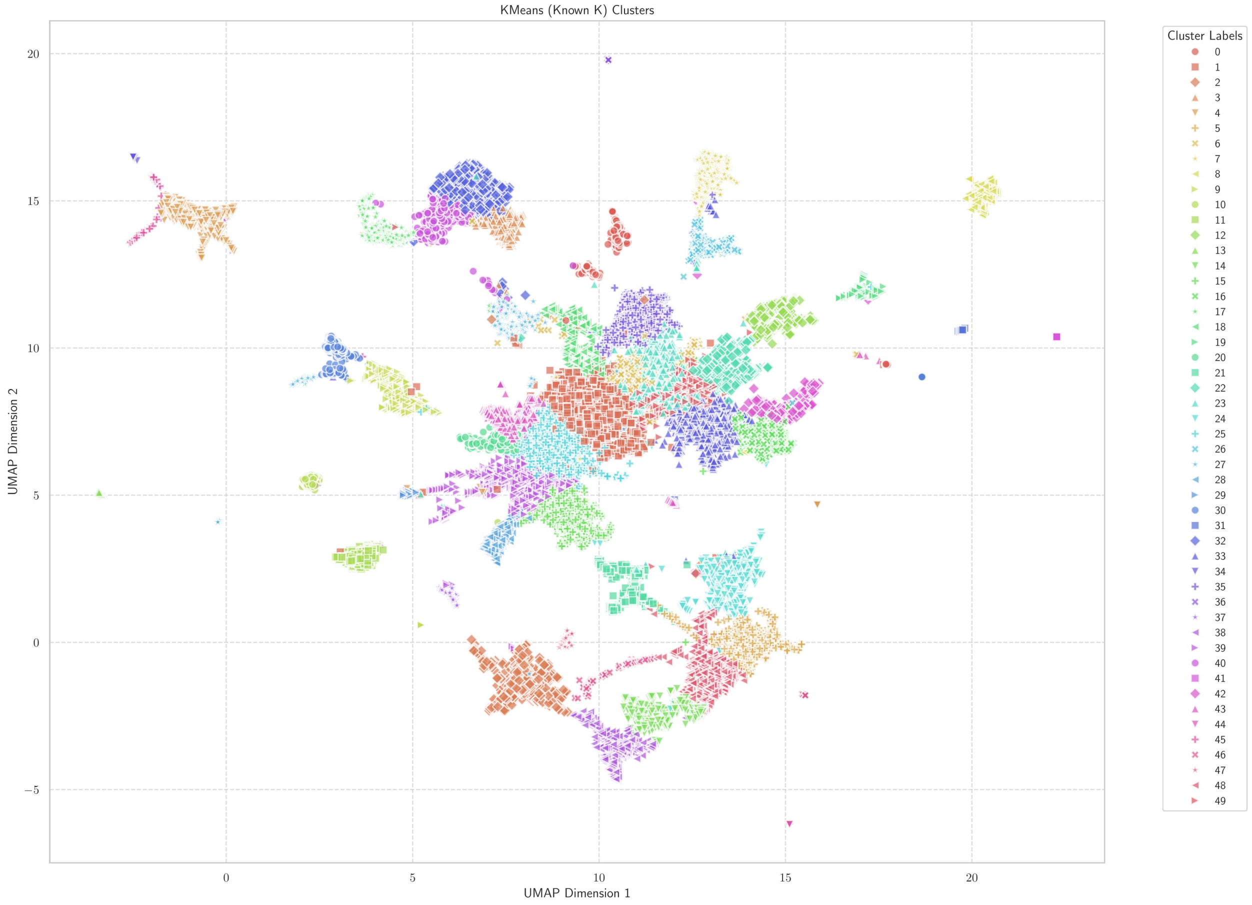

Figure S33: RFID - (known k) k-Means

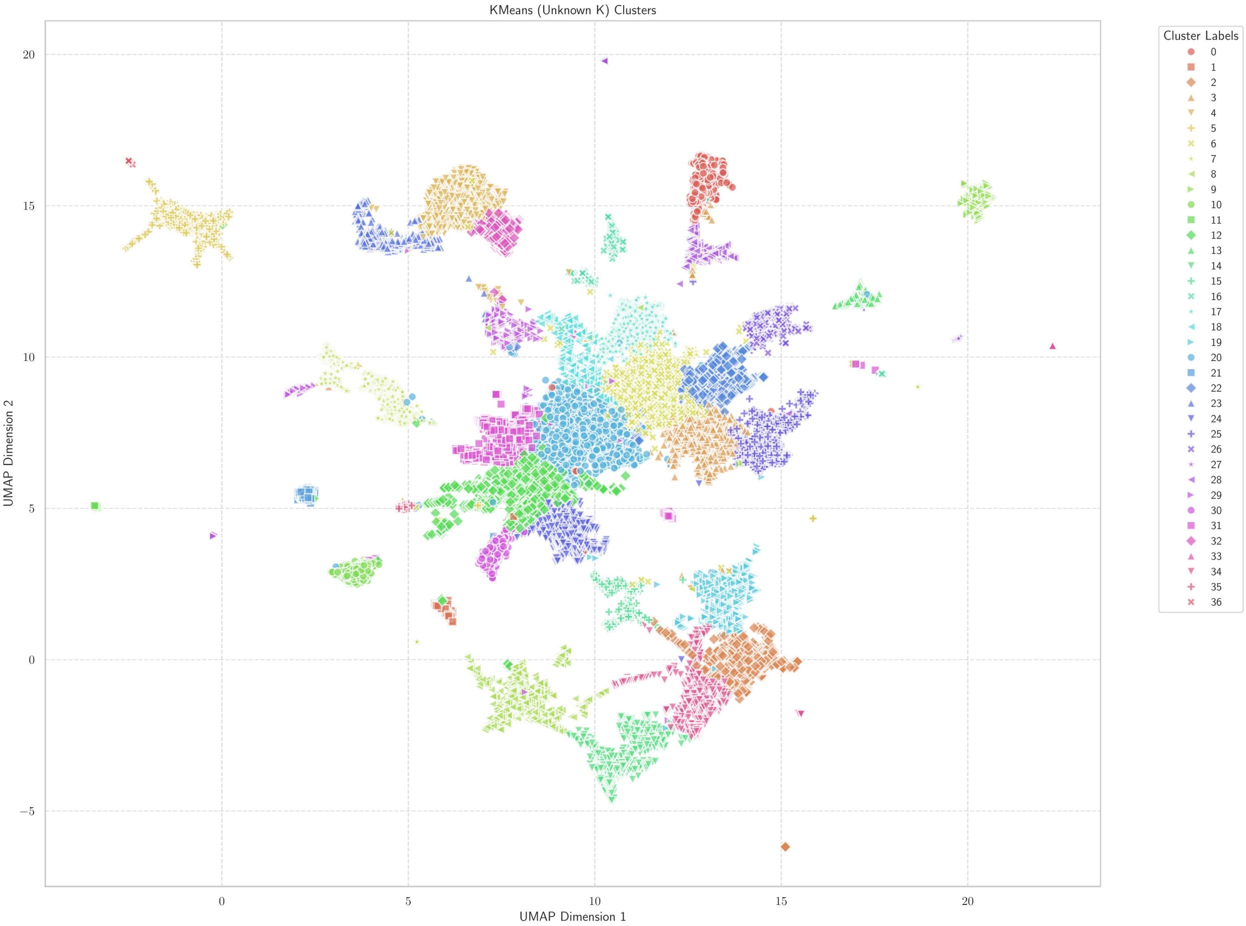

Figure S34: RFID - (unknown k) k-Means

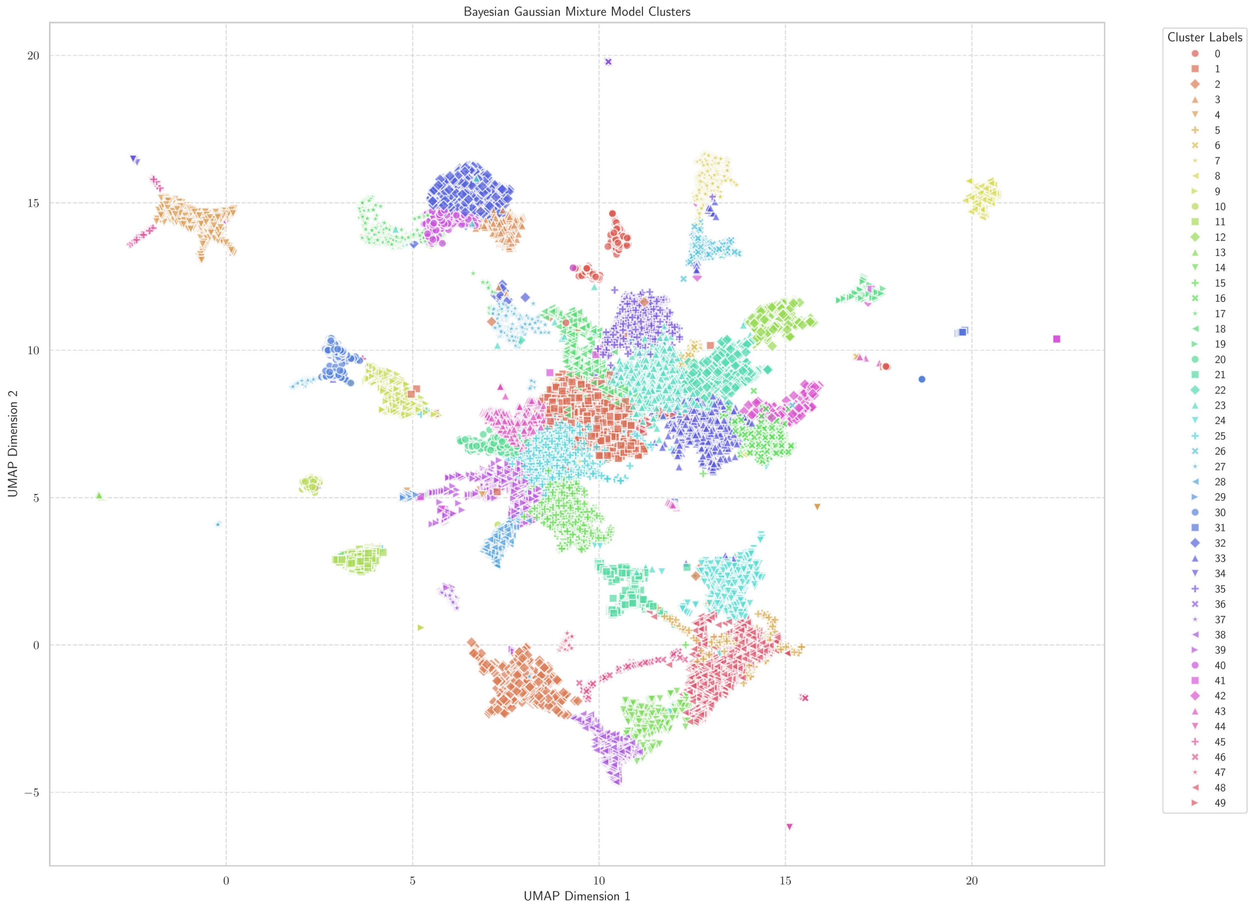

Figure S35: RFID - BGMM

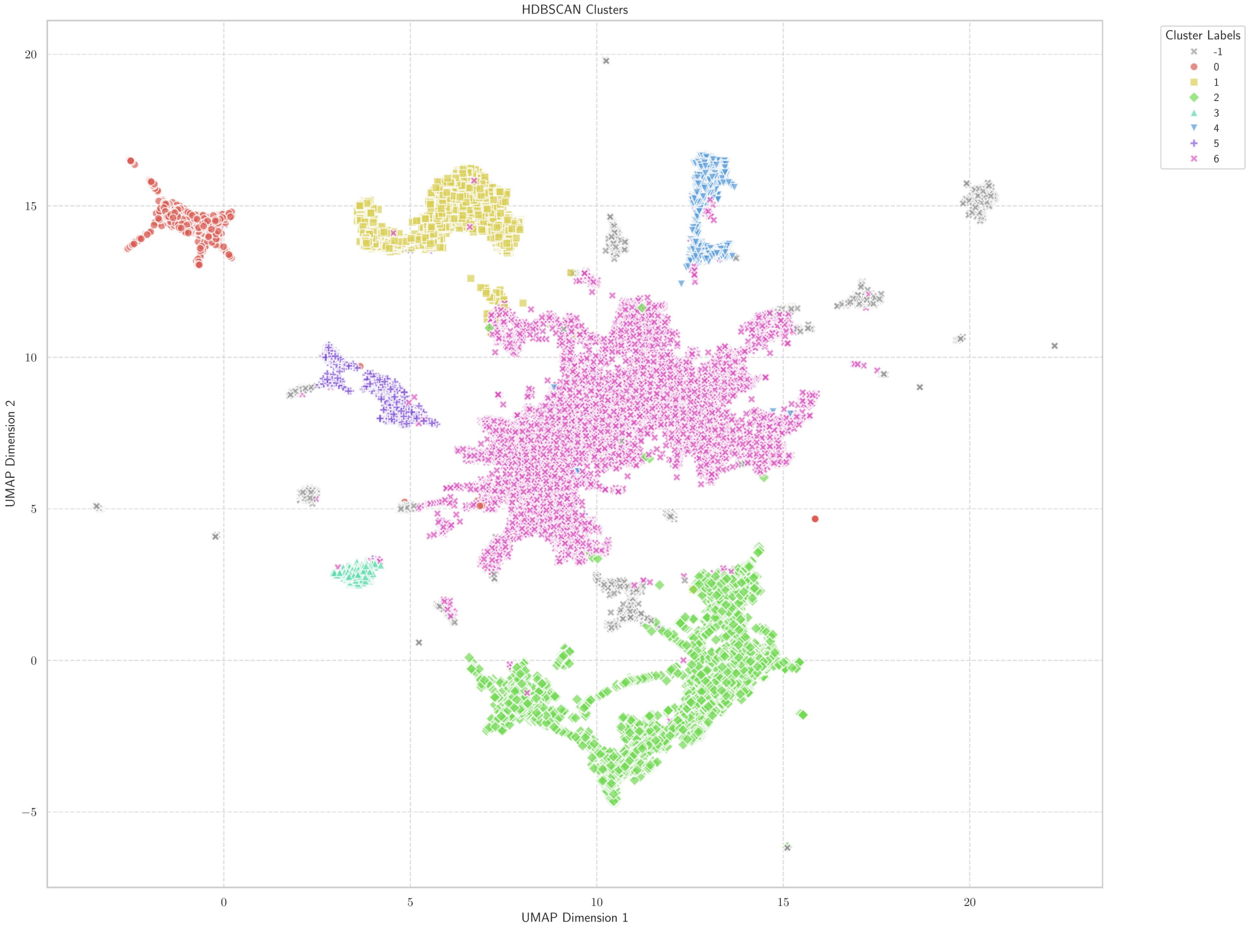

Figure S36: RFID - HDBSCAN
